# Raptor genomes reveal evolutionary signatures of predatory and nocturnal lifestyles

**DOI:** 10.1101/598821

**Authors:** Yun Sung Cho, JeHoon Jun, Jung A Kim, Hak-Min Kim, Oksung Chung, Seung-Gu Kang, Jin-Young Park, Hwa-Jung Kim, Sunghyun Kim, Hee-Jong Kim, Jin-ho Jang, Ki-Jeong Na, Jeongho Kim, Seung Gu Park, Hwang-Yeol Lee, Andrea Manica, David P. Mindell, Jérôme Fuchs, Jeremy S. Edwards, Jessica A. Weber, Christopher C. Witt, Joo-Hong Yeo, Soonok Kim, Jong Bhak

## Abstract

**Background:** Birds of prey (raptors) are dominant apex predators in terrestrial communities, with hawks (Accipitriformes) and falcons (Falconiformes) hunting by day, and owls (Strigiformes) hunting by night.

**Results:** Here, we report new genomes and transcriptomes for 20 species of birds, including 16 species of birds of prey, and high-quality reference genomes for the Eurasian eagle-owl (*Bubo bubo*), oriental scops-owl (*Otus sunia*), eastern buzzard (*Buteo japonicus*), and common kestrel (*Falco tinnunculus*). Our extensive genomic analysis and comparisons with non-raptor genomes identified common molecular signatures that underpin anatomical structure and sensory, muscle, circulatory, and respiratory systems related to a predatory lifestyle. Compared with diurnal birds, owls exhibit striking adaptations to the nocturnal environment, including functional trade-offs in the sensory systems (e.g., loss of color vision genes and selection for enhancement of nocturnal vision and other sensory systems) that are probably convergent with other nocturnal avian orders. Additionally, we found that a suite of genes associated with vision and circadian rhythm were differentially expressed between nocturnal and diurnal raptors, indicating adaptive expression change during the transition to nocturnality.

**Conclusions:** Overall, raptor genomes showed genomic signatures associated with the origin and maintenance of several specialized physiological and morphological features essential to be apex predators.

## Background

Birds of prey, also known as raptors, are key apex predators in nearly every terrestrial biotic community. Species in this guild comprise a non-monophyletic set of three orders within the core landbirds clade, and recent large-scale phylogenomic studies have led to the suggestion that the common ancestor of this clade may have been a predator [1]. There are three main orders of birds of prey: Strigiformes (true and barn owls), Falconiformes (falcons and caracaras), and Accipitriformes (eagles, buzzards, hawks, kites, and vultures). Species in each of these three raptor clades are obligate predators with adaptations for hunting, killing, and/or eating meat [2, 3]. Additionally, the common ancestor of owls evolved nocturnality, and most extant owl species are nocturnal, a habit they share with two other avian orders for which we have genome sequences (Caprimulgiformes and Apterygiformes). These independent transitions in lifestyle provide an opportunity to test for patterns of genome evolution that are linked with being raptorial and nocturnal, respectively [3-5].

Genomes have been published for more than 50 avian species, including nine birds of prey (peregrine and saker falcons, bald, white-tailed, and golden eagles, turkey vulture, barn owl, northern spotted owl, and burrowing owl) [3, 6-9]. However, the barn owl, white-tailed eagle, and turkey vulture genomes were assembled at low-quality [6], and a detailed comparative evolutionary analysis was performed only for the falcons [3]. Here, we report new high-quality whole genome reference sequences of four raptor species (Eurasian eagle-owl [*Bubo bubo*] and oriental scops-owl [*Otus sunia*] in Strigiformes, eastern buzzard [*Buteo japonicus*] in Accipitriformes, and common kestrel [*Falco tinnunculus*] in Falconiformes) with a set of raptor whole-genome and transcriptome data, extending the genomic coverage of raptors (Fig. 1, Additional file 1: Figure S1 and Tables S1-S3). Our investigation revealed numerous genomic signatures of evolution that are shared among the three raptor orders or that appear to be associated with nocturnal adaptations of owls.

**Fig. 1.**
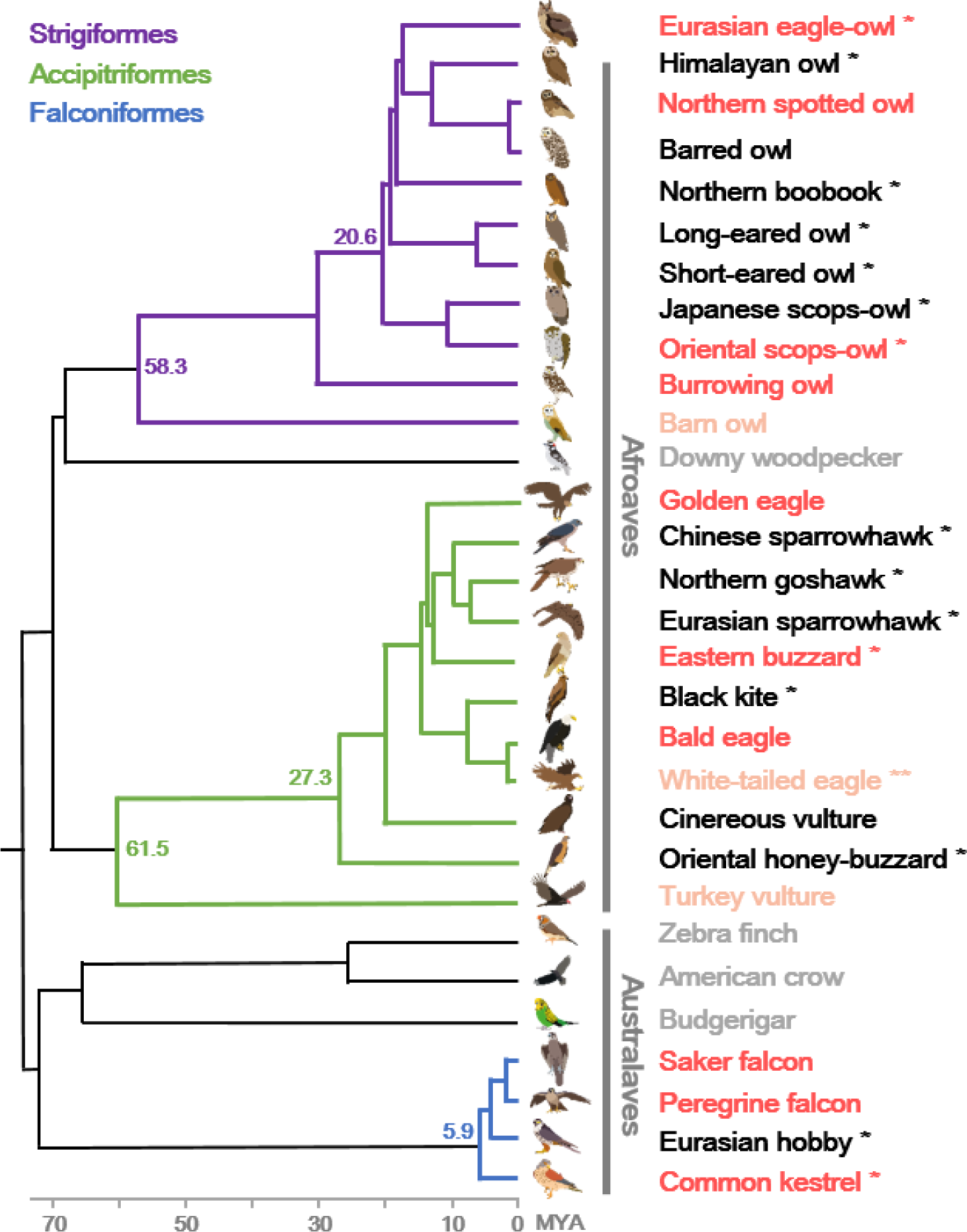
Phylogeny and genomic data of birds of prey. The phylogenetic tree topology was adapted from the Avian Phylogenomics Project [1] and TimeTree database. The estimated divergence time from present (million years ago; MYA) is given at the nodes. Dark red indicates species with higher-quality (scaffold N50 length > 1 Mb) genome assemblies, light red indicates species with lower-quality genome assemblies, black indicates species for which the whole genome was resequenced, and grey indicates non-raptor species high-quality genome assemblies. * denotes birds of prey sequenced from this study. The white-tailed eagle (denoted with **) was previously assembled at low-quality and also whole genome resequenced from this study.

## Results and discussion

### Raptor genome sequencing and assembly

We applied whole-genome shotgun sequencing and *de novo* assembly strategies [10-12] to build reference genomes of the four raptor species (Eurasian eagle-owl, oriental scops-owl, eastern buzzard, and common kestrel). The extracted DNA samples from wild individuals were sequenced at high coverage (>185×) using various insert sizes of short-insert and long-mate pair libraries (Additional file 1: Tables S4 and S5). The four raptor genomes showed relatively higher levels of genomic diversity compared to the previously assembled genomes of eagles and falcons (Additional file 1: Figures S2 and S3). By assembling in various conditions and evaluating assembly quality, we obtained the four raptor reference genomes at a high-quality, resulting in scaffold N50 sizes from 7.49 to 29.92 Mb (Additional file 1: Tables S6-S9). Protein-coding genes (∼16,000 to 18,000 genes) for these four species were predicted by combining *de novo* and homologous gene prediction methods with whole blood transcriptome data (Additional file 1: Table S10). Roughly 9.2% of the raptor genomes were predicted as transposable elements (Additional file 1: Table S11), consistent with the composition of other avian genomes [6]. Additionally, we sequenced the whole genome and blood transcriptome from another twelve raptors (five owls, six accipitrids, and a falconid) and four non-raptor birds (Additional file 1: Tables S12-S14), most of which were sequenced for the first time.

### Evolutionary analysis of raptors compared to non-raptor birds

To identify the genetic basis of predation and nocturnality in raptors, we performed in-depth comparative evolutionary analyses for 25 birds of prey (including ten nocturnal owls and 15 diurnal raptors) and 23 non-raptor bird species (including nocturnal brown kiwi [13] and chuck-will’s-widow [6], and other avian representatives genome-assembled at a high-quality; Fig. 2, Additional file 1: Figure S4 and Tables S1, S2, and S15). Birds have evolved to employ many different strategies to obtain food, and raptors are specialized for hunting [2, 3, 7]. Several molecular signatures were shared by the three raptor orders, and the ancestral branches of these orders each showed an expansion of gene families associated with regulation of anatomical structure size, embryonic appendage morphogenesis, regulation of responses to stimulus and wounding, and learning or memory functions (*P* <0.05, Fisher’s exact test; Additional file 1: Tables S16 and S17). When comparing gene family sizes between the extant species, immune system associated gene families were expanded in the birds of prey (Additional file 1: Table S18).

**Fig. 2.**
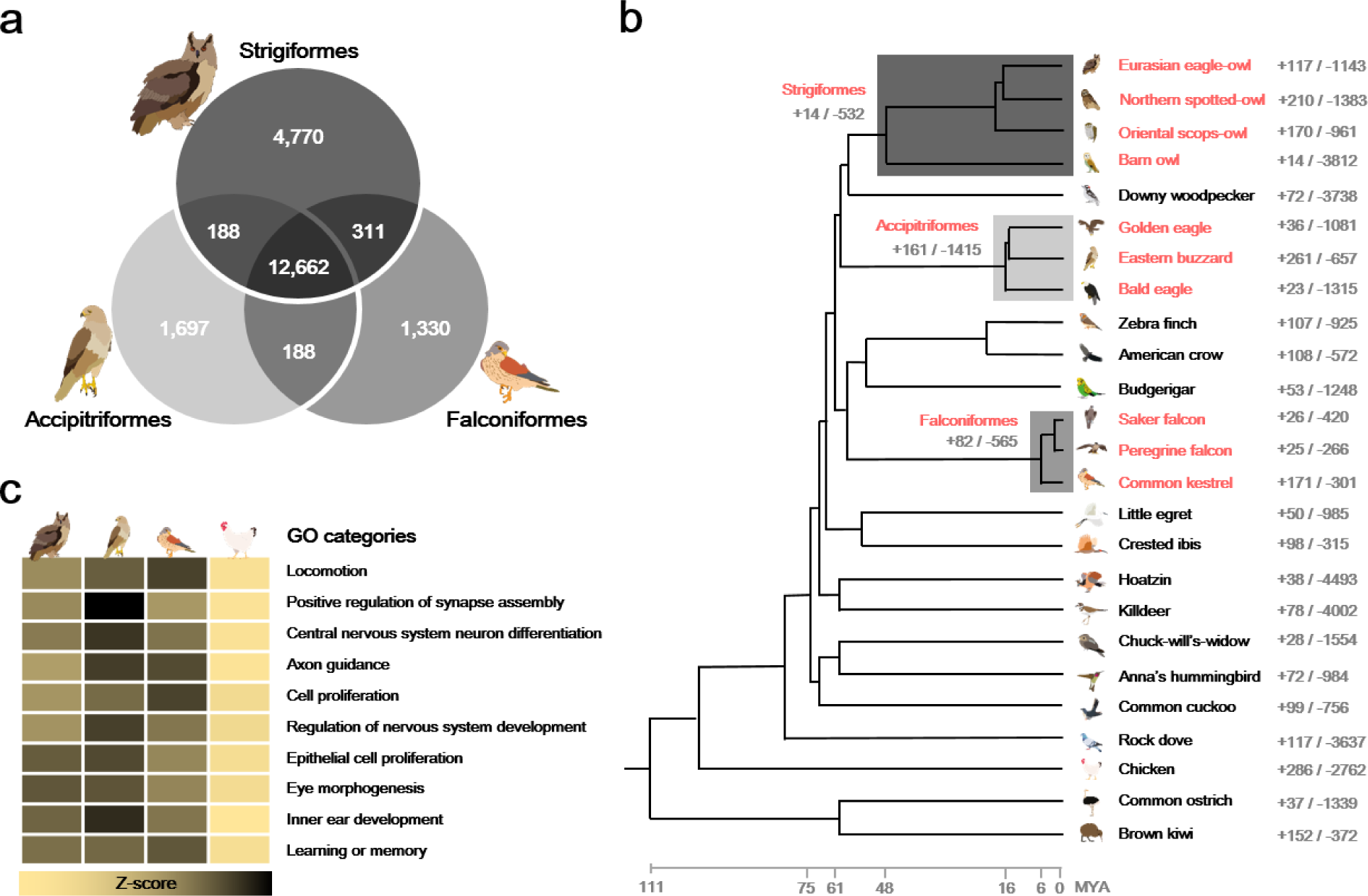
Relationship of birds of prey to other avian species. **a** Venn diagrams of orthologous gene clusters in the birds of prey. Orthologous gene clusters were constructed using 25 avian genomes. Only raptor gene clusters are displayed. **b** Gene expansion or contraction in avian species. The numbers near order and species names indicate the number of gene families that have expanded (+) and contracted (-) in each branch and species. Species in red are birds of prey. **c** Heatmap of enriched Gene Ontology (GO) categories for raptor common GC3 biased genes. Bird icons from left to right indicate Strigiformes, Accipitriformes, Falconiformes, and non-raptor birds. Z-scores for the average of normalized GC3 percentages are shown as a yellow-to-black color scale.

To further examine the shared evolutionary adaptations related to avian predatory lifestyles, we identified selection signatures shared by the three orders of birds of prey at the gene sequence level; which possibly reflects their shared requirement for highly-developed sensory systems, efficient circulatory and respiratory systems, and exceptional flight capabilities necessary to capture prey [2-5, 7, 8]. Based upon *d*_*N*_/*d*_*S*_ ratio calculation [14, 15], only *RHCE* and *CENPQ* genes were commonly found as positively selected genes (PSGs) in the three raptor ancestral branches of the Strigiformes, Accipitriformes, and Falconiformes (Additional file 2: Datasheets S1-S3); consistent with the results from mammals demonstrating that adaptive molecular convergence linked to phenotypic convergence is rare [12, 16]. In addition, we identified three genes as positively selected in the ancestral branches of two raptor orders (*SFTPA1* in the Strigiformes and Falconiformes; *TFF2* and *PARL* in the Strigiformes and Accipitriformes). A lung surfactant protein encoded by *SFTPA1* paly an essential role in the defense against respiratory pathogens and normal respiration [17]. *TFF2* gene encodes a protein that mediate gastric wound repair and inhibit gastric acid secretion [18]. Finally, we found that 148 genes showed accelerated *d*_*N*_/*d*_*S*_ in the raptor ancestral branches (Additional file 1: Table S19). Of these, *SLC24A1, NDUFS3*, and *PPARA* encode proteins that play roles in visual transduction cascade, mitochondrial membrane respiratory chain, and lipid metabolism, respectively [17, 19, 20]. Out of 50 collected beak development associated genes, 17, 17, and 18 genes (34 to 36%) showed accelerated *d*_*N*_/*d*_*S*_ in the ancestral branches of Strigiformes, Accipitriformes, and Falconiformes, respectively (Additional file 1: Table S20), hinting at adaptation for enhanced beaks for killing and flesh-tearing [2, 3]. Of these, four genes (*BMP10, GDF9, NAB1*, and *TRIP11*) showed common acceleration signatures in the three raptor orders.

It has been suggested that genes with elevated frequencies of Guanine-Cytosine at the third codon position (GC3) are more adaptable to external stresses, through providing more targets for *de novo* methylation that affect the variability of gene expression [23]. Therefore, we analyzed the GC3 content in the three raptor orders, and we found that regulation of nervous system development, central nervous system neuron differentiation, and locomotion associated genes showed high GC3 bias (Fig. 2c, Additional file 1: Figure S5, Table S21 and Additional file 2: Datasheet S6). In the highly conserved genomic regions (HCRs) among species belonging to the same order [12], 79 functional categories were commonly enriched in the three raptor orders (Additional file 1: Tables S22-S31). Among these categories, eye, sensory organ, muscle organ, epithelium, and limb development functions were commonly conserved in the three raptor orders, but not in Passeriformes (a control avian order in this analysis), suggesting that those functions are important in raptors for their predatory lifestyle.

### Evolutionary analysis of nocturnal birds compared to diurnal birds

Since several avian clades have adapted to a nocturnal lifestyle independently, the comparative method can be used to identify genes underlying convergent phenotypes that are associated with nocturnal adaptation [5]. Three nocturnal bird groups (the ancestral branch of owls, chuck-will’s-widow, and brown kiwi) shared an expansion of gene families associated with synapse organization, cellular response to stimulus, and bile secretion functions (*P* <0.05; Additional file 1: Tables S32, S33). As expected, gene families associated with vision were commonly contracted in the nocturnal birds (Additional file 1: Tables S34 and S35). Specifically, gene loss of the violet/ultraviolet-sensitive opsin *SWS1* (*OPN1SW*) was found in all of the nocturnal bird genomes, as previously reported [4, 22]. The nocturnal birds also showed common selection signatures likely linked to their adaptation to a nocturnal environment. A total 14 PSGs were shared among the three nocturnal groups, and 98 PSGs were shared by at least two nocturnal bird groups (Additional file 2: Datasheets S1, S4 and S5). The shared PSGs were over-represented in detection of mechanical stimulus involved in sensory perception of sound, wound healing, and skin development functions (Additional file 1: Table S36), although the enrichment did not pass the false discovery rate criterion. Interestingly, at least one of two wound healing associated genes (*TFF2* and *COL3A1*) [23, 24] was found to be positively selected in the nocturnal birds. Additionally, six genes (*RHO, BEST1, PDE6B, RPE65, OPN4-1*, and *RRH*) involved in light detection, and *RDH8* that is involved in retinol (vitamin A_1_) metabolism [17, 25], showed accelerated *d*_*N*_/*d*_*S*_ in the nocturnal birds (Additional file 1: Table S37). It is well-known that rhodopsin encoded by *RHO* is a light-sensitive receptor and thus enables vision in low-light conditions [26]. Notably, *RHO* also showed a high level of GC3 biases in the nocturnal birds (Additional file 2: Datasheet S7). Furthermore, *RPE65* encodes a protein that is a component of the vitamin A visual cycle of the retina, while *PDE6B* plays a key role in the phototransduction cascade and mutations in this gene result in congenital stationary night blindness. In addition, melanopsin encoded by *OPN4-1* is a photoreceptor required for regulation of circadian rhythm [17, 25]. We also found that only *SLC51A* gene possesses specific amino acid sequences to the nocturnal birds (Additional file 1: Figure S6). *SLC51A*, also known as *OST-*α, is essential for intestinal bile acid transport [27], and it has been suggested that the bile acids affect the circadian rhythms by regulating the expression level of circadian clock associated gene families [28, 29]. Interestingly, burrowing owl (*Athene cunicularia*), which is known as one of diurnal/crepuscular owls, showed a different sequence alteration pattern from the other nocturnal or diurnal birds in *SLC51A* locus.

### Sensory adaptations to nocturnal environment

Modifications of the major sensory systems (not only vision, but also olfaction, hearing, and circadian rhythm) are among the most common changes that occur when shifting from a diurnal to a nocturnal lifestyle [5]. Analysis of the major sensory systems in the nocturnal bird genomes revealed evidence of highly developed senses for adaptation to nocturnality. First, vision system associated genes showed significantly accelerated *d*_*N*_/*d*_*S*_ in the three nocturnal birds compared to diurnal birds (*P* <0.05, Mann-Whitney *U* test; Fig. 3). Owls and chuck-will’s-widow (Caprimulgiformes) had the highest acceleration in vision-related genes. The total number of functional olfactory receptors (ORs) was not larger in the nocturnal birds than that in the diurnal birds. However, the numbers of γ-clade ORs in the nocturnal birds and γ-c-clade ORs in the owls were significantly larger than others (after excluding two outlier species showing extensive γ-c-clade ORs expansion, chicken and zebra finch; *P* <0.05, Mann-Whitney *U* test; Fig. 3 and Additional file 1: Table S38). The diversity of ORs is thought to be related to a detection range of odors [30], and we found that the diversity of α-clade ORs was significantly higher in the nocturnal birds (Additional file 1: Table S39). Additionally, the diversity in the γ-c-clade ORs was much higher in the owls and brown kiwi (Apterygiformes) compared to their sister groups (downy woodpecker in Piciformes and common ostrich in Struthioniformes, respectively), suggesting that increased olfactory abilities evolved repeatedly under nocturnal conditions [5, 13]. Hearing system associated genes showed a relatively high-level of *d*_*N*_/*d*_*S*_ ratio in the owls and brown kiwi; interestingly, two vocal learning species (budgerigar in Psittaciformes and Anna’s hummingbird in Apodiformes) had the first and third most accelerated *d*_*N*_/*d*_*S*_ for hearing associated genes, which may be linked with their highly developed cognitive abilities [31, 32]. Circadian rhythm associated genes showed the first and second largest acceleration in the owls and brown kiwi, but the lowest in chuck-will’s-widow, suggesting that these independent instances of adaptation to nocturnality occurred by different mechanisms [5]. Additionally, we found that 33 hearing system and 18 circadian rhythm associated genes showed accelerated *d*_*N*_/*d*_*S*_ in the three nocturnal bird groups (Additional file 1: Table S40). Considered together, these results suggest that selection to augment nocturnal vision and other sensory systems predictably compensates for loss of color vision, supporting a functional trade-off of sensory systems in nocturnal birds [4, 5, 13].

**Fig. 3.**
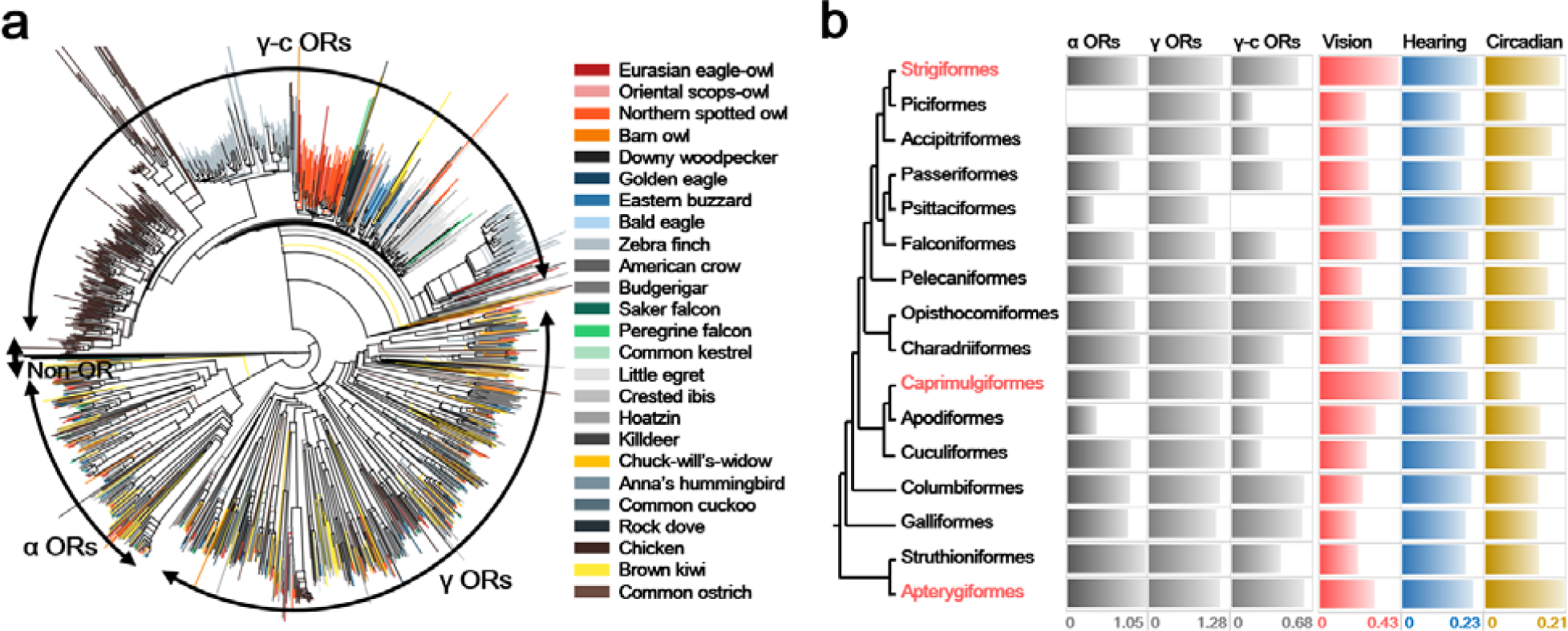
A functional trade-off of sensory systems in nocturnal birds. **a** The phylogeny of the α and γ olfactory receptor (OR) genes identified in 25 avian genomes. Only intact OR genes were used. **b** Selection constraints on sensory systems. Values for α, γ, and γ-c ORs are the diversity of ORs in each clade. For avian orders including two or more genomes (Strigiformes, Accipitriformes, Passeriformes, Falconiformes, and Pelecaniformes), the average diversity values were used. The diversity of α ORs in Piciformes and γ-c ORs in Psittaciformes were not calculated as the number of identified OR genes were smaller than two. Values for vision, hearing, and circadian rhythm are *d*_*N*_/*d*_*S*_ ratios of each set of sensory system associated genes. For avian orders including two or more genomes, *d*_*N*_/*d*_*S*_ ratios of the ancestral branches were used. Three avian orders in red are nocturnal.

Changes in gene expression are thought to underlie many of the phenotypic differences between species [33]. Therefore, we carried out cross-species comparison of gene expression among the blood transcriptomes from 13 raptors (five owls, four accipitrids, and four falconids) and five non-raptor birds. We found that several vision-associated genes [17, 25] were differentially expressed in the raptor orders (*P* <0.05, moderated t-test; Additional file 1: Figures. S7 and S8, and Additional file 2: Datasheets S8-11). For example, *PDCL* (lowly-expressed) and *WFS1* (highly-expressed) genes were differentially expressed specific to the owls. Interestingly, we could also find several circadian rhythm-related genes that were differentially expressed between the nocturnal and diurnal raptors. Three circadian rhythm-associated genes (*ATF4, PER3*, and *NRIP1*) were lowly expressed and two genes (*BTBD9* and *SETX*) were highly expressed in the owls, whereas *ATF4* and *SIRT1* in the falconids and *NRIP1* in the accipitrids were highly expressed. These results likely indicate that selectively driven expression switches contributed to nocturnal adaptation of owls [33].

## Conclusions

Our study provides whole genome assemblies of Eurasian eagle-owl, oriental scops-owl, eastern buzzard, and common kestrel, as well as a suite of whole-genome resequencing and transcriptome data from birds of prey. This is the first in-depth genomics study comparing the three raptor orders, and we identified a number of shared molecular adaptations associated with a predatory lifestyle. Furthermore, compared with diurnal birds, owls and other nocturnal birds showed distinct genomic features, especially in sensory systems. While functional studies of candidate genes will be needed to understand the molecular mechanisms of adaptation, these results provide a genome-wide description and gene candidates of adaptations that have allowed each of these three raptor groups to evolve into diverse, ecologically dominant apex predators.

## Methods

### Sample and genome sequencing

All blood samples used for genome and transcriptome sequencing were collected from individuals being euthanized during wound treatment of rescued animals, except blood samples of *A. flammeus, O. semitorques*, and *P. ptilorhynchus* that were obtained from the live individuals during medical check-up at the wildlife rescue center. Muscle tissues samples collected in 2017 were obtained from the fresh carcasses.

To build reference genome assemblies of the four raptor species (Eurasian eagle-owl, oriental scops-owl, eastern buzzard, and common kestrel), we constructed eleven genomic libraries with various insert sizes (Illumina short insert and long mate-pair libraries) for each species, according to the manufacturer’s protocol. The libraries were sequenced using Illumina HiSeq platforms. The remaining twelve raptor and four non-raptor bird samples were re-sequenced using Illumina HiSeq platforms with a short-insert libraries. Blood transcriptomes of ten raptors and four non-raptor birds were sequenced using Illumina HiSeq platforms according to the manufacturer’s instructions.

### Genome assembly and annotation

To assemble the raptor genomes, PCR duplicated, sequencing and junction adaptor contaminated, and low quality (Q20) reads were filtered out. The short-insert and long-mate library reads were trimmed into 90bp and 50bp, respectively to remove low-quality bases at the ends of the reads. The quality-filtered reads were used to assemble the four raptor genomes using the SOAPdenovo2 software [10]. We applied various *K*-mer values (33, 43, 53, and 63) to obtain fragments with long contiguity. In this process, oriental scops-owl genome was assembled poorly when using SOAPdenovo2, probably because of its high level of genomic heterozygosity. Therefore, we also assembled the four raptor genomes using Platanus assembler, which is more efficient for highly heterozygous genomes [11]. To reduce the number of gaps in the scaffolds, we closed the gaps using the short-insert library reads in two iterations. To correct base-pair level errors, we performed two iterations of aligning the short-insert library reads to the gap-closed scaffolds using BWA-MEM [34] and calling variants using SAMtools [35]. In this process, homozygous variants were assumed as erroneous sequences from the assembly process, and thus substituted for the correction purpose.

To select final high-quality reference assemblies for the four raptors, we annotated all assemblies and evaluated quality of each assembly. We first searched the genomes for tandem repeats and transposable elements using Tandem Repeats Finder (version 4.07b) [36], Repbase (version 19.03) [37], RepeatMasker (version 4.0.5) [38], RMBlast (version 2.2.28) [39], and RepeatModeler (version 1.0.7) [40]. The protein-coding genes were predicted by combining *de novo* and homology-based gene prediction methods with the blood transcriptome data for each assembly. For the homology-based gene prediction, we searched for avian protein sequences from the NCBI database using TblastN (version 2.2.26) [41] with an *E*-value cutoff of 1E-5. The matched sequences were clustered using GenBlastA (version 1.0.4) [42] and filtered by coverage and identity of >40% criterion. Gene models were predicted using Exonerate (version 2.2.0) [43]. For the *de novo* gene prediction, AUGUSTUS (version 3.0.3) [44] was used with the blood transcriptome for each species. We filtered out possible pseudogenes having premature stop-codons and single exon genes that were likely to be derived from retro-transposition. The assembly and gene annotation qualities were assessed by aligning independently *de novo* assembled transcripts using the Trinity software [45] and by searching for evolutionary conserved orthologs using BUSCO software [46]. By considering the assembly statistics (e.g., N50 values and assembled sequence length) and the completeness of the genome assembly, final high-quality reference assemblies for the four raptors were obtained. Genome, transcriptome, and protein sequences for other comparison species were downloaded from the NCBI database. Genes with possible premature stop-codons were excluded in the comparative analyses. The northern-spotted owl’s genome and protein sequences were acquired from the Zenodo linked in the published paper [8].

### Comparative evolutionary analyses

Orthologous gene families were constructed for avian genomes using the OrthoMCL 2.0.9 software [47]. To estimate divergence times of the 25 avian representatives, protein sequences of the avian single-copy gene families were aligned using the MUSCLE program [48]. The poorly aligned regions from the alignments were trimmed using the trimAl software [49]. The divergence times were estimated using the MEGA7 program [50] with the phylogenetic tree topology of published previous studies [1, 6] and the TimeTree database [51]. The date of the node between chicken and rock dove was constrained to 98 million years ago (MYA), chicken and brown kiwi was constrained to 111 MYA and common ostrich and brown kiwi was constrained to 50-105 according to the divergence times from TimeTree. To estimate divergence times among the birds of prey, the date of the node between downy woodpecker and Eurasian eagle-owl constrained to 61-78 MYA and common kestrel and budgerigar was constrained to 60-80 MYA according to the divergence times from the previous studies [1, 6] and TimeTree. A gene family expansion and contraction analysis for the ancestral branches of the three bird of prey orders was conducted using the CAFÉ program [52] with a *P* <0.05 criterion. The significantly different gene family sizes of the present species were identified by performing the Mann-Whitney *U* test (*P* <0.05).

To identify selection at the gene sequence level, two orthologous gene sets were compiled, as previously reported [3]: the single-copy orthologs among avian species and representative genes from multiple-copy orthologs. The representative genes from multiple-copy orthologs were selected, if all species’ protein sequences are reciprocally best matched to a chicken protein sequence using BLASTp with an *E*-value cutoff of 1E-5. PRANK [53] was used to construct multiple sequence alignments among the orthologs. The CODEML program in PAML 4.5 was used to estimate the *d*_*N*_/*d*_*S*_ ratio (non-synonymous substitutions per non-synonymous site to synonymous substitutions per synonymous site) [14]. The one-ratio model was used to estimate the general selective pressure acting among comparison species. The two-ratio model (model=2) was used to ensure that the *d*_*N*_/*d*_*S*_ ratio is difference between foreground species (raptors and nocturnal birds, respectively) and other species. Additionally, the *d*_*N*_/*d*_*S*_ ratios for each order-level branch of raptors and nocturnal birds were used to confirm if the foreground *d*_*N*_/*d*_*S*_ ratio is not biased to a specific raptor and nocturnal bird order. The branch-site test was also conducted [15]. Statistical significance was assessed using likelihood ratio tests with a conservative 10% false discovery rate criterion.

We identified target species-specific amino acid sequences. To filter out biases derived from individual-specific variants, we used all of the raptor re-sequencing data by mapping to the Eurasian eagle-owl genome for Strigiformes, the eastern buzzard genome for Accipitriformes, and the common kestrel genome for Falconiformes. The mapping was conducted using BWA-MEM, and consensus sequences were generated using SAMtools with the default options, except the “-d 5” option. When we identified the specific amino acid sequences, protein sequences of other birds from the NCBI database were also compared. We also checked multiple-sequence alignments manually to remove artifacts. To identify genetic diversity based on heterozygous SNV rates, variants were also called using Sentieon pipeline [54] with the default options, except the “--algo Genotyper” option. The heterozygous SNV rates were calculated by dividing the total number of heterozygous SNVs by the length of sufficiently mapped (>5 depth) genomic regions.

To identify HCRs in the three raptor orders and Passeriformes, we scanned genomic regions that show significantly reduced genetic variation by comparing variations of each window and whole genome as previously suggested [12]. In the case of Passeriformes, whole genome data of four Passeriformes species (medium ground-finch, white-throated sparrow, common canary, and collared flycatcher) were mapped to the zebra finch genome assembly, and then variants were identified using the same methods used for the three raptor orders. Genetic variation was estimated by calculating the number of different bases in the same order genomes within each 100 Kb window. *P*-value was calculated by performing Fisher’s exact test to test whether the genetic variation of each window is significantly different from that of whole genome. Only adjusted *P*-values (*q*-values) [55] of <0.0001 were considered significant. The middle 10 Kb of each significantly different window were considered as HCRs.

For functional enrichment tests of candidate genes, GO annotations of chicken, zebra finch, turkey, flycatcher, duck, anole lizard, and human genomes were downloaded from the Ensembl database [56] and used to assign the avian protein-coding genes with GO categories. A KEGG pathway was assigned using KAAS [57]. Functional information of candidate genes was retrieved from the GO, KEGG, UniProt [58], and GeneCards [17] databases.

### *De novo* transcriptome assembly and differentially expressed genes

The blood transcriptome data were assembled using Trinity software. Contaminated transcripts were searched for bacteria and fungi sequence from the Ensembl database using BLASTN, and filtered by identity of > 95% and *E*-value cutoff of 1E-6 criteria. Coding sequence (CDS) were predicted using TransDecoder [45, 59]. To identify differentially expressed genes, RNA reads were aligned to the reference genome (species whole genome assembled) or the assembled transcripts (species without reference genome) using TopHat2 software [60]. The number of reads that were mapped to orthologous genes were counted using HTSeq-0.6.1 software [61] and then converted into RPKM (Reads per kilobase per million mapped reads) value. The RPKM values were normalized with the Trimmed Mean of M-values (TMM) [62] correction using the R package edgeR [63]. The significance of differential expression was calculated by the moderated t-test [64] (ebayes function) using the R package limma (*P* <0.05) [65].

### Sensory system and beak development associated gene analysis

To compare the olfactory sense across avian clades, we collected a total of 215 chicken olfactory receptor (OR) gene sequences (functional only) from a previously published paper [66]. These ORs were then searched against the 25 avian species genomes using TblastN with default parameters. For OR candidates lacking start/stop codons, we searched 90bp upstream to find start codons and 90bp downstream to find stop codons. After collecting sequences for each species, the CD-HIT program [67] was used to remove redundant sequences with an identity cut-off of 100%. A Pfam [68] search against sequences using hmmer-3.1 program [69] with an *E*-value cutoff of 1.0 was used to identify sequences that contained 7tm_4 domain. To assign OR clades and filter out non-OR genes, the multiple sequence alignments and phylogenetic analysis were conducted with previously clade-assigned OR and non-OR genes of human, anole lizard, and chicken [70] using ClustalW2 program [71]. The remaining OR candidates were classified into three categories: 1) intact genes with normal start and stop codons and longer than 215 amino acid sequences, thus can code for seven transmembrane domains; 2) partial genes without start and/or stop codons; and 3) pseudogenes with frameshift mutations and/or premature stop codons. OR genes have evolved by multiple duplications and display a large number of pseudogenes, which makes the assembly of OR regions challenging and complicates the annotation process of OR genes [5, 13, 72, 73]. To overcome these issues, we also calculated the diversity of OR genes by Shannon entropy [74] using BioEdit [75] as previously suggested [5, 13]. Amino acid positions with above 20% of gaps were excluded, and entropy was averaged across all amino acid positions.

The vision system associated genes were retrieved from previous studies [5, 13]. Hearing associated genes were retrieved from the AmiGO database [76] using GO categories related to hearing [5]. Circadian rhythm related genes were retrieved from the AmiGO database using “biorhythm/circadian” as search keywords. For the beak development analysis, beak development associated genes were retrieved from the falcon genome study [3]. The protein sequences with the same gene name were aligned using ClustalW2 and manually inspected one by one for quality. A total of 50 beak development associated genes shared by at least two Strigiformes, two Accipitriformes, and two Falconiformes, and 402 sensory system associated genes (64 genes for vision, 219 genes for hearing, and 133 genes for circadian rhythm) shared by the brown kiwi, chuck-will’s-widow, and at least two Strigiformes were included for selection constraint (the *d*_*N*_/*d*_*S*_ ratio) analyses.

## Supporting information

Supplemental datasheet

## Additional files

**Additional file 1: Figures S1-S8, Tables S1-S40**, and Supplementary Methods.

**Additional file 2: Datasheet S1.** PSGs in the ancestral branch of Strigiformes. **Datasheet S2.** PSGs in the ancestral branch of Accipitriformes. **Datasheet S3.** PSGs in the ancestral branch of Falconiformes. **Datasheet S4.** PSGs in chuck-will’s-widow genome. **Datasheet S5.** PSGs in brown kiwi genome. **Datasheet S6.** List of genes showing a high level of GC3 biases in the raptors. **Datasheet S7.** List of genes showing a high level of GC3 biases in the nocturnal birds. **Datasheet S8.** Differentially expressed genes in Strigiformes. **Datasheet S9.** Differentially expressed genes in Accipitriformes. **Datasheet S10.** Differentially expressed genes in Falconiformes. **Datasheet S11.** Differentially expressed genes in raptors

## Abbreviations

PSG: Positively selected gene
GC3: Guanine-Cytosine at the third codon position
HCR: Highly conserved genomic region
OR: Olfactory receptor

## Acknowledgements

Korea Institute of Science and Technology Information (KISTI) provided us with Korea Research Environment Open NETwork (KREONET), which is the Internet connection service for efficient information and data transfer. We thank ARA Jo for bird illustrations.

## Availability of data and materials

The Eurasian eagle-owl, oriental scops-owl, eastern buzzard, and common kestrel genomes have been deposited at DDBJ/EMBL/GenBank under the accession numbers PYWY00000000, PYXB00000000, PYWZ00000000, and PYXA00000000, respectively. The versions described in this paper are the first versions, PYWY01000000, PYXB01000000, PYWZ01000000, and PYXA01000000. All the DNA and RNA sequencing data have been deposited into the NCBI Sequence Read Archive under the accession number SRP131743.

## Authors’ contributions

The birds of prey genome project was initiated by the National Institute of Biological Resources, Korea. SoonokK, JHY, and JB supervised and coordinated the project. SoonokK, JB, and YSC conceived and designed the experiments. JAK, SGK, JYP, Hwa-JungK, SunghyunK, Hee-JongK, Jin-hoJ, KJN, and JK provided samples, advice and associated information. YSC, JeHoonJ, HMK, OC, SGP, HYL, and JF conducted the bioinformatics data processing and analyses. YSC, JeHoonJ, SoonokK, and JB wrote and revised the manuscript. AM, DPM, JF, JSE, JAW, and CCW reviewed and edited the manuscript.

## Ethics approval and consent to participate

Permissions for the endangered species of Korea or animals listed as natural monument were obtained from the Ministry of Environment (MOE) or from the Cultural Heritage Administration (CHA), respectively (see Additional file 1: Table S3 for detailed sampling and permission information). No animals were killed or captured as a result of these studies.

## Consent for publication

Not applicable.

