## Supplemental figures, tables, and method for "Raptor genomes reveal evolutionary signatures of predatory and nocturnal lifestyles"

**Additional file 1**

**Table of contents**

**List of Supplementary Figures**

Figure S1. Species identification for the new 20 avian species sequenced	4

Figure S2. *K*-mer (*K*=17) analyses of 20 avian genomes sequenced in the present study	10

Figure S3. Genetic diversity in 25 bird of prey species	12

Figure S4. Composition of avian orthologous genes	13

Figure S5. Genomic context among 25 avian species	14

Figure S6. *SLC51A* gene variants in nocturnal birds	15

Figure S7. Differentially expressed genes (DEGs) in the birds of prey species	16

Figure S8. Differentially expressed genes associated with the vision system and circadian rhythm	17

**List of Supplementary Tables**

Table S1. Bird of prey genome and transcriptome data used in this study	19

Table S2. Non-raptor bird genome and transcriptome data used for comparative evolutionary analysis	20

Table S3. Sampling information on bird species sequenced in this study	21

Table S4. Sequencing library statistics used for the four bird of prey genome assemblies	23

Table S5. Filtered sequence information of the four birds of prey	25

Table S6. 17-mer statistics information for 20 avian species	27

Table S7. Global assembly statistics of the four bird of prey genomes	28

Table S8. Assessment of gene coverage by assembled bird of prey transcripts	29

Table S9. Evaluation of the completeness of bird of prey assemblies and gene sets using single-copy orthologs mapping approach	30

Table S10. Protein-coding gene prediction statistics for the four bird of prey genomes	31

Table S11. Transposable element statistics for the four bird of prey genomes	33

Table S12. Whole genome sequencing and mapping statistics for the birds of prey and non-raptor birds	35

Table S13. Variant statistics for the birds of prey and non-raptor birds	36

Table S14. Blood transcriptome *de novo* assembly and mapping statistics	37

Table S15. Genome assembly information for the 25 avian species used in this study	38

Table S16. Commonly enriched Gene Ontology (GO) categories of expanded gene families in the ancestral branches of Strigiformes, Accipitriformes, and Falconiformes	39

Table S17. Commonly enriched KEGG pathways of expanded gene families in the ancestral branches of Strigiformes, Accipitriformes, and Falconiformes	41

Table S18. Gene Ontology (GO) enrichment of gene families that were expanded in the present birds of prey	42

Table S19. List of genes showing accelerated d_N_/d_S_ in the three raptor orders	44

Table S20. Selection constraints of beak development associated genes in the ancestral branches of birds of prey	47

Table S21. Gene Ontology (GO) enrichment of genes that have a high level of GC3 biases in the bird of prey genomes	48

Table S22. Statistics regarding highly conserved regions in Strigiformes, Accipitriformes, Falconiformes, and Passeriformes	49

Table S23. Commonly enriched Gene Ontology (GO) categories of genes in the highly conserved genomic regions (HCRs) of Strigiformes, Accipitriformes, and Falconiformes	50

Table S24. Commonly enriched KEGG pathways of genes in the highly conserved genomic regions (HCRs) of Strigiformes, Accipitriformes, and Falconiformes	52

Table S25. Strigiformes specific GO enrichment of genes in the highly conserved genomic regions (HCRs)	53

Table S26. Strigiformes specifically enriched KEGG pathways of genes in the highly conserved genomic regions (HCRs)	55

Table S27. Accipitriformes specific GO enrichment of genes in the highly conserved genomic regions (HCRs)	56

Table S28. Accipitriformes specifically enriched KEGG pathways of genes in the highly conserved genomic regions (HCRs)	57

Table S29. Falconiformes specific GO enrichment of genes in the highly conserved genomic regions (HCRs)	58

Table S30. Falconiformes specifically enriched KEGG pathways of genes in the highly conserved genomic regions (HCRs)	60

Table S31. Passeriformes specific GO enrichment of genes in the highly conserved genomic regions (HCRs)	61

Table S32. Commonly enriched Gene Ontology (GO) categories of expanded gene families in the common ancestor of Strigiformes, brown kiwi and chuck-will’s widow.	62

Table S33. Commonly enriched KEGG pathways of expanded gene families in the common ancestor of Strigiformes, brown kiwi and chuck-will’s widow	63

Table S34. Gene Ontology (GO) enrichment of gene families that were contracted in size the present nocturnal bird species genomes	64

Table S35. Gene Ontology (GO) enrichment of gene families that were expanded in the present nocturnal bird species genomes	66

Table S36. Gene Ontology (GO) enrichment of PSGs that were shared in two and/or three nocturnal bird groups	67

Table S37. List of genes showing accelerated d_N_/d_S_ in the nocturnal birds	69

Table S38. Olfactory receptors identified in 25 avian genomes	71

Table S39. The diversity of olfactory receptors in 25 avian genomes	72

Table S40. Sensory system associated genes showing accelerated d_N_/d_S_ in the nocturnal birds	73

**Supplementary Methods**

Genome and transcriptome sequencing	74

Species identification	75

Sequence filtering criteria	75

Repeat annotation	76

**Supplementary References**	77

**Supplementary Figures**

**Falconiformes – *COI* gene**

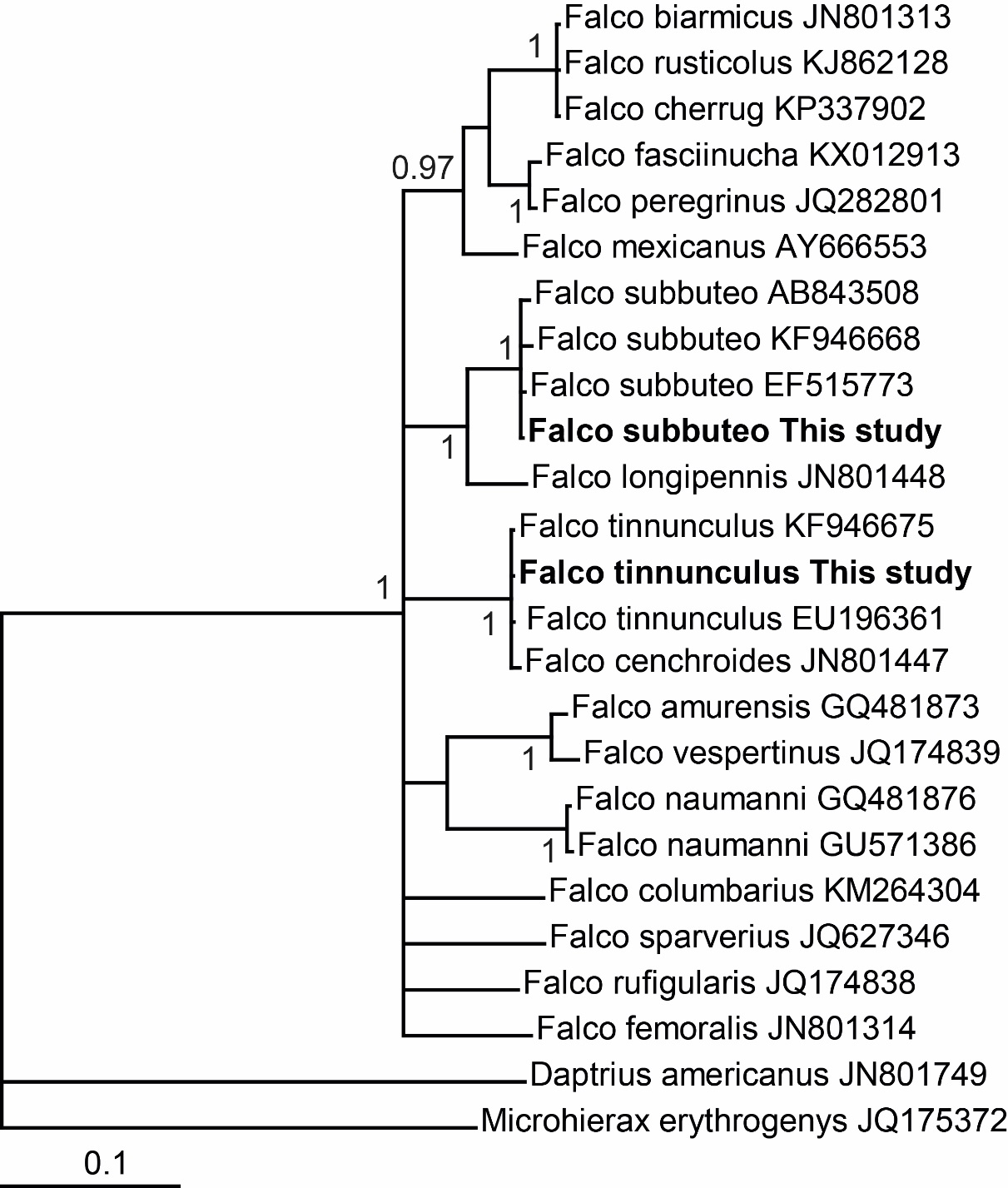

**Accipitriformes – *CYTB* gene**

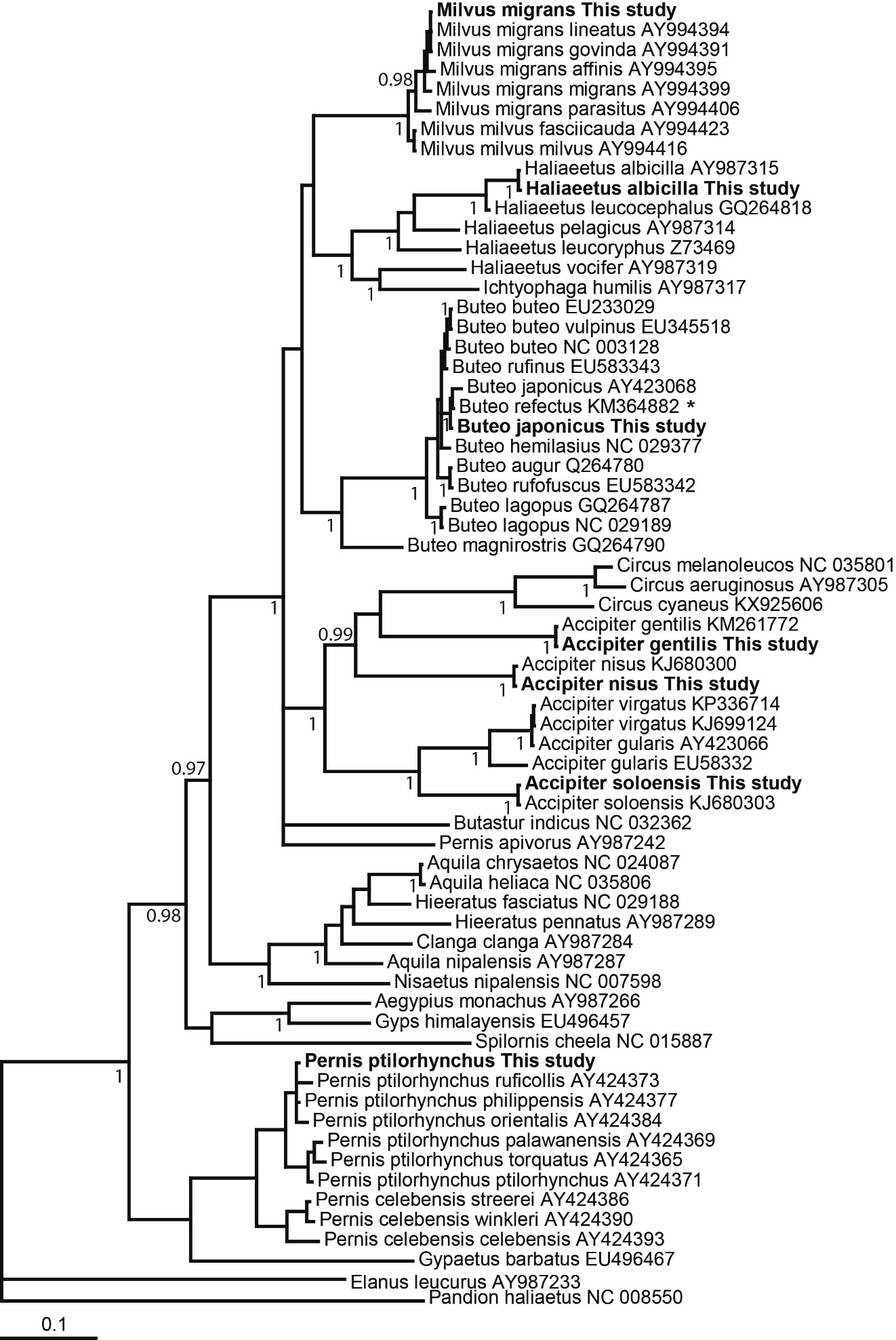

**Strigiformes – *CYTB* gene**

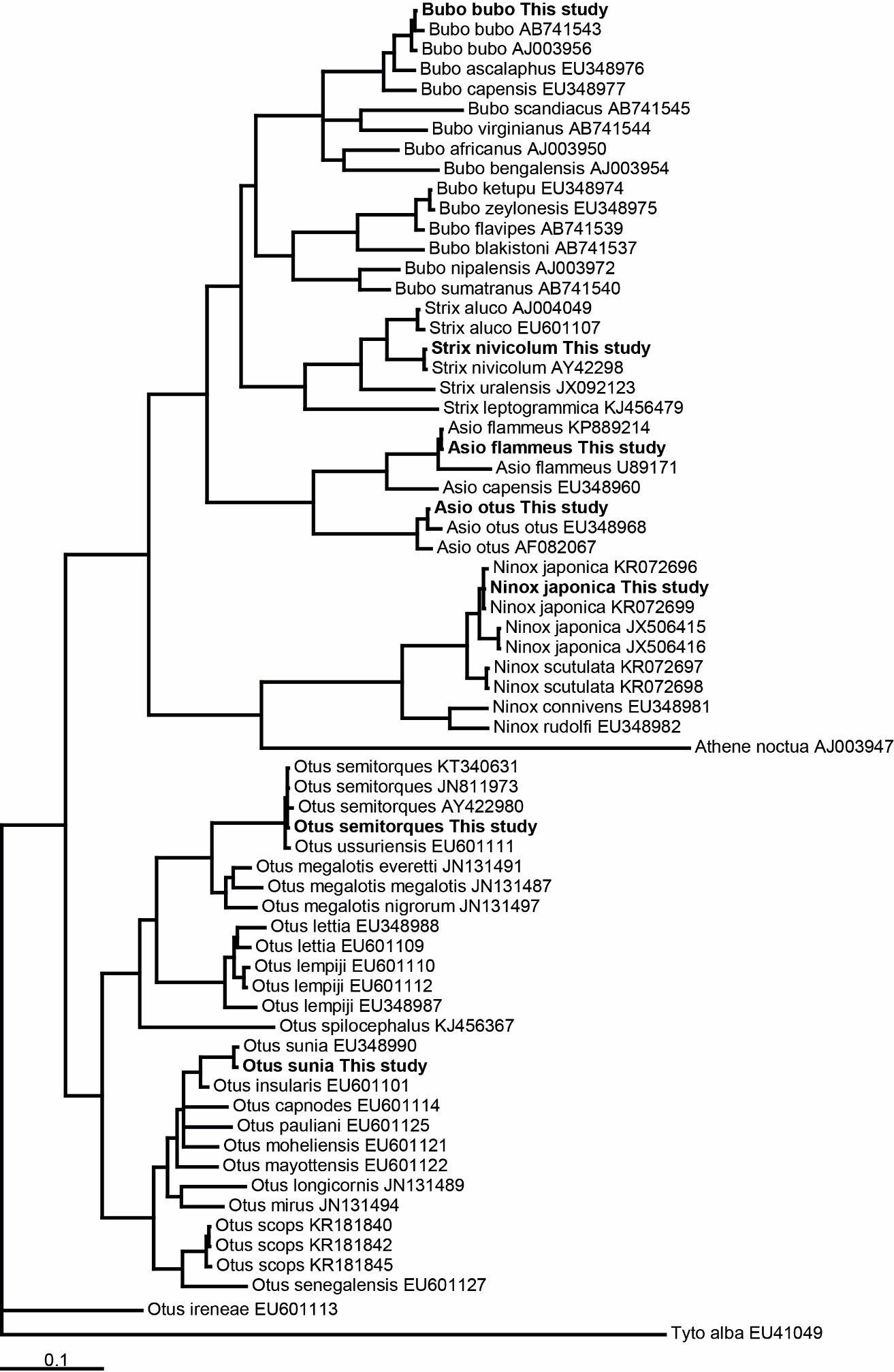

**Ardeidae and Threskiornithidae – *COI* gene**

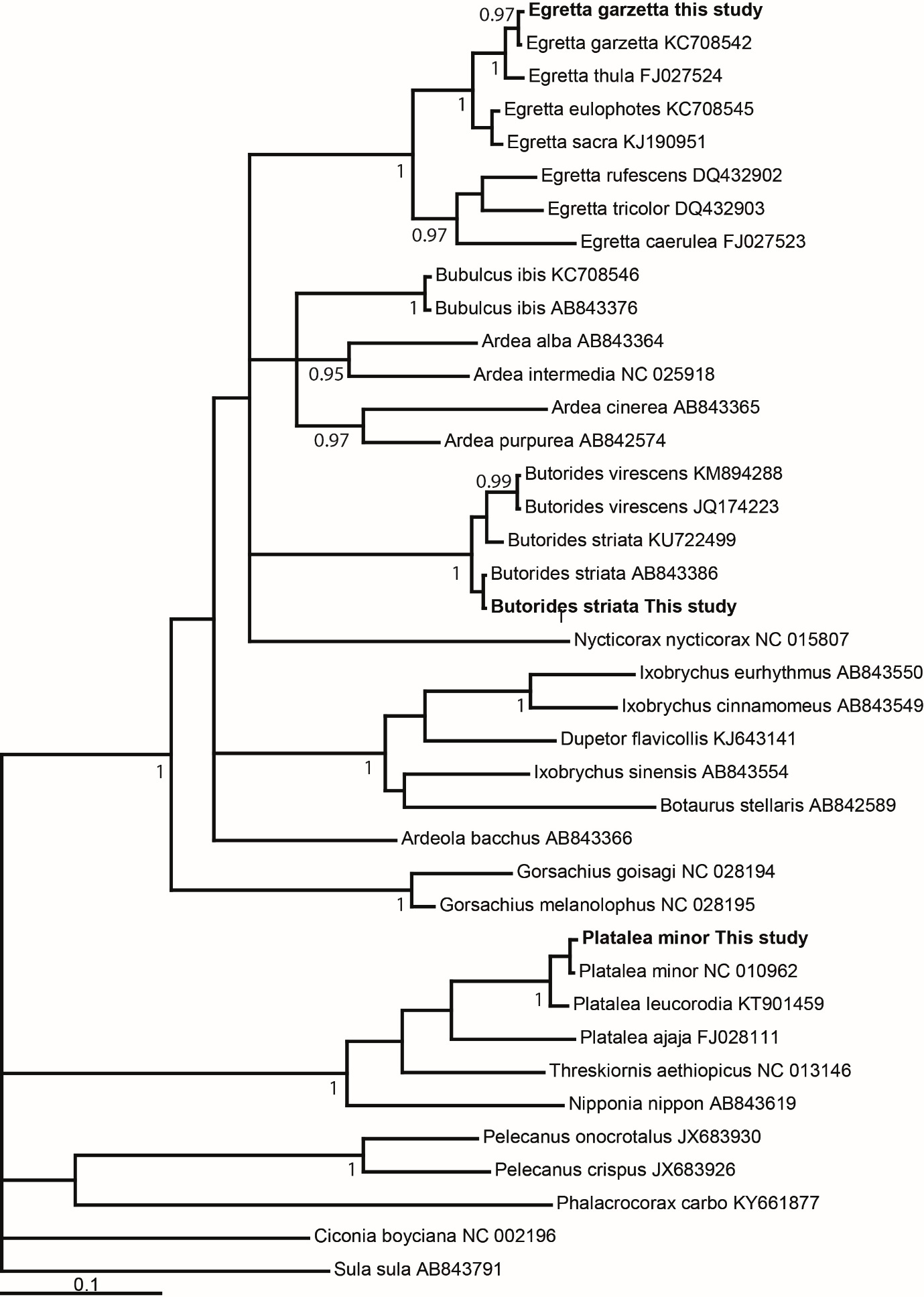

**Picidae – *COI* gene**

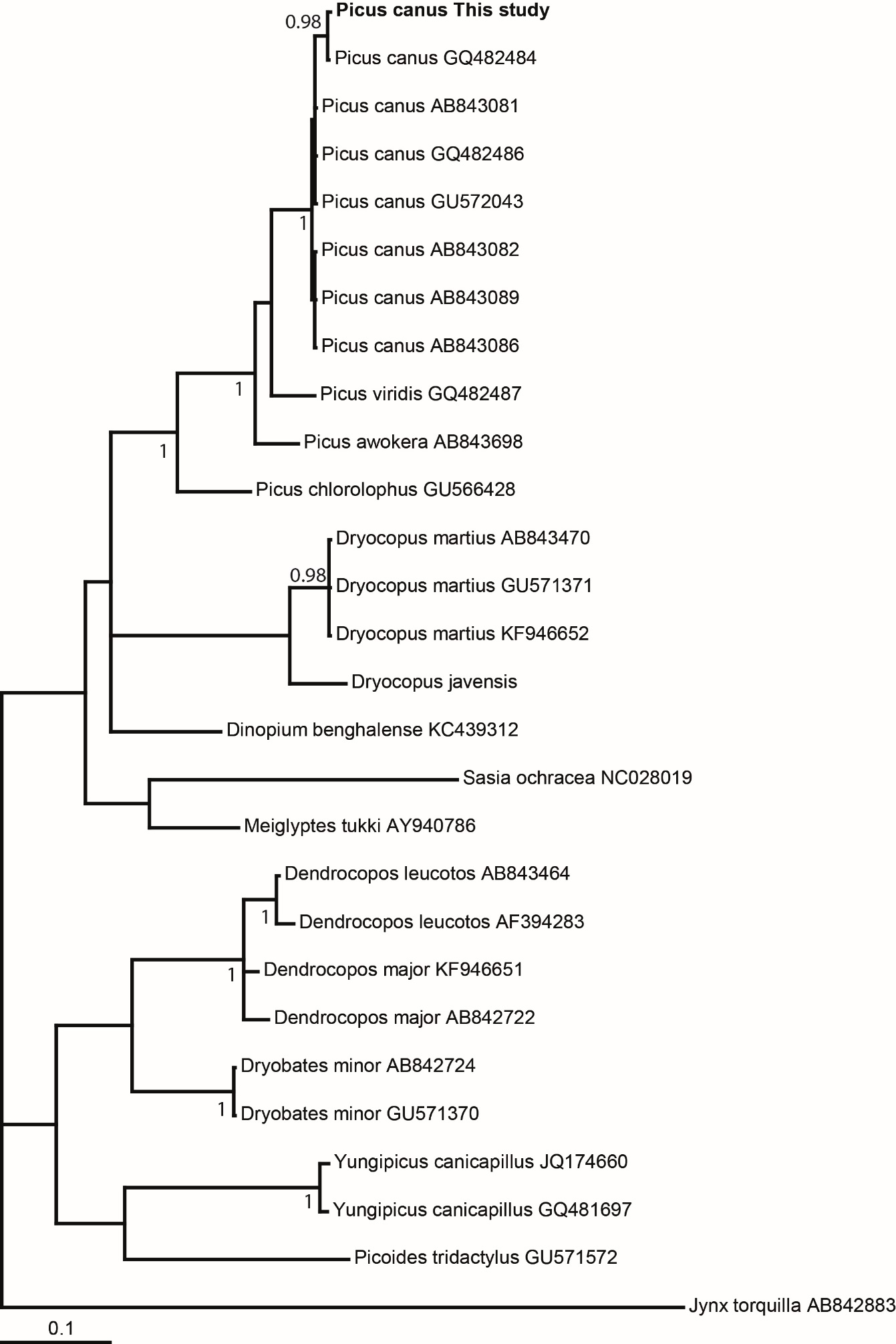

**Figure S1. Species identification for the new 20 avian species sequenced.** The consensus sequences of avian samples were generated by mapping their reads to previously reported mitochondrial sequences (*COI* and *CYTB* genes) for closely related species. The *COI* gene of common kestrel was sequenced by Sanger method. The names that include “This study” were originally sequenced in this study. Numbers close to nodes are posterior probabilities. *This sequence was first attributed to *B. buteo burmanicus*, a junior synonym of *B. refectus*; the sampling locality is outside the known range of *B. refectus* and suggests that it is a misidentified *B. japonicus*.

| 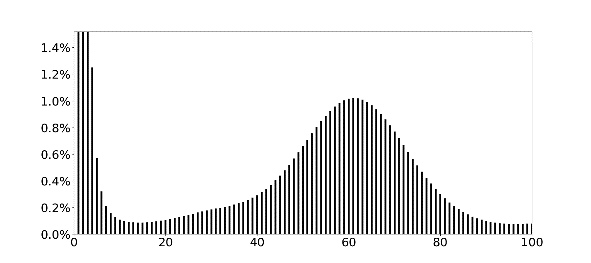 | 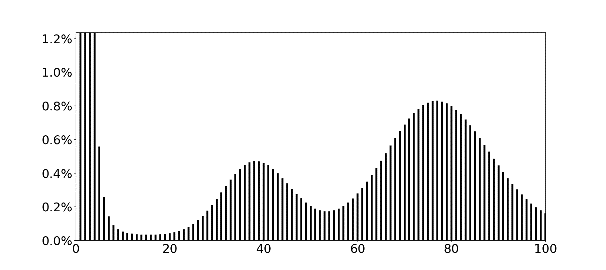 |
| --- | --- |
| Eurasian eagle-owl | Oriental scops-owl |
| 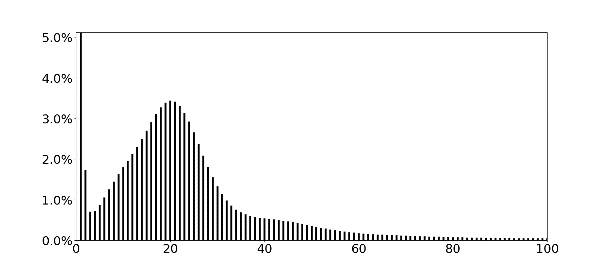 | 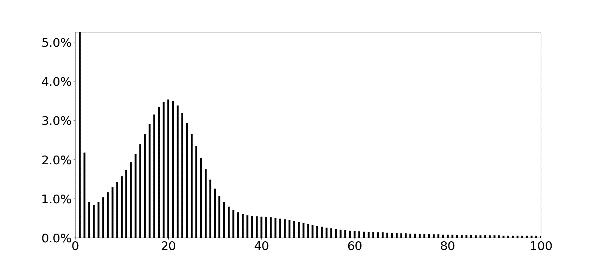 |
| Himalayan owl | Northern boobook |
| 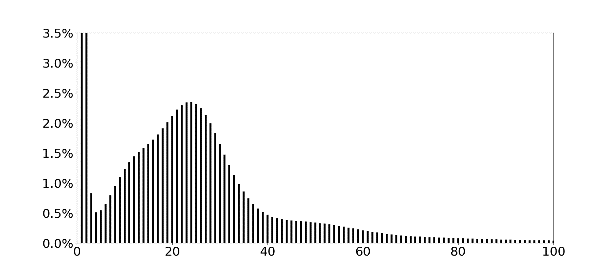 | 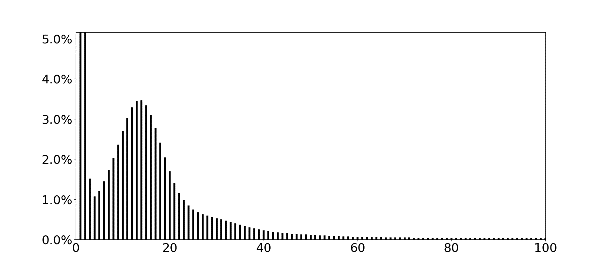 |
| Long-eared owl | Short-eared owl |
| 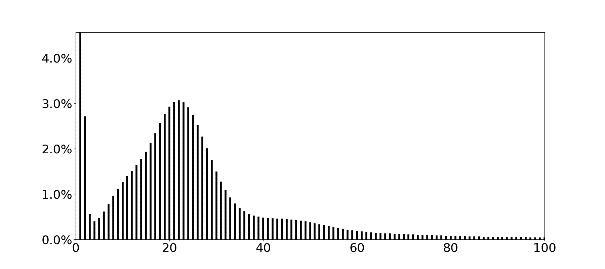 | 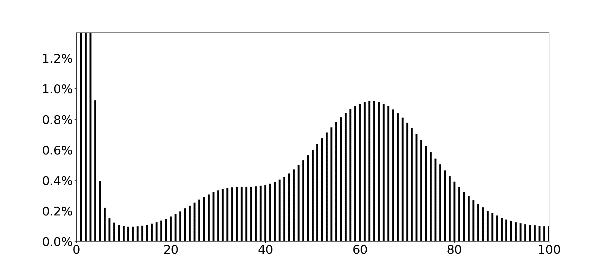 |
| Japanese scops-owl | Eastern buzzard |
| 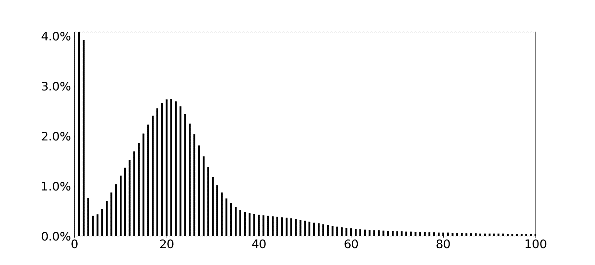 | 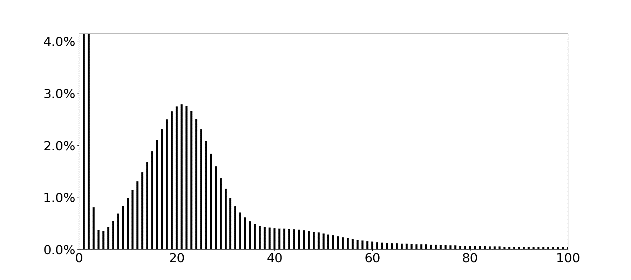 |
| Eurasian sparrowhawk | Northern goshawk |
| 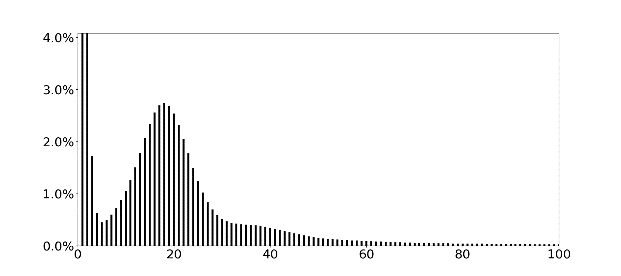 | 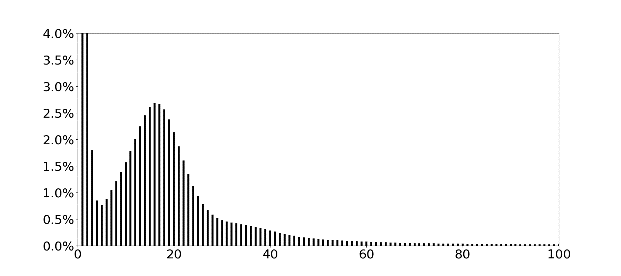 |
| White-tailed eagle | Oriental honey-buzzard |

| 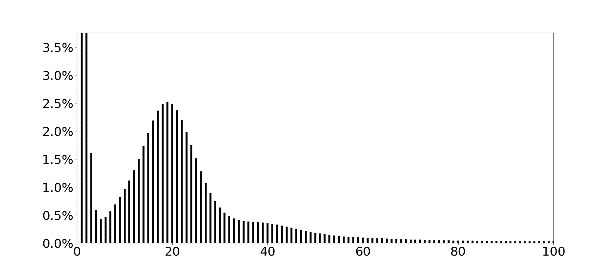 | 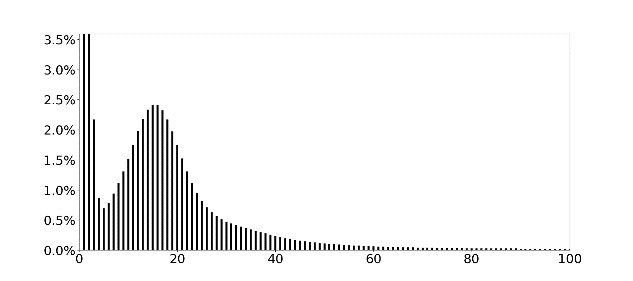 |
| --- | --- |
| Black kite | Chinese sparrowhawk |
| 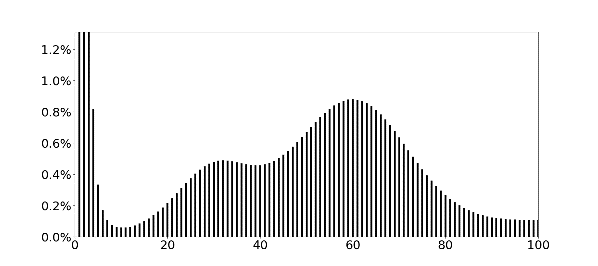 | 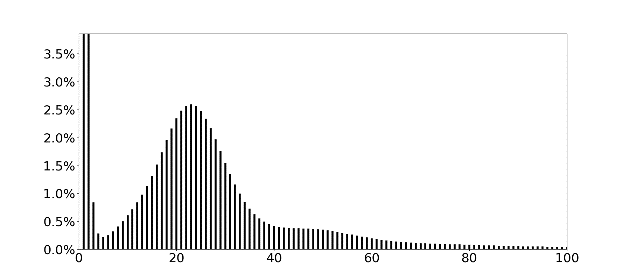 |
| Common kestrel | Eurasian hobby |
| 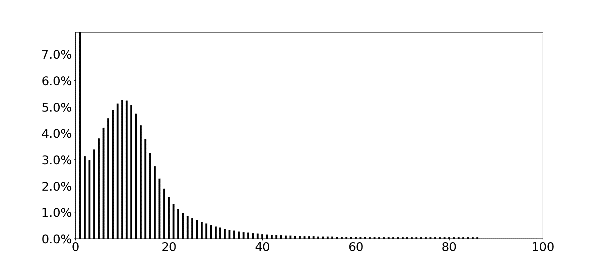 | 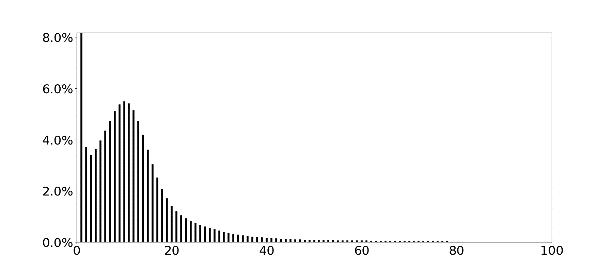 |
| Grey-headed woodpecker | Little egret |
| 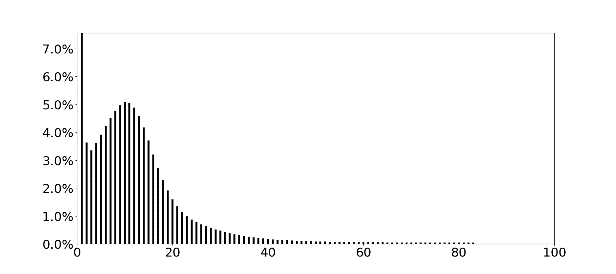 | 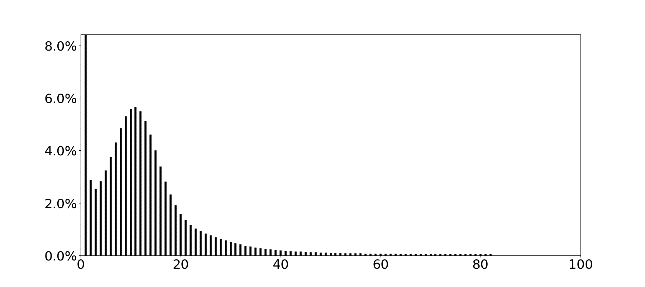 |
| Striated heron | Black-faced spoonbill |

**Figure S2. *K*-mer (*K*=17) analyses of 20 avian genomes sequenced in the present study.** The *x*-axis represents *K*-mer depth, and the *y*-axis represents the proportion of *K*-mer count at that depth.

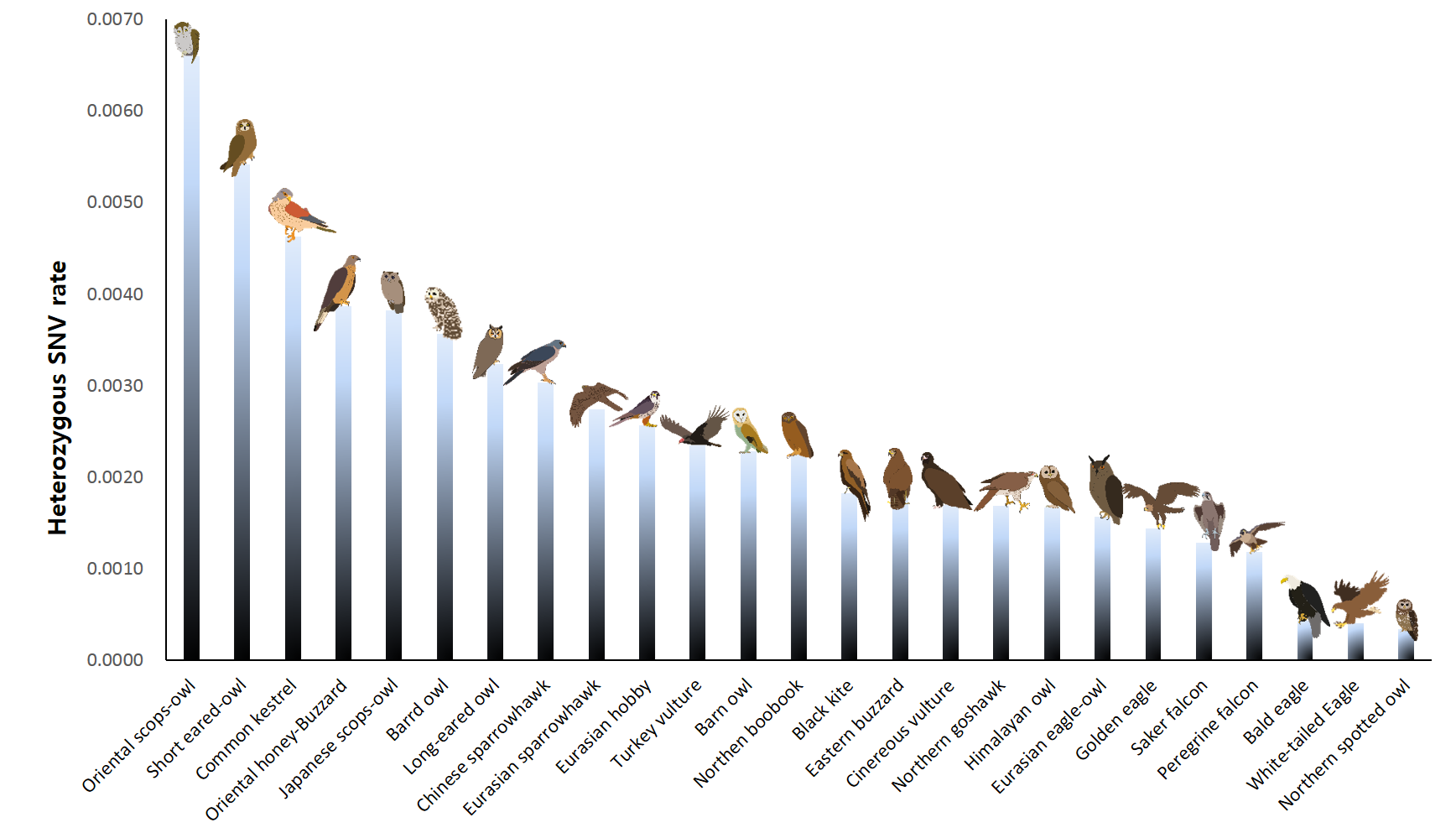

**Figure S3. Genetic diversity in 25 bird of prey species.** The heterozygous SNVs rates (*y*-axis) were calculated by dividing the total number of heterozygous SNVs by the length of sufficiently mapped (>5 depth) genomic regions. The estimated heterozygous SNVs rates were based on single individuals. The heterozygous SNVs rates can be altered according to which reference assembly is used and the assembly quality.

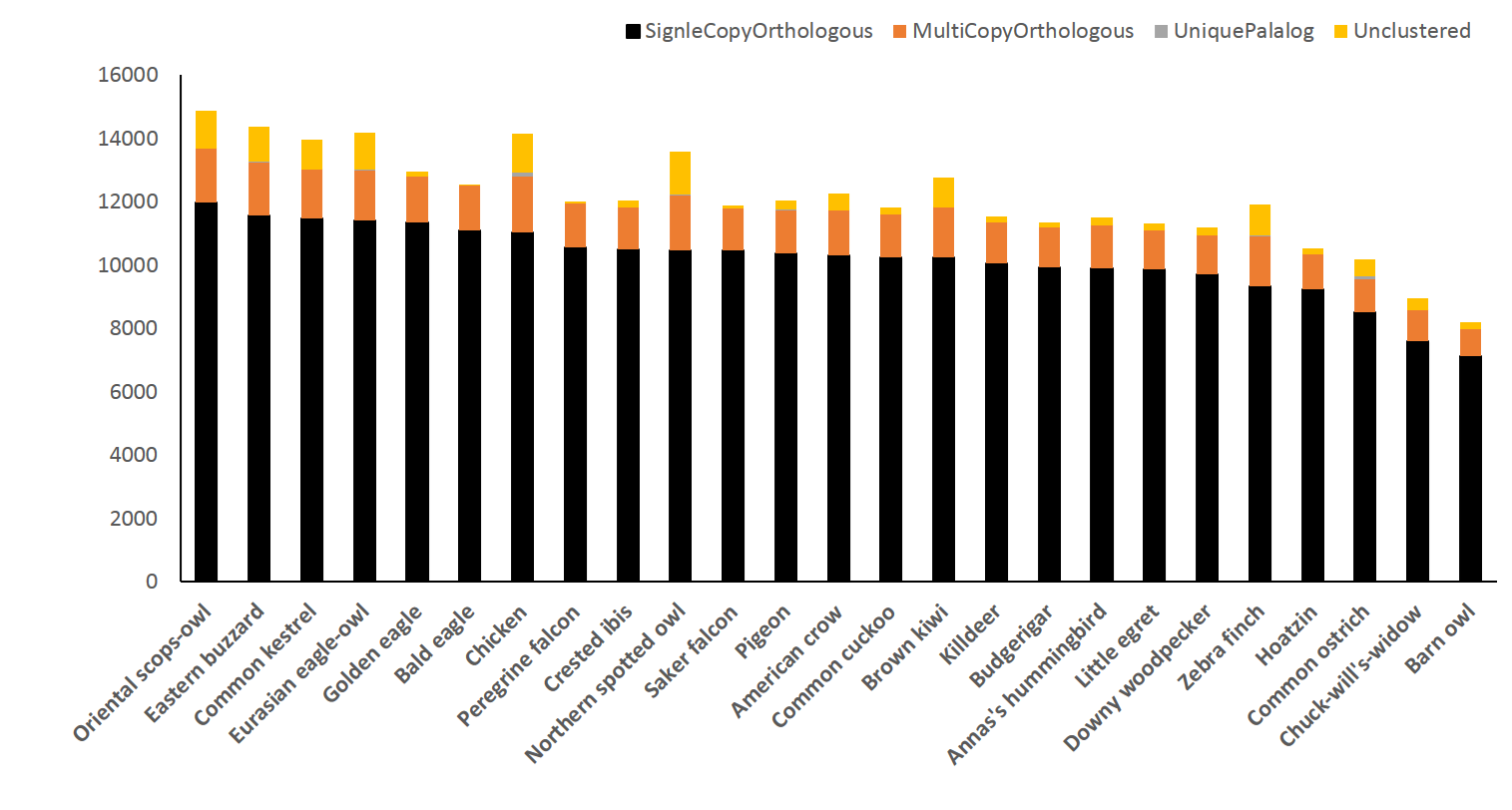

**Figure S4. Composition of avian orthologous genes.** A comparative representation of orthologous and paralogous genes in 25 avian genomes are shown. Two low-quality genomes (chuck-will’s-widow and barn owl) showed low numbers of gene clusters.

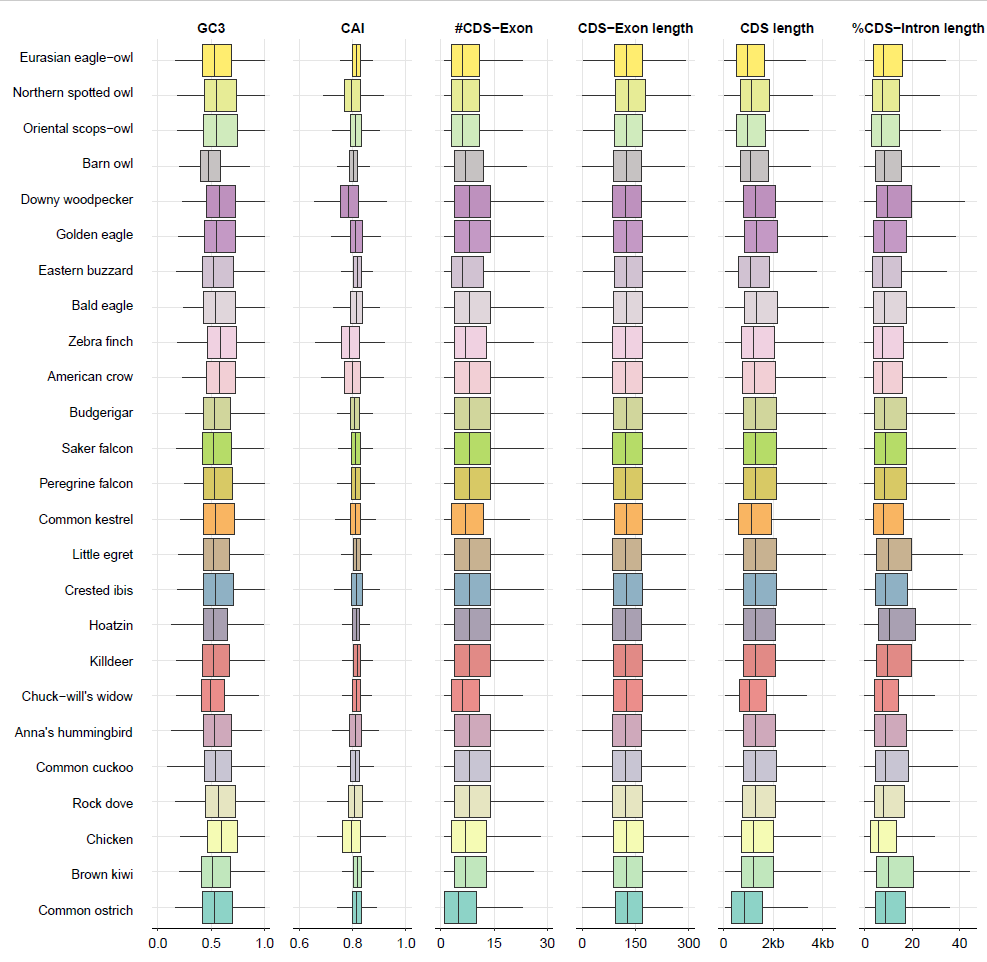

**Figure S5. Genomic context among 25 avian species.** GC3 ratio is the Guanine-Cytosine ratio at the third codon positions. CAI is the codon adaptation index. # CDS is the average number of CDS. Intron/coding regions is average length of sum of intron length divided by coding region length.

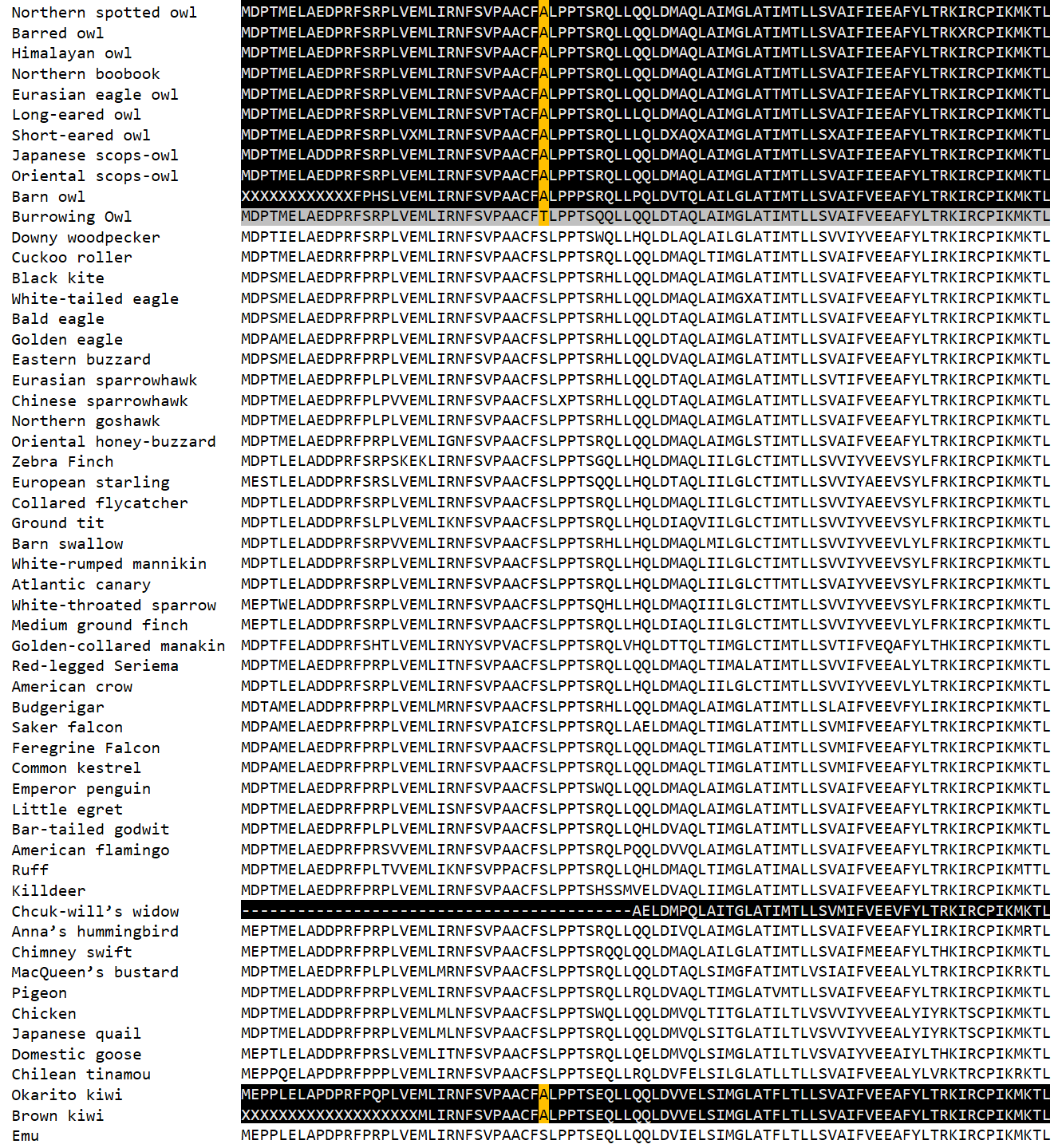

**Figure S6. *SLC51A* gene variants in nocturnal birds.** An amino acid unique to nocturnal birds and burrowing owl (33^th^ residue in the chicken *SLC51A* protein sequence) is highlighted in yellow. Sequences in the nocturnal birds and burrowing owl are highlighted in black and gray, respectively. We could not find the amino acid residue in the chuck-will’s widow, as *SLC51A* gene is partially annotated in cuck-will’s widow genome.

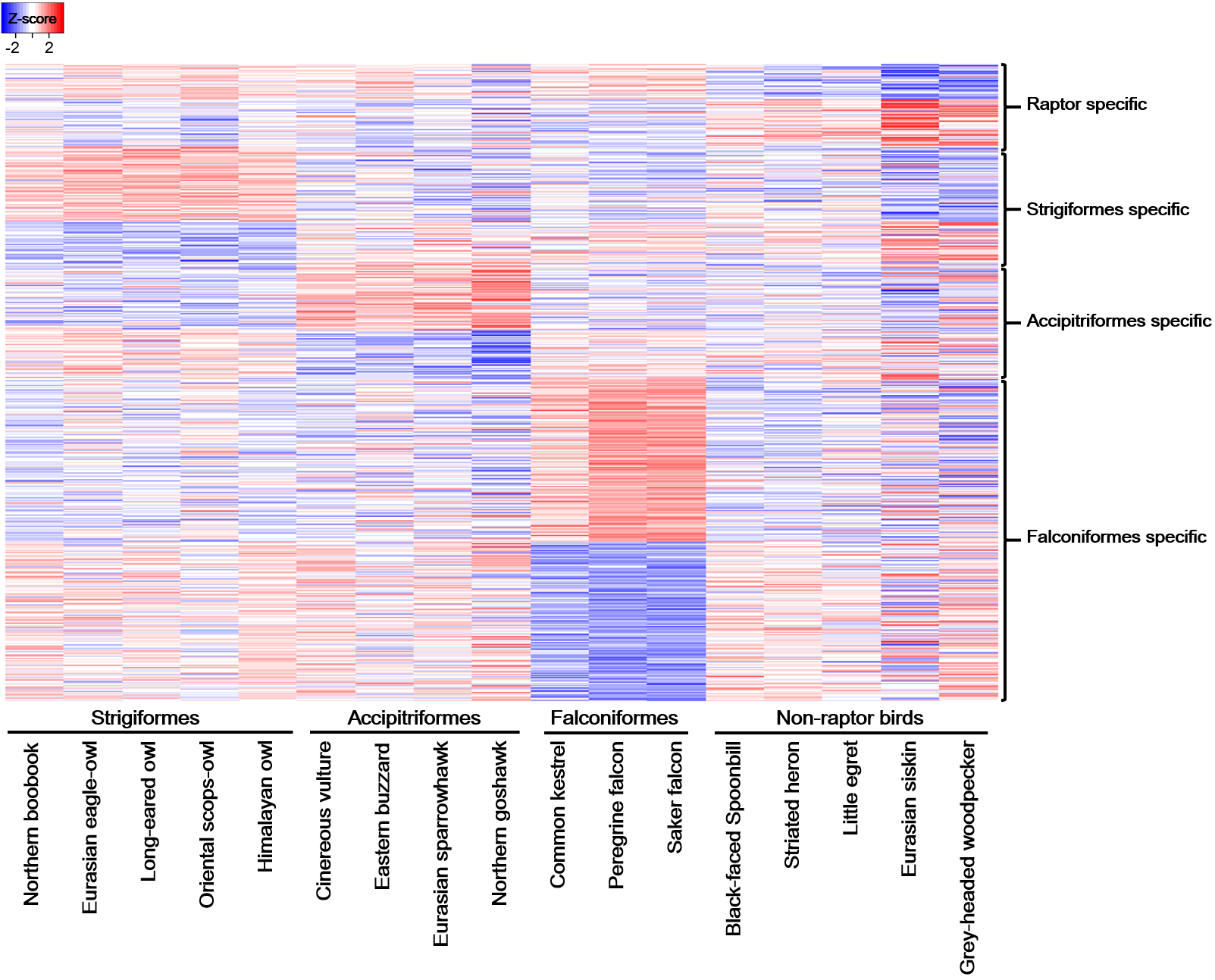

**Figure S7. Differentially expressed genes (DEGs) in the birds of prey species.** *P*-value (*P* <0.05) heatmap of differentially expressed genes in the blood transcriptome of three raptor orders (Strigiformes, Accipitriformes, and Falconiformes). Greater and less than 2-fold expressions are shown in each column. The orders of DEGs were sorted by *P*-values in each target group (raptor-specific, Strigiformes-specific, Accipitriformes-specific, and Falconiformes-specific). The blood transcriptomes of black-faced spoonbill, striated heron, little egret, Eurasian siskin, and grey-headed woodpecker were used as a control group (non-raptor birds).

| 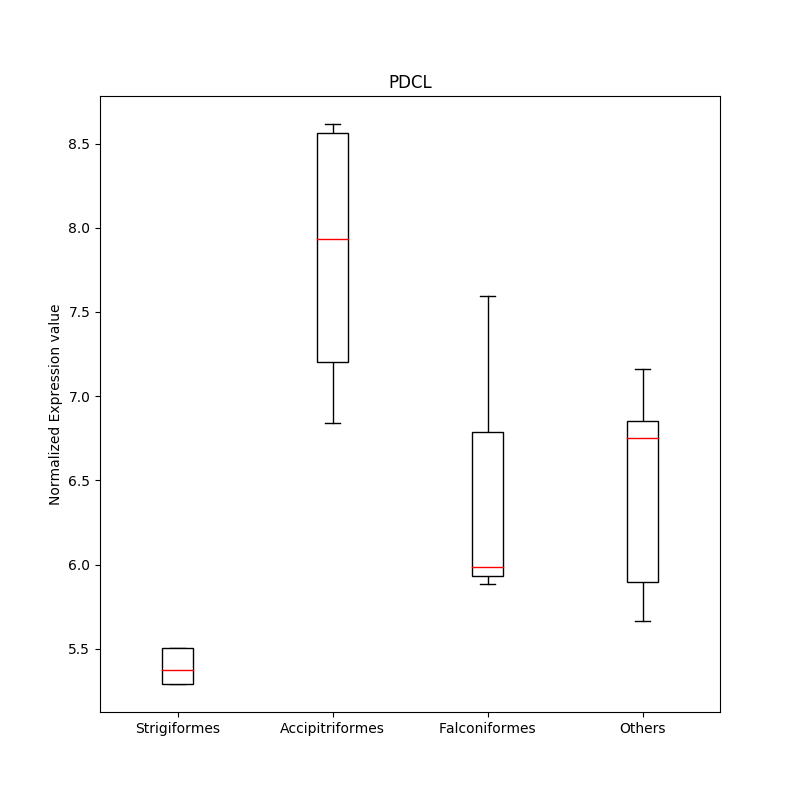 | 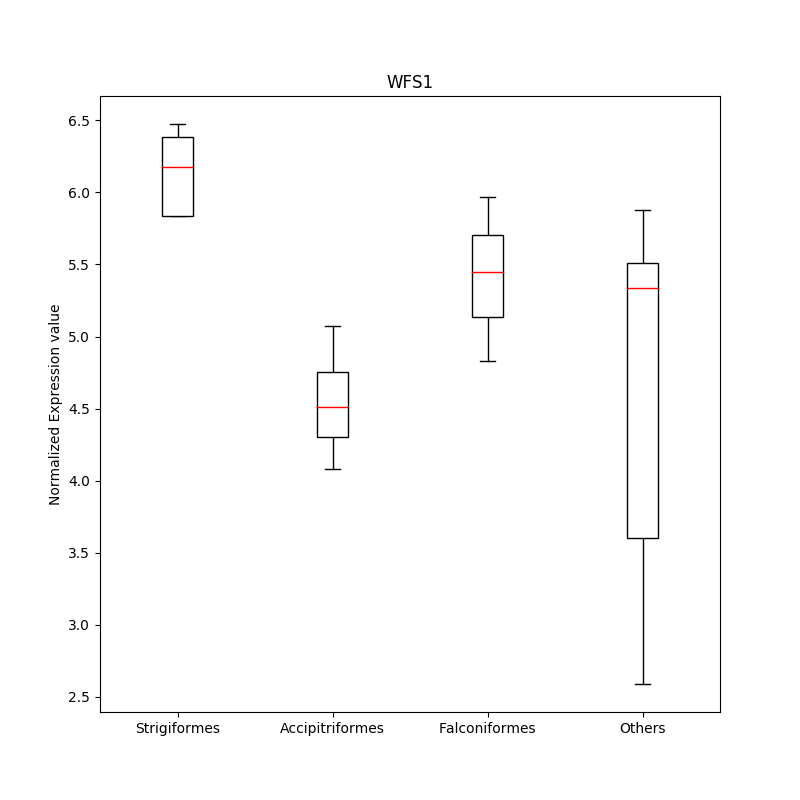 |
| --- | --- |
| *PDCL* gene | *WFS1* gene |

| 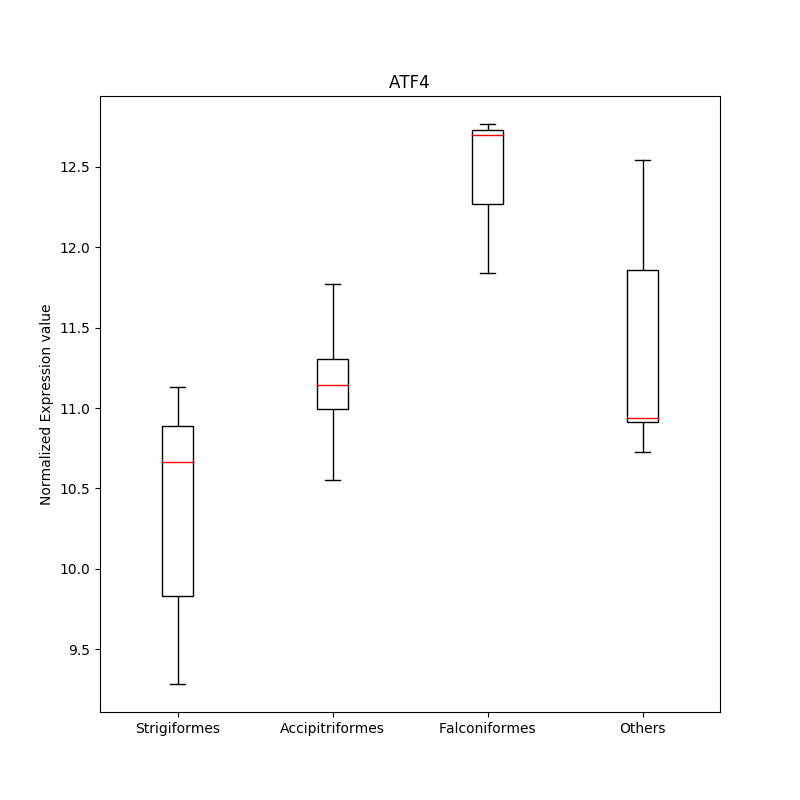 | 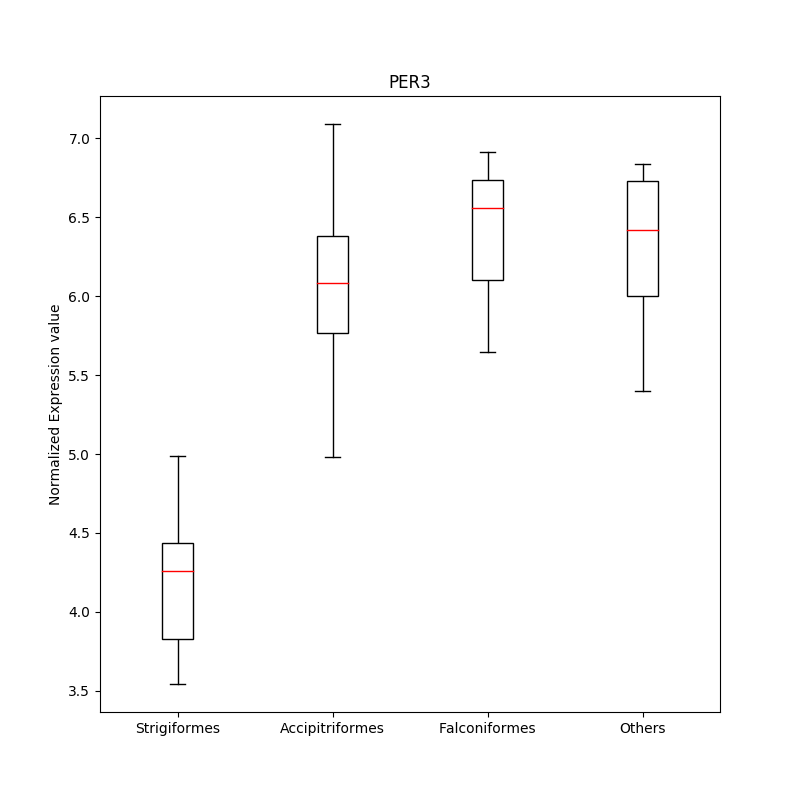 |
| --- | --- |
| *ATF4* gene | *PER3* gene |

| 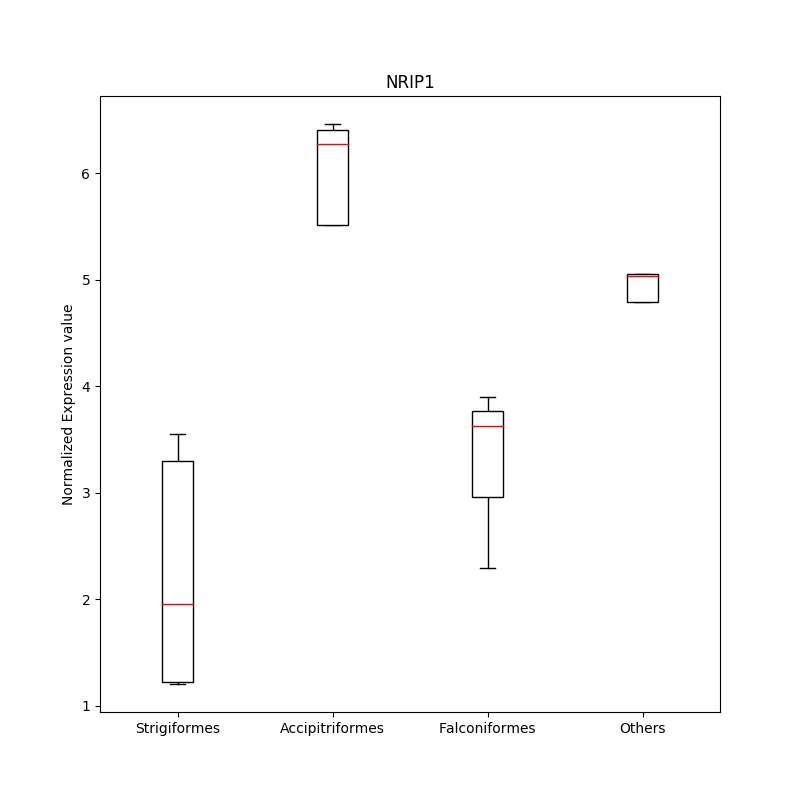 | 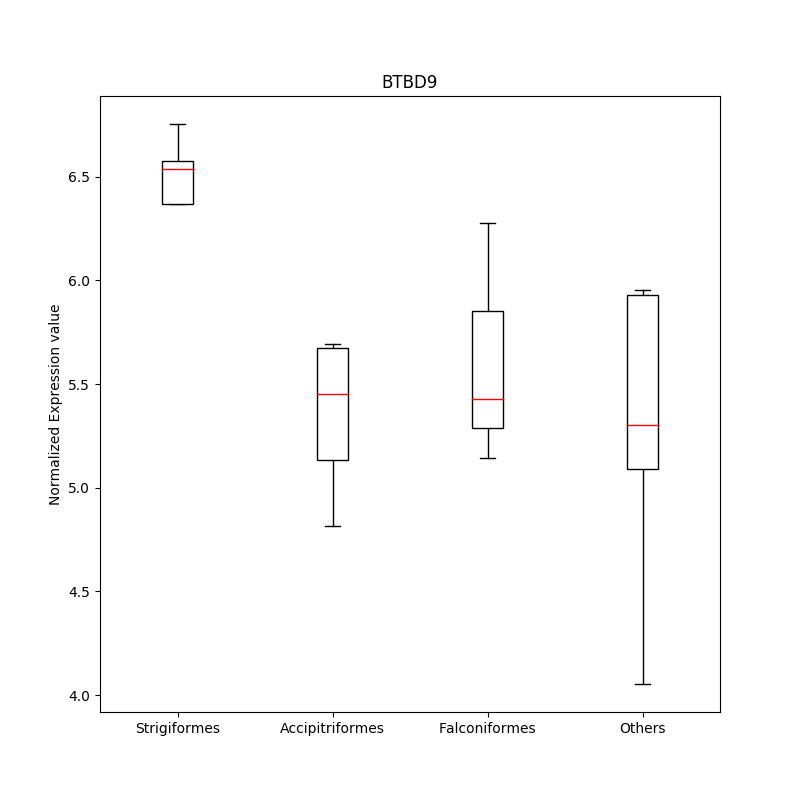 |
| --- | --- |
| *NRIP1* gene | *BTBD9* gene |
| 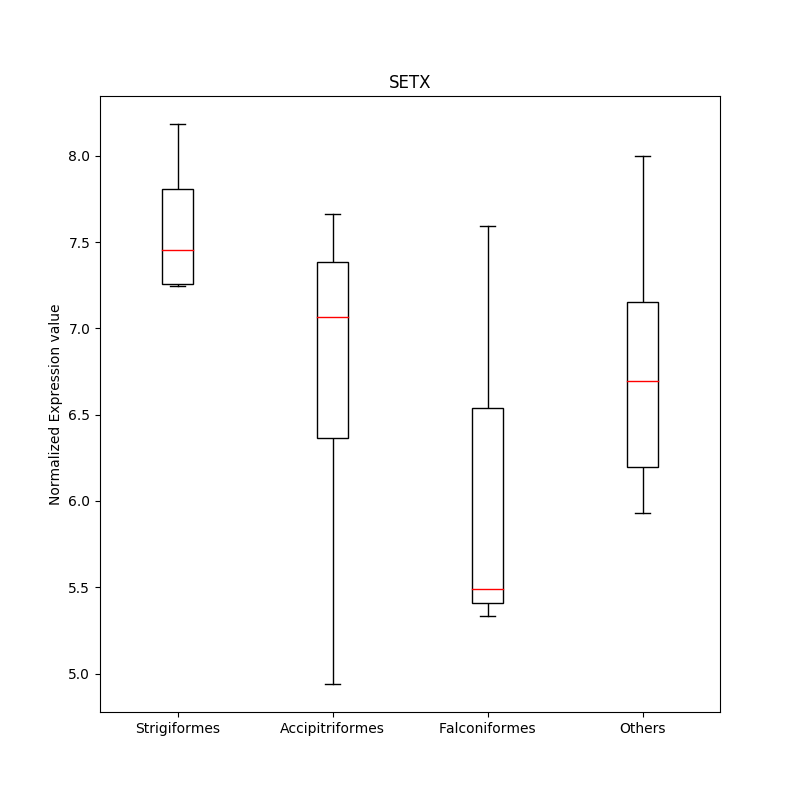 | 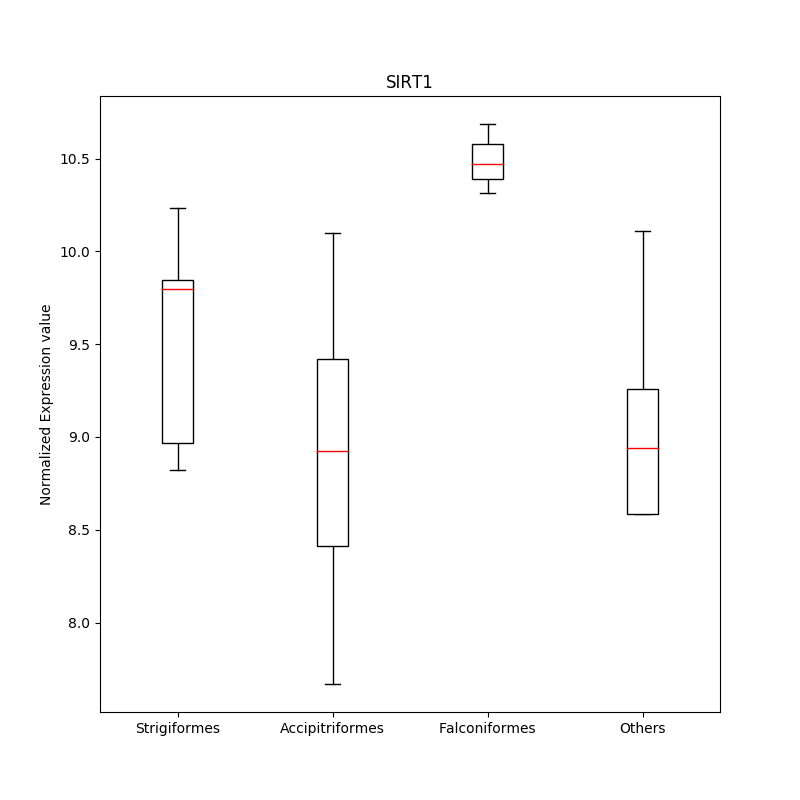 |
| *SETX* gene | *SIRT1* gene |

**Figure S8. Differentially expressed genes associated with the vision system and circadian rhythm.** The normalized expression values were compared among Strigiformes (northern boobook, Eurasian eagle-owl, long-eared owl, oriental scops-owl, and Himalayan owl), Accipitriformes (cinereous vulture, eastern buzzard, Eurasian sparrowhawk, and northern goshawk), Falconiformes (common kestrel, peregrine falcon, and saker falcon), and non-raptor birds (black-faced spoonbill, striated heron, little egret, Eurasian siskin, and grey-headed woodpecker).

**Supplementary Tables**

**Table S1. Bird of prey genome and transcriptome data used in this study**

| Order | Family | Species | Common name | Data | IUCN  Red List | Source |
| --- | --- | --- | --- | --- | --- | --- |
| Strigiformes | Strigidae | *Bubo bubo* | Eurasian eagle-owl | Assembly, Transcriptome | Least Concern | This study |
|  |  | *Otus sunia* | Oriental scops-owl | Assembly, Transcriptome | Least Concern | This study |
|  |  | *Strix nivicolum* | Himalayan owl | Resequencing, Transcriptome | Least Concern | This study |
|  |  | *Strix occidentalis* | Northern spotted owl | Assembly | Near Threatened | 1 |
|  |  | *Strix Varia* | Barred owl | Resequencing | Least Concern | 1 |
|  |  | *Ninox japonica* | Northern boobook | Resequencing, Transcriptome | Least Concern | This study |
|  |  | *Asio otus* | Long-eared owl | Resequencing, Transcriptome | Least Concern | This study |
|  |  | *Asio flammeus* | Short-eared owl | Resequencing | Least Concern | This study |
|  |  | *Otus semitorques* | Japanese scops-owl | Resequencing | Least Concern | This study |
|  | Tytonidae | *Tyto alba* | Barn owl | Assembly  (low quality) | Least Concern | 2 |
| Accipitriformes | Accipitridae | *Buteo japonicus* | Eastern buzzard | Assembly, Transcriptome | Least Concern | This study |
|  |  | *Haliaeetus leucocephalus* | Bald eagle | Assembly | Least Concern | 2 |
|  |  | *Aquila chrysaetos* | Golden eagle | Assembly | Least Concern | 3 |
|  |  | *Accipiter nisus* | Eurasian sparrowhawk | Resequencing, Transcriptome | Least Concern | This study |
|  |  | *Accipiter gentilis* | Northern goshawk | Resequencing, Transcriptome | Least Concern | This study |
|  |  | *Aegypius monachus* | Cinereous vulture | Resequencing, Transcriptome | Near Threatened | 4 |
|  |  | *Haliaeetus albicilla* | White-tailed eagle | Resequencing;  low-quality assembly is available, but not used in this study | Least Concern | This study,  2 |
|  |  | *Pernis ptilorhynchus* | Oriental honey-buzzard | Resequencing | Least Concern | This study |
|  |  | *Milvus migrans* | Black kite | Resequencing | Least Concern | This study |
|  |  | *Accipiter soloensis* | Chinese sparrowhawk | Resequencing | Least Concern | This study |
|  | Cathartidae | *Cathartes aura* | Turkey vulture | Resequencing;  low-quality assembly is available, but not used in this study | Least Concern | 2 |
| Falconiformes | Falconidae | *Falco tinnunculus* | Common kestrel | Assembly,  Transcriptome | Least Concern | This study |
|  |  | *Falco cherrug* | Saker falcon | Assembly, Transcriptome | Endangered | 5 |
|  |  | *Falco peregrinus* | Peregrine falcon | Assembly, Transcriptome | Least Concern | 5 |
|  |  | *Falco subbuteo* | Eurasian hobby | Resequencing, Transcriptome | Least Concern | This study |

**Table S2. Non-raptor bird genome and transcriptome data used for comparative evolutionary analysis**

| Order | Family | Species | Common name | Data | Source |
| --- | --- | --- | --- | --- | --- |
| Piciformes | Picidae | *Dryobates pubescens* | Downy woodpecker | Assembly | 2 |
|  | Picidae | *Picus canus* | Grey-headed woodpecker | Resequencing,  Transcriptome | This study |
| Psittaciformes | Psittaculidae | *Melopsittacus undulatus* | Budgerigar | Assembly | 6 |
| Passeriformes | Corvidae | *Corvus brachyrhynchos* | American crow | Assembly | 2 |
|  | Estrildidae | *Taeniopygia guttata* | Zebra finch | Assembly | 7 |
|  | Thraupidae | *Geospiza fortis* | Medium ground-finch | Resequencing; assemblies are available, but used as resequencing data for HCR analysis | 2 |
|  | Emberizidae | *Zonotrichia albicollis* | White-throated sparrow |  | SRR1796638,  SRR1796642 |
|  | Fringillidae | *Serinus canaria* | Common canary |  | SRR2895902 |
|  | Muscicapidae | *Ficedula albicollis* | Collared flycatcher |  | ERR637368 |
|  | Fringillidae | *Spinus spinus* | Eurasian siskin | Transcriptome | 8 |
| Pelecaniformes | Threskiornithidae | *Nipponia nippon* | Crested ibis | Assembly | 2 |
|  | Ardeidae | *Egretta garzetta* | Little egret | Assembly | 2 |
|  |  |  |  | Resequencing,  Transcriptome | This study |
|  | Ardeidae | *Butorides striata* | Striated heron | Resequencing,  Transcriptome | This study |
|  | Threskiornithidae | *Platalea minor* | Black-faced spoonbill | Resequencing,  Transcriptome | This study |
| Opisthocomiformes | Opisthocomidae | *Opisthocomus hoazin* | Hoatzin | Assembly | 2 |
| Charadriiformes | Charadriidae | *Charadrius vociferus* | Killdeer | Assembly | 2 |
| Cuculiformes | Cuculidae | *Cuculus canorus* | Common cuckoo | Assembly | 2 |
| Caprimulgiformes | Caprimulgidae | *Antrostomus carolinensis* | Chuck-will’s-widow | Assembly  (low quality) | 2 |
| Apodiformes | Trochilidae | *Calypte anna* | Anna’s hummingbird | Assembly | 2 |
| Columbiformes | Columbidae | *Columba livia* | Rock dove | Assembly | 9 |
| Galliformes | Phasianidae | *Gallus gallus* | Chicken | Assembly | 10 |
| Struthioniformes | Struthionidae | *Struthio camelus* | Common ostrich | Assembly | 2 |
| Apterygiformes | Apterygidae | *Apteryx australis* | Brown kiwi | Assembly | 11 |

**Table S3. Sampling information on bird species sequenced in this study.** All samples were acquired from South Korea. Genomic DNA of each sample was deposited at the Wildlife Genetic Resources Bank of National Institute of Biological Resources.

| Scientific name | Collection number | Sampling date | Sample type^1^ | Sampling site^2^ | Accession number | Conservation Status^3^ | Permission number^4^ |
| --- | --- | --- | --- | --- | --- | --- | --- |
| **Strigiformes** |  |  |  |  |  |  |  |
| *Bubo bubo* | CN14-281 | Jun. 17, 2014 | Blood | Dangjin, CN | NIBRGR0000424809 | E(II), NM | CHA20150724 |
| *Otus sunia* | CN15-715 | Sep. 10, 2015 | Blood | Cheonan, CN | NIBRGR0000424814 |  |  |
| *Strix nivicolum* | CN14-119 | Jun. 17, 2014 | Blood | Cheongyang, CN | NIBRGR0000424811 | E(II), NM | CHA 20150724 |
| *Ninox japonica* | CN14-387 | Jun. 25, 2014 | Blood | Dangjin, CN | NIBRGR0000424812 | NM | CHA 20150724 |
| *Asio otus* | CN14-021 | Aug. 5, 2014 | Blood | Asan, CN | NIBRGR0000424813 | NM | CHA 20150724 |
| *Asio flammeus* | IN1668 | Feb. 6, 2017 | Blood* | Cheongju, CB | NIBRGR0000424831 | NM | Not necessary |
| *Otus semitorques* | CN17-040 | Feb. 15, 2017 | Blood* | Asan, CN | NIBRGR0000424811 | NM | CHA 20170215 |
| **Accipitriformes** |  |  |  |  |  |  |  |
| *Buteo japonicus* | CN15-030 | Jan. 26, 2015 | Blood | Gwangju | NIBRGR0000424805 |  |  |
| *Accipiter nisus* | CN15-041 | Feb. 5, 2015 | Blood | Cheonan, CN | NIBRGR0000424806 | E(II), NM | CHA 20150724 |
| *Accipiter gentilis* | CN14-679 | Nov. 13, 2014 | Blood | Nonsan, CN | NIBRGR0000424819 | E(II), NM | CHA 20150724 |
| *Haliaeetus albicilla* | CN16-053 | Feb. 7, 2017 | Blood | Seosan, CN | NIBRGR0000424828 | E(I), NM | MOE 2017-03 |
| *Pernis ptilorhynchus* | CN11-616 | Jan. 5, 2017 | Blood* | Seosan, CN | NIBRGR0000424829 | E(II) | MOE 2017-01 |
| *Milvus migrans* | IN1594 | Jun. 1, 2017 | Muscle | Busan | NIBRGR0000424830 | E(II) | MOE 2017-18 |
| *Accipiter soloensis* | IN1803 | Jun. 12, 2017 | Muscle | Busan | NIBRGR0000424832 | NM | CHA 20170608 |
| **Falconiformes** |  |  |  |  |  |  |  |
| *Falco tinnunculus* | CN14-642 | Nov. 24, 2014 | Blood | Seosan, CN | NIBRGR0000424802 | NM | CHA 20150724 |
| *Falco subbuteo* | CN14-472 | Jul. 19, 2014 | Blood | Seocheon, CN | NIBRGR0000424810 | E(II) | MOE 2017-34 |
| **Others** |  |  |  |  |  |  |  |
| *Picus canus* | CN14-463 | Jul. 18, 2014 | Blood | Cheonan, CN | NIBRGR0000424815 |  |  |
| *Egretta garzetta* | CN14-545 | Aug. 14, 2014 | Blood | Nonsan, CN | NIBRGR0000424817 |  |  |
| *Butorides striata* | CN14-540 | Aug. 14, 2014 | Blood | Yesan, CN | NIBRGR0000424816 |  |  |
| *Platalea minor* | CN14-386 | Jun. 25, 2014 | Blood | Incheon | NIBRGR0000424818 | E(I), NM | CHA 20150724 |

1. Blood samples for *A*. *flammeus,* *O*. *semitorques*, and *P*. *ptilorhynchus* (denoted with *) were obtained from the live individuals during medical check-up at the wild animal rescue center or at the zoo.
2. Abbreviation for sampling sites are: CN, Chungcheongnam-do and CB, Chungcheongbuk-do
3. Conservation status: Endangered species of Korea (E) in tier I or tier II were listed by Ministry of Environment (MOE). Natural Monument (NM) were listed by Cultural Heritage Administration (CHA).
4. Permissions for sampling were obtained from MOE for the endangered species of Korea and from the CHA for the natural monuments, respectively. Permission for *A. flammeus*, a natural monument, was not necessary because blood samples were obtained during medical check-up at the zoo.

**Table S4. Sequencing library statistics used for the four bird of prey genome assemblies**

**a.** Eurasian eagle-owl

| Insert size | Libraries | Number of reads | Read length (bp) | Total bases (bp) | Depth (×)  (Genome size: 1.2 Gb) | |
| --- | --- | --- | --- | --- | --- | --- |
| 170bp | L1_1 | 249,291,256 | 101 | 25,178,416,856 | 20.98 | 41.96 |
|  | L1_2 | 249,291,256 | 101 | 25,178,416,856 | 20.98 |  |
| 500bp | L1_1 | 213,203,268 | 101 | 21,533,530,068 | 17.94 | 35.89 |
|  | L1_2 | 213,203,268 | 101 | 21,533,530,068 | 17.94 |  |
| 700bp | L1_1 | 240,640,668 | 101 | 24,304,707,468 | 20.25 | 40.51 |
|  | L1_2 | 240,640,668 | 101 | 24,304,707,468 | 20.25 |  |
| 2 Kb | L1_1 | 144,374,940 | 51 | 7,363,121,940 | 6.14 | 24.60 |
|  | L1_2 | 144,374,940 | 51 | 7,363,121,940 | 6.14 |  |
|  | L2_1 | 145,023,351 | 51 | 7,396,190,901 | 6.16 |  |
|  | L2_2 | 145,023,351 | 51 | 7,396,190,901 | 6.16 |  |
| 5 Kb | L1_1 | 168,001,761 | 51 | 8,568,089,811 | 7.14 | 25.02 |
|  | L1_2 | 168,001,761 | 51 | 8,568,089,811 | 7.14 |  |
|  | L2_1 | 126,388,158 | 51 | 6,445,796,058 | 5.37 |  |
|  | L2_2 | 126,388,158 | 51 | 6,445,796,058 | 5.37 |  |
| 10 Kb | L1_1 | 135,923,860 | 51 | 6,932,116,860 | 5.78 | 23.22 |
|  | L1_2 | 135,923,860 | 51 | 6,932,116,860 | 5.78 |  |
|  | L2_1 | 137,270,089 | 51 | 7,000,774,539 | 5.83 |  |
|  | L2_2 | 137,270,089 | 51 | 7,000,774,539 | 5.83 |  |
| 15 Kb | L1_1 | 136,679,009 | 51 | 6,970,629,459 | 5.81 | 22.81 |
|  | L1_2 | 136,679,009 | 51 | 6,970,629,459 | 5.81 |  |
|  | L2_1 | 131,722,262 | 51 | 6,717,835,362 | 5.60 |  |
|  | L2_2 | 131,722,262 | 51 | 6,717,835,362 | 5.60 |  |
| Total | | 3,657,037,244 | - | 256,822,418,644 | 214.02 | |

**b.** Oriental scops-owl

| Insert size | Libraries | Number of reads | Read length (bp) | Total bases (bp) | Depth (×)  (Genome size: 1.2 Gb) | |
| --- | --- | --- | --- | --- | --- | --- |
| 170bp | L1_1 | 180,852,772 | 101 | 18,266,129,972 | 15.22 | 30.44 |
|  | L1_2 | 180,852,772 | 101 | 18,266,129,972 | 15.22 |  |
| 500bp | L1_1 | 254,984,477 | 101 | 25,753,432,177 | 21.46 | 42.92 |
|  | L1_2 | 254,984,477 | 101 | 25,753,432,177 | 21.46 |  |
| 700bp | L1_1 | 296,423,657 | 101 | 29,938,789,357 | 24.95 | 49.90 |
|  | L1_2 | 296,423,657 | 101 | 29,938,789,357 | 24.95 |  |
| 2 Kb | L1_1 | 125,243,856 | 51 | 6,387,436,656 | 5.32 | 10.65 |
|  | L1_2 | 125,243,856 | 51 | 6,387,436,656 | 5.32 |  |
|  | L2_1 | 118,398,262 | 51 | 6,038,311,362 | 5.03 |  |
|  | L2_2 | 118,398,262 | 51 | 6,038,311,362 | 5.03 |  |
| 5 Kb | L1_1 | 113,197,829 | 51 | 5,773,089,279 | 4.81 | 21.69 |
|  | L1_2 | 113,197,829 | 51 | 5,773,089,279 | 4.81 |  |
|  | L2_1 | 141,928,164 | 51 | 7,238,336,364 | 6.03 |  |
|  | L2_2 | 141,928,164 | 51 | 7,238,336,364 | 6.03 |  |
| 10 Kb | L1_1 | 145,025,441 | 51 | 7,396,297,491 | 6.16 | 25.79 |
|  | L1_2 | 145,025,441 | 51 | 7,396,297,491 | 6.16 |  |
|  | L2_1 | 158,432,357 | 51 | 8,080,050,207 | 6.73 |  |
|  | L2_2 | 158,432,357 | 51 | 8,080,050,207 | 6.73 |  |
| 15 Kb | L1_1 | 125,577,776 | 51 | 6,404,466,576 | 5.34 | 22.57 |
|  | L1_2 | 125,577,776 | 51 | 6,404,466,576 | 5.34 |  |
|  | L2_1 | 139,895,616 | 51 | 7,134,676,416 | 5.95 |  |
|  | L2_2 | 139,895,616 | 51 | 7,134,676,416 | 5.95 |  |
| Total | | 3,599,920,414 | - | 256,822,031,714 | 214.02 | |

**c.** Eastern buzzard

| Insert size | Libraries | Number of reads | Read length (bp) | Total bases (bp) | Depth (×)  (Genome size: 1.2 Gb) | |
| --- | --- | --- | --- | --- | --- | --- |
| 170bp | L1_1 | 185,324,078 | 101 | 18,717,731,878 | 15.60 | 31.20 |
|  | L1_2 | 185,324,078 | 101 | 18,717,731,878 | 15.60 |  |
| 500bp | L1_1 | 191,710,339 | 101 | 19,362,744,239 | 16.14 | 32.27 |
|  | L1_2 | 191,710,339 | 101 | 19,362,744,239 | 16.14 |  |
| 700bp | L1_1 | 196,469,467 | 101 | 19,843,416,167 | 16.54 | 33.07 |
|  | L1_2 | 196,469,467 | 101 | 19,843,416,167 | 16.54 |  |
| 2 Kb | L1_1 | 132,171,642 | 51 | 6,740,753,742 | 5.62 | 20.94 |
|  | L1_2 | 132,171,642 | 51 | 6,740,753,742 | 5.62 |  |
|  | L2_1 | 114,155,584 | 51 | 5,821,934,784 | 4.85 |  |
|  | L2_2 | 114,155,584 | 51 | 5,821,934,784 | 4.85 |  |
| 5 Kb | L1_1 | 142,102,000 | 51 | 7,247,202,000 | 6.04 | 21.15 |
|  | L1_2 | 142,102,000 | 51 | 7,247,202,000 | 6.04 |  |
|  | L2_1 | 106,748,138 | 51 | 5,444,155,038 | 4.54 |  |
|  | L2_2 | 106,748,138 | 51 | 5,444,155,038 | 4.54 |  |
| 10 Kb | L1_1 | 120,979,918 | 51 | 6,169,975,818 | 5.14 | 22.52 |
|  | L1_2 | 120,979,918 | 51 | 6,169,975,818 | 5.14 |  |
|  | L2_1 | 143,946,999 | 51 | 7,341,296,949 | 6.12 |  |
|  | L2_2 | 143,946,999 | 51 | 7,341,296,949 | 6.12 |  |
| 15 Kb | L1_1 | 128,477,416 | 51 | 6,552,348,216 | 5.46 | 24.28 |
|  | L1_2 | 128,477,416 | 51 | 6,552,348,216 | 5.46 |  |
|  | L2_1 | 157,147,833 | 51 | 8,014,539,483 | 6.68 |  |
|  | L2_2 | 157,147,833 | 51 | 8,014,539,483 | 6.68 |  |
| Total | | 3,238,466,828 | - | 222,512,196,628 | 185.43 | |

**d.** Common kestrel

| Insert size | Libraries | Number of reads | Read length (bp) | Total bases (bp) | Depth (×)  (Genome size: 1.2 Gb) | |
| --- | --- | --- | --- | --- | --- | --- |
| 350bp | L1_1 | 256,002,867 | 101 | 25,856,289,567 | 21.55 | 43.09 |
|  | L1_2 | 256,002,867 | 101 | 25,856,289,567 | 21.55 |  |
| 550bp | L2_1 | 265,998,079 | 101 | 26,865,805,979 | 22.39 | 44.78 |
|  | L2_2 | 265,998,079 | 101 | 26,865,805,979 | 22.39 |  |
| 2 Kb | L1_1 | 252,588,739 | 101 | 25,511,462,639 | 21.26 | 125.53 |
|  | L1_2 | 252,588,739 | 101 | 25,511,462,639 | 21.26 |  |
|  | L2_1 | 257,523,464 | 101 | 26,009,869,864 | 21.67 |  |
|  | L2_2 | 257,523,464 | 101 | 26,009,869,864 | 21.67 |  |
|  | L3_1 | 235,638,354 | 101 | 23,799,473,754 | 19.83 |  |
|  | L3_2 | 235,638,354 | 101 | 23,799,473,754 | 19.83 |  |
| 5 Kb | L1_1 | 266,270,102 | 101 | 26,893,280,302 | 22.41 | 90.39 |
|  | L1_2 | 266,270,102 | 101 | 26,893,280,302 | 22.41 |  |
|  | L2_1 | 270,713,864 | 101 | 27,342,100,264 | 22.79 |  |
|  | L2_2 | 270,713,864 | 101 | 27,342,100,264 | 22.79 |  |
| 10 Kb | L1_1 | 264,016,247 | 101 | 26,665,640,947 | 22.22 | 92.75 |
|  | L1_2 | 264,016,247 | 101 | 26,665,640,947 | 22.22 |  |
|  | L2_1 | 286,961,420 | 101 | 28,983,103,420 | 24.15 |  |
|  | L2_2 | 286,961,420 | 101 | 28,983,103,420 | 24.15 |  |
| 15 Kb | L1_1 | 291,186,433 | 101 | 29,409,829,733 | 24.51 | 97.88 |
|  | L1_2 | 291,186,433 | 101 | 29,409,829,733 | 24.51 |  |
|  | L2_1 | 290,303,234 | 101 | 29,320,626,634 | 24.43 |  |
|  | L2_2 | 290,303,234 | 101 | 29,320,626,634 | 24.43 |  |
| Total | | 5,874,405,606 | - | 593,314,966,206 | 494.43 | |

**Table S5. Filtered sequence information of the four birds of prey**

**a.** Eurasian eagle-owl

| Libraries | | Number of  raw reads | Number of  remained reads | Trimmed read  length (bp) | Remained total  bases (bp) | Remained  sequence depth (×) |
| --- | --- | --- | --- | --- | --- | --- |
| 170bp | L1_1 | 249,291,256 | 236,762,446 | 90 | 21,308,620,140 | 17.76 |
|  | L1_2 | 249,291,256 | 236,762,446 | 90 | 21,308,620,140 | 17.76 |
| 500bp | L1_1 | 213,203,268 | 196,010,740 | 90 | 17,640,966,600 | 14.70 |
|  | L1_2 | 213,203,268 | 196,010,740 | 90 | 17,640,966,600 | 14.70 |
| 700bp | L1_1 | 240,640,668 | 196,547,773 | 90 | 17,689,299,570 | 14.74 |
|  | L1_2 | 240,640,668 | 196,547,773 | 90 | 17,689,299,570 | 14.74 |
| 2 Kb | L1_1 | 144,374,940 | 14,584,273 | 50 | 729,213,650 | 0.61 |
|  | L1_2 | 144,374,940 | 14,584,273 | 50 | 729,213,650 | 0.61 |
|  | L2_1 | 145,023,351 | 20,366,074 | 50 | 1,018,303,700 | 0.85 |
|  | L2_2 | 145,023,351 | 20,366,074 | 50 | 1,018,303,700 | 0.85 |
| 5 Kb | L1_1 | 168,001,761 | 19,431,467 | 50 | 971,573,350 | 0.81 |
|  | L1_2 | 168,001,761 | 19,431,467 | 50 | 971,573,350 | 0.81 |
|  | L2_1 | 126,388,158 | 14,265,502 | 50 | 713,275,100 | 0.59 |
|  | L2_2 | 126,388,158 | 14,265,502 | 50 | 713,275,100 | 0.59 |
| 10 Kb | L1_1 | 135,923,860 | 11,653,024 | 50 | 582,651,200 | 0.49 |
|  | L1_2 | 135,923,860 | 11,653,024 | 50 | 582,651,200 | 0.49 |
|  | L2_1 | 137,270,089 | 11,107,921 | 50 | 555,396,050 | 0.46 |
|  | L2_2 | 137,270,089 | 11,107,921 | 50 | 555,396,050 | 0.46 |
| 15 Kb | L1_1 | 136,679,009 | 11,014,715 | 50 | 550,735,750 | 0.46 |
|  | L1_2 | 136,679,009 | 11,014,715 | 50 | 550,735,750 | 0.46 |
|  | L2_1 | 131,722,262 | 11,041,607 | 50 | 552,080,350 | 0.46 |
|  | L2_2 | 131,722,262 | 11,041,607 | 50 | 552,080,350 | 0.46 |
| Total | | 3,657,037,244 | 1,485,571,084 | - | 124,624,230,920 | 103.85 |

**b.** Oriental scops-owl

| Libraries | | Number of  raw reads | Number of remained reads | Trimmed read length (bp) | Remained total bases (bp) | Remained  sequence depth (×) |
| --- | --- | --- | --- | --- | --- | --- |
| 170bp | L1_1 | 180,852,772 | 175,287,324 | 90 | 15,775,859,160 | 13.15 |
|  | L1_2 | 180,852,772 | 175,287,324 | 90 | 15,775,859,160 | 13.15 |
| 500bp | L1_1 | 254,984,477 | 231,289,349 | 90 | 20,816,041,410 | 17.35 |
|  | L1_2 | 254,984,477 | 231,289,349 | 90 | 20,816,041,410 | 17.35 |
| 700bp | L1_1 | 296,423,657 | 281,922,971 | 90 | 25,373,067,390 | 21.14 |
|  | L1_2 | 296,423,657 | 281,922,971 | 90 | 25,373,067,390 | 21.14 |
| 2 Kb | L1_1 | 125,243,856 | 33,676,570 | 50 | 1,683,828,500 | 1.40 |
|  | L1_2 | 125,243,856 | 33,676,570 | 50 | 1,683,828,500 | 1.40 |
|  | L2_1 | 118,398,262 | 28,542,283 | 50 | 1,427,114,150 | 1.19 |
|  | L2_2 | 118,398,262 | 28,542,283 | 50 | 1,427,114,150 | 1.19 |
| 5 Kb | L1_1 | 113,197,829 | 16,117,739 | 50 | 805,886,950 | 0.67 |
|  | L1_2 | 113,197,829 | 16,117,739 | 50 | 805,886,950 | 0.67 |
|  | L2_1 | 141,928,164 | 18,751,681 | 50 | 937,584,050 | 0.78 |
|  | L2_2 | 141,928,164 | 18,751,681 | 50 | 937,584,050 | 0.78 |
| 10 Kb | L1_1 | 145,025,441 | 11,072,239 | 50 | 553,611,950 | 0.46 |
|  | L1_2 | 145,025,441 | 11,072,239 | 50 | 553,611,950 | 0.46 |
|  | L2_1 | 158,432,357 | 13,443,971 | 50 | 672,198,550 | 0.56 |
|  | L2_2 | 158,432,357 | 13,443,971 | 50 | 672,198,550 | 0.56 |
| 15 Kb | L1_1 | 125,577,776 | 8,202,795 | 50 | 410,139,750 | 0.34 |
|  | L1_2 | 125,577,776 | 8,202,795 | 50 | 410,139,750 | 0.34 |
|  | L2_1 | 139,895,616 | 10,677,382 | 50 | 533,869,100 | 0.44 |
|  | L2_2 | 139,895,616 | 10,677,382 | 50 | 533,869,100 | 0.44 |
| Total | | 3,599,920,414 | 1,657,968,608 | - | 137,978,401,920 | 114.98 |

**c.** Eastern buzzard

| Libraries | | Number of  raw reads | Number of  remained reads | Trimmed read  length (bp) | Remained total  bases (bp) | Remained  sequence depth (×) |
| --- | --- | --- | --- | --- | --- | --- |
| 170bp | L1_1 | 185,324,078 | 178,589,150 | 90 | 16,073,023,500 | 13.39 |
|  | L1_2 | 185,324,078 | 178,589,150 | 90 | 16,073,023,500 | 13.39 |
| 500bp | L1_1 | 191,710,339 | 178,043,957 | 90 | 16,023,956,130 | 13.35 |
|  | L1_2 | 191,710,339 | 178,043,957 | 90 | 16,023,956,130 | 13.35 |
| 700bp | L1_1 | 196,469,467 | 164,591,674 | 90 | 14,813,250,660 | 12.34 |
|  | L1_2 | 196,469,467 | 164,591,674 | 90 | 14,813,250,660 | 12.34 |
| 2 Kb | L1_1 | 132,171,642 | 21,346,869 | 50 | 1,067,343,450 | 0.89 |
|  | L1_2 | 132,171,642 | 21,346,869 | 50 | 1,067,343,450 | 0.89 |
|  | L2_1 | 114,155,584 | 42,036,718 | 50 | 2,101,835,900 | 1.75 |
|  | L2_2 | 114,155,584 | 42,036,718 | 50 | 2,101,835,900 | 1.75 |
| 5 Kb | L1_1 | 142,102,000 | 21,715,302 | 50 | 1,085,765,100 | 0.90 |
|  | L1_2 | 142,102,000 | 21,715,302 | 50 | 1,085,765,100 | 0.90 |
|  | L2_1 | 106,748,138 | 11,802,508 | 50 | 590,125,400 | 0.49 |
|  | L2_2 | 106,748,138 | 11,802,508 | 50 | 590,125,400 | 0.49 |
| 10 Kb | L1_1 | 120,979,918 | 11,519,217 | 50 | 575,960,850 | 0.48 |
|  | L1_2 | 120,979,918 | 11,519,217 | 50 | 575,960,850 | 0.48 |
|  | L2_1 | 143,946,999 | 13,612,955 | 50 | 680,647,750 | 0.57 |
|  | L2_2 | 143,946,999 | 13,612,955 | 50 | 680,647,750 | 0.57 |
| 15 Kb | L1_1 | 128,477,416 | 9,490,611 | 50 | 474,530,550 | 0.40 |
|  | L1_2 | 128,477,416 | 9,490,611 | 50 | 474,530,550 | 0.40 |
|  | L2_1 | 157,147,833 | 13,085,512 | 50 | 654,275,600 | 0.55 |
|  | L2_2 | 157,147,833 | 13,085,512 | 50 | 654,275,600 | 0.55 |
| Total | | 3,238,466,828 | 1,331,668,946 | - | 108,281,429,780 | 90.23 |

**d.** Common kestrel

| Libraries | | Number of  raw reads | Number of remained reads | Trimmed read length (bp) | Remained total bases (bp) | Remained  sequence depth (×) |
| --- | --- | --- | --- | --- | --- | --- |
| 350bp | L1_1 | 256,002,867 | 252,540,520 | 90 | 22,728,646,800 | 18.94 |
|  | L1_2 | 256,002,867 | 252,540,520 | 90 | 22,728,646,800 | 18.94 |
| 550bp | L1_1 | 265,998,079 | 261,449,755 | 90 | 23,530,477,950 | 19.61 |
|  | L1_2 | 265,998,079 | 261,449,755 | 90 | 23,530,477,950 | 19.61 |
| 2 Kb | L1_1 | 252,588,739 | 113,264,462 | 50 | 5,663,223,100 | 4.72 |
|  | L1_2 | 252,588,739 | 113,264,462 | 50 | 5,663,223,100 | 4.72 |
|  | L2_1 | 257,523,464 | 106,894,951 | 50 | 5,344,747,550 | 4.45 |
|  | L2_2 | 257,523,464 | 106,894,951 | 50 | 5,344,747,550 | 4.45 |
|  | L3_1 | 235,638,354 | 116,404,884 | 50 | 5,820,244,200 | 4.85 |
|  | L3_2 | 235,638,354 | 116,404,884 | 50 | 5,820,244,200 | 4.85 |
| 5 Kb | L1_1 | 266,270,102 | 82,940,503 | 50 | 4,147,025,150 | 3.46 |
|  | L1_2 | 266,270,102 | 82,940,503 | 50 | 4,147,025,150 | 3.46 |
|  | L2_1 | 270,713,864 | 88,939,720 | 50 | 4,446,986,000 | 3.71 |
|  | L2_2 | 270,713,864 | 88,939,720 | 50 | 4,446,986,000 | 3.71 |
| 10 Kb | L1_1 | 264,016,247 | 117,476,768 | 50 | 5,873,838,400 | 4.89 |
|  | L1_2 | 264,016,247 | 117,476,768 | 50 | 5,873,838,400 | 4.89 |
|  | L2_1 | 286,961,420 | 84,772,614 | 50 | 4,238,630,700 | 3.53 |
|  | L2_2 | 286,961,420 | 84,772,614 | 50 | 4,238,630,700 | 3.53 |
| 15 Kb | L1_1 | 291,186,433 | 73,705,167 | 50 | 3,685,258,350 | 3.07 |
|  | L1_2 | 291,186,433 | 73,705,167 | 50 | 3,685,258,350 | 3.07 |
|  | L2_1 | 290,303,234 | 70,574,972 | 50 | 3,528,748,600 | 2.94 |
|  | L2_2 | 290,303,234 | 70,574,972 | 50 | 3,528,748,600 | 2.94 |
| Total | | 5,874,405,606 | 2,737,928,632 | - | 178,015,653,600 | 148.35 |

**Table S6. 17-mer statistics information for 20 avian species**

| Species | *K*-mer total number | Peak depth | Estimated genome size (Gb) |
| --- | --- | --- | --- |
| Eurasian eagle-owl | 93,139,501,932 | 63 | 1.478 |
| Oriental scops-owl | 101,897,947,312 | 77 | 1.323 |
| Himalayan owl | 25,301,442,982 | 20 | 1.265 |
| Northern boobook | 25,148,256,432 | 20 | 1.257 |
| Long-eared owl | 29,586,880,640 | 24 | 1.233 |
| Short-eared owl | 23,353,960,560 | 14 | 1.668 |
| Japanese scops-owl | 27,992,952,456 | 22 | 1.272 |
| Eastern buzzard | 77,141,267,588 | 63 | 1.224 |
| Eurasian sparrowhawk | 29,072,816,384 | 21 | 1.384 |
| Northern goshawk | 30,522,477,568 | 21 | 1.453 |
| White-tailed eagle | 24,390,282,000 | 18 | 1.355 |
| Oriental honey-buzzard | 25,305,632,016 | 16 | 1.582 |
| Black kite | 25,712,659,392 | 19 | 1.353 |
| Chinese sparrowhawk | 22,366,586,088 | 15 | 1.491 |
| Common kestrel | 76,070,560,700 | 60 | 1.268 |
| Eurasian hobby | 30,173,856,256 | 23 | 1.312 |
| Grey-headed woodpecker | 13,036,606,884 | 10 | 1.304 |
| Little Egret | 11,702,977,902 | 10 | 1.170 |
| Striated heron | 11,750,011,056 | 10 | 1.175 |
| Black-faced spoonbill | 12,485,284,056 | 11 | 1.135 |

**Table S7. Global assembly statistics of the four bird of prey genomes.** Final assemblies were selected by assessing assembly statistics, transcripts mapping results, and single-copy orthologs mapping results. Oriental scops-owl genome could not be assembled when using SOAPdenovo2, probably because of its high level of heterozygosity.

**a.** Eurasian eagle-owl

| Assembler | **SOAPdenovo2 (selected)** | | Platanus | |
| --- | --- | --- | --- | --- |
| Assembly level | Contig | Scaffold | Contig | Scaffold |
| # of sequences | 133,374 | 35,672 | 119,288 | 72,452 |
| Total bases (bp) | 1,201,498,232 | 1,258,075,470 | 1,168,795,702 | 1,194,726,138 |
| Longest length (bp) | 391,237 | 29,915,079 | 374,073 | 33,945,943 |
| Shortest length (bp) | 55 | 200 | 1 | 200 |
| N50 (bp) | 37,088 | 8,019,613 | 46,324 | 6,858,015 |
| GC contents | 41.39 % | 41.39 % | 41.10 % | 41.10 % |
| N base ratio | 0.00 % | 4.50 % | 0.00 % | 2.17 % |

**b.** Oriental scops-owl

| Assembler | **Platanus (selected)** | |
| --- | --- | --- |
| Assembly level | Contig | Scaffold |
| # of sequences | 151,529 | 108,835 |
| Total bases (bp) | 1,212,043,918 | 1,231,452,760 |
| Longest length (bp) | 406,680 | 81,210,816 |
| Shortest length (bp) | 2 | 200 |
| N50 (bp) | 52,366 | 18,540,887 |
| GC contents | 41.70 % | 41.70 % |
| N base ratio | 0.00 % | 1.58 % |

**c.** Eastern buzzard

| Assembler | **SOAPdenovo2 (selected)** | | Platanus | |
| --- | --- | --- | --- | --- |
| Assembly level | Contig | Scaffold | Contig | Scaffold |
| # of sequences | 127,106 | 29,208 | 112,214 | 65,890 |
| Total bases (bp) | 1,216,111,145 | 1,259,752,395 | 1,178,354,735 | 1,203,691,678 |
| Longest length (bp) | 370,481 | 31,377,212 | 389,759 | 32,375,696 |
| Shortest length (bp) | 55 | 200 | 5 | 200 |
| N50 (bp) | 36,722 | 7,468,869 | 46,157 | 8,836,483 |
| GC contents | 41.66 % | 41.66 % | 41.38 % | 41.38 % |
| N base ratio | 0.00 % | 3.46 % | 0.00 % | 2.11 % |

**d.** Common kestrel

| Assembler | SOAPdenovo2 | | **Platanus (selected)** | |
| --- | --- | --- | --- | --- |
| Assembly level | Contig | Scaffold | Contig | Scaffold |
| # of sequences | 245,216 | 34,131 | 123,820 | 69,121 |
| Total bases (bp) | 1,342,122,283 | 1,430,330,502 | 1,173,072,102 | 1,179,916,286 |
| Longest length (bp) | 138,002 | 44,828,457 | 305,150 | 64,290,118 |
| Shortest length (bp) | 1 | 199 | 1 | 200 |
| N50 (bp) | 12,031 | 13,066,975 | 37,053 | 21,232,185 |
| GC contents | 42.20 % | 42.20 % | 42.11 % | 42.11 % |
| N base ratio | 0.00 % | 6.17 % | 0.00 % | 0.58 % |

**Table S8. Assessment of gene coverage by assembled bird of prey transcripts.** Final assemblies were selected by assessing assembly statistics, transcripts mapping results, and single-copy ortholog mapping results.

| Species | Assembler | Transcript length | Number | Total length (bp) | Sequence covered by Assembly (%) | With >90% sequence in one scaffold | | With >50% sequence in one scaffold | |
| --- | --- | --- | --- | --- | --- | --- | --- | --- | --- |
|  |  |  |  |  |  | Number | Percent (%) | Number | Percent (%) |
| Eurasian eagle-owl | **SOAPdenovo2**  **(selected)** | All | 192,557 | 247,620,874 | 94.20 | 164,428 | 85.39 | 175,583 | 91.18 |
|  |  | >200bp | 192,557 | 247,620,874 | 94.20 | 164,428 | 85.39 | 175,583 | 91.18 |
|  |  | >500bp | 98,755 | 216,127,215 | 95.15 | 85,725 | 86.81 | 93,076 | 94.25 |
|  |  | >1000bp | 58,927 | 188,526,444 | 95.84 | 51,799 | 87.90 | 56,745 | 96.30 |
|  | Platanus | All | 192,557 | 247,620,874 | 89.46 | 145,471 | 75.55 | 164,060 | 85.20 |
|  |  | >200bp | 192,557 | 247,620,874 | 89.46 | 145,471 | 75.55 | 164,060 | 85.20 |
|  |  | >500bp | 98,755 | 216,127,215 | 90.79 | 76,580 | 77.55 | 88,193 | 89.30 |
|  |  | >1000bp | 58,927 | 188,526,444 | 91.68 | 46,400 | 78.74 | 53,814 | 91.32 |
| Oriental scops-owl | **Platanus**  **(selected)** | All | 188,436 | 226,146,141 | 96.19 | 164,344 | 87.21 | 178,179 | 94.56 |
|  |  | >200bp | 188,436 | 226,146,141 | 96.19 | 164,344 | 87.21 | 178,179 | 94.56 |
|  |  | >500bp | 94,884 | 195,260,793 | 97.03 | 85,930 | 90.56 | 82,093 | 97.06 |
|  |  | >1000bp | 54,707 | 167,470,729 | 97.52 | 50,922 | 93.08 | 53,697 | 98.15 |
| Eastern buzzard | **SOAPdenovo2**  **(selected)** | All | 177,945 | 231,089,047 | 97.84 | 166,819 | 93.75 | 175,078 | 98.39 |
|  |  | >200bp | 177,945 | 231,089,047 | 97.84 | 166,819 | 93.75 | 175,078 | 98.39 |
|  |  | >500bp | 95,936 | 203,493,171 | 97.96 | 89,978 | 93.79 | 94,883 | 98.90 |
|  |  | >1000bp | 56,968 | 176,443,974 | 98.07 | 53,696 | 94.26 | 56,545 | 99.26 |
|  | Platanus | All | 177,945 | 231,089,047 | 94.28 | 150,231 | 84.43 | 166,059 | 93.32 |
|  |  | >200bp | 177,945 | 231,089,047 | 94.28 | 150,231 | 84.43 | 166,059 | 93.32 |
|  |  | >500bp | 95,936 | 203,493,171 | 94.82 | 81,824 | 85.29 | 91,211 | 95.07 |
|  |  | >1000bp | 56,968 | 176,443,974 | 95.21 | 49,053 | 86.11 | 54,562 | 95.78 |
| Common kestrel | SOAPdenovo2 | All | 181,982 | 226,109,028 | 99.26 | 177,597 | 97.59 | 181,297 | 99.62 |
|  |  | >200bp | 181,982 | 226,109,028 | 99.26 | 177,597 | 97.59 | 181,297 | 99.62 |
|  |  | >500bp | 97,087 | 197,379,462 | 99.32 | 95,323 | 98.18 | 96,839 | 99.74 |
|  |  | >1000bp | 57,126 | 169,661,898 | 99.36 | 56,200 | 98.38 | 57,023 | 99.82 |
|  | **Platanus**  **(selected)** | All | 181,982 | 226,109,028 | 97.70 | 171,187 | 94.07 | 178,926 | 98.32 |
|  |  | >200bp | 181,982 | 226,109,028 | 97.70 | 171,187 | 94.07 | 178,926 | 98.32 |
|  |  | >500bp | 97,087 | 197,379,462 | 97.73 | 91,572 | 94.32 | 95,139 | 97.99 |
|  |  | >1000bp | 57,126 | 169,661,898 | 97.85 | 54,237 | 94.94 | 56,027 | 98.08 |

**Table S9. Evaluation of the completeness of bird of prey assemblies and gene sets using single-copy orthologs mapping approach.** Final assemblies were selected by assessing assembly statistics, transcripts mapping results, and single-copy ortholog mapping results. For the common kestrel, only the Platanus assembly was tested, since the N50 lengths of contig and scaffold were much longer than those of SOAPdenovo2 assembly.

| Species | Assembler | Number of predicted genes | Complete  (%) | Duplicated  (%) | Fragment  (%) | Missing  (%) | Number of single-copy  orthologs genes |
| --- | --- | --- | --- | --- | --- | --- | --- |
| Eurasian eagle-owl | **SOAPdenovo2**  **(selected)** | 16,897 | 92.09% | 1.02% | 5.96% | 1.95% | 4,915 |
|  | Platanus | 15,632 | 91.27% | 1.00% | 6.57% | 2.16% | 4,915 |
| Oriental scops-owl | **Platanus**  **(selected)** | 17,710 | 91.35% | 1.44% | 6.37% | 2.28% | 4,915 |
| Eastern buzzard | **SOAPdenovo2**  **(selected)** | 17,376 | 96.72% | 1.18% | 2.38% | 0.90% | 4,915 |
|  | Platanus | 16,047 | 96.38% | 1.04% | 2.67% | 0.96% | 4,915 |
| Common kestrel | **Platanus**  **(selected)** | 16,481 | 93.90% | 1.22% | 4.46% | 1.65% | 4,915 |

**Table S10. Protein-coding gene prediction statistics for the four bird of prey genomes**

**a.** Eurasian eagle-owl

| Gene set | | Number | Average transcript  length (bp) | Average CDS length (bp) | Average no. of exons per gene | Average exon length (bp) | Average  intron length (bp) |
| --- | --- | --- | --- | --- | --- | --- | --- |
| *de novo* | AUGUSTUS | 27,448 | 15,625.3 | 1,605.8 | 9.0 | 178.5 | 3,441.4 |
| Homolog | Bald eagle | 13,546 | 23,302.6 | 1,598.9 | 9.3 | 171.4 | 2,605.4 |
|  | Barn owl | 14,897 | 15,380.9 | 1,270.6 | 7.6 | 167.5 | 2,141.8 |
|  | Brown kiwi | 14,218 | 20,732.4 | 1,434.5 | 8.5 | 169.5 | 2,585.9 |
|  | Chicken | 14,105 | 22,318.1 | 1,512.0 | 8.9 | 170.6 | 2,646.5 |
|  | Common cuckoo | 13,366 | 22,770.5 | 1,545.2 | 9.1 | 169.0 | 2,607.3 |
|  | Crested ibis | 13,173 | 22,868.5 | 1,539.9 | 9.2 | 166.9 | 2,592.2 |
|  | Downy woodpecker | 13,054 | 22,747.4 | 1,510.0 | 9.0 | 168.2 | 2,662.2 |
|  | Golden eagle | 13,939 | 22,994.4 | 1,598.0 | 9.3 | 171.6 | 2,574.6 |
|  | Peregrine falcon | 13,059 | 23,252.2 | 1,566.2 | 9.3 | 168.6 | 2,616.4 |
|  | Zebra finch | 13,941 | 20,970.8 | 1,427.0 | 8.3 | 171.2 | 2,664.7 |
| Final | | 16,897 | 18,578.1 | 1,335.5 | 8.1 | 164.6 | 2,569.9 |

**b.** Oriental scops-owl

| Gene set | | Number | Average transcript  length (bp) | Average CDS length (bp) | Average no. of exons per gene | Average exon length (bp) | Average  intron length (bp) |
| --- | --- | --- | --- | --- | --- | --- | --- |
| *de novo* | AUGUSTUS | 26,719 | 15,018.1 | 1,583.3 | 9.4 | 168.6 | 3,190.8 |
| Homolog | Bald eagle | 13,736 | 23,055.6 | 1,651.1 | 9.7 | 170.3 | 2,462.1 |
|  | Barn owl | 14,707 | 15,171.4 | 1,280.4 | 7.6 | 168.1 | 2,099.9 |
|  | Brown kiwi | 14,492 | 19,803.5 | 1,435.5 | 8.5 | 169.9 | 2,466.3 |
|  | Chicken | 14,445 | 22,156.5 | 1,561.3 | 9.2 | 170.4 | 2,523.4 |
|  | Common cuckoo | 13,791 | 21,929.5 | 1,569.9 | 9.2 | 169.9 | 2,471.5 |
|  | Crested ibis | 13,397 | 22,349.9 | 1,584.0 | 9.5 | 167.5 | 2,455.3 |
|  | Downy woodpecker | 13,270 | 22,134.3 | 1,547.6 | 9.2 | 168.3 | 2,511.5 |
|  | Golden eagle | 14,259 | 22,772.8 | 1,656.2 | 9.7 | 171.4 | 2,437.9 |
|  | Peregrine falcon | 13,298 | 22,601.0 | 1,607.8 | 9.6 | 168.4 | 2,456.2 |
|  | Zebra finch | 13,756 | 21,011.2 | 1,459.0 | 8.6 | 168.9 | 2,559.8 |
| Final | | 17,710 | 17,497.8 | 1,349.1 | 8.2 | 165.2 | 2,402.5 |

**c.** Eastern buzzard

| Gene set | | Number | Average transcript  length (bp) | Average CDS length (bp) | Average no. of exons per gene | Average exon length (bp) | Average  intron length (bp) |
| --- | --- | --- | --- | --- | --- | --- | --- |
| *de novo* | AUGUSTUS | 23,535 | 17,609.1 | 1,817.1 | 10.9 | 167.4 | 3,193.1 |
| Homolog | Bald eagle | 14,928 | 22,831.0 | 1,613.9 | 9.5 | 170.6 | 2,508.0 |
|  | Barn owl | 15,423 | 15,000.1 | 1,247.6 | 7.5 | 167.0 | 2,125.6 |
|  | Budgerigar | 13,699 | 22,816.1 | 1,540.9 | 9.1 | 168.8 | 2,617.6 |
|  | Chicken | 14,635 | 21,995.1 | 1,526.5 | 9.0 | 169.4 | 2,554.9 |
|  | Common cuckoo | 13,680 | 22,806.9 | 1,558.7 | 9.4 | 166.6 | 2,542.6 |
|  | Crested ibis | 13,605 | 22,829.6 | 1,557.0 | 9.4 | 166.2 | 2,541.1 |
|  | Downy woodpecker | 13,321 | 22,874.1 | 1,524.1 | 9.2 | 165.9 | 2,607.0 |
|  | Golden Eagle | 15,039 | 22,920.9 | 1,625.7 | 9.6 | 169.9 | 2,485.0 |
|  | Peregrine falcon | 13,778 | 22,611.3 | 1,565.7 | 9.3 | 168.6 | 2,539.8 |
|  | Zebra finch | 14,209 | 20,810.9 | 1,433.1 | 8.4 | 170.2 | 2,610.7 |
| Final | | 17,376 | 21,776.7 | 1,509.1 | 9.1 | 165.9 | 2,654.5 |

**d.** Common kestrel

| Gene set | | Number | Average transcript  length (bp) | Average CDS length (bp) | Average no. of exons per gene | Average exon length (bp) | Average intron length (bp) |
| --- | --- | --- | --- | --- | --- | --- | --- |
| *de novo* | AUGUSTUS | 20,277 | 19,782.4 | 1,938.0 | 12.1 | 160.5 | 3,222.7 |
| Homolog | Bald eagle | 13,339 | 23,913.7 | 1,688.2 | 10.0 | 168.9 | 2,470.4 |
|  | Barn owl | 14,132 | 15,516.0 | 1,298.8 | 7.8 | 165.9 | 2,081.5 |
|  | Budgerigar | 13,084 | 23,564.1 | 1,594.0 | 9.5 | 167.9 | 2,586.7 |
|  | Chicken | 13,521 | 24,248.2 | 1,653.8 | 9.8 | 168.8 | 2,568.9 |
|  | Common cuckoo | 13,200 | 22,945.3 | 1,610.3 | 9.7 | 166.4 | 2,458.1 |
|  | Crested ibis | 13,035 | 22,877.4 | 1,605.6 | 9.6 | 167.1 | 2,470.5 |
|  | Downy woodpecker | 12,779 | 23,058.3 | 1,573.3 | 9.5 | 165.4 | 2,524.4 |
|  | Golden Eagle | 13,598 | 23,869.9 | 1,690.9 | 10.0 | 168.4 | 2,453.7 |
|  | Peregrine falcon | 13,838 | 23,504.8 | 1,652.8 | 9.9 | 166.5 | 2,447.3 |
|  | Zebra finch | 13,088 | 22,327.8 | 1,511.0 | 9.1 | 166.4 | 2,576.5 |
| Final | | 16,481 | 22,122.1 | 1,553.8 | 9.4 | 164.6 | 2,584.6 |

**Table S11. Transposable element statistics for the four bird of prey genomes**

**a.** Eurasian eagle-owl

| Type | *Ab initio*  based (bp) | Homology  based (bp) | Total (bp) | Percentage of  genome (%) |
| --- | --- | --- | --- | --- |
| DNA | 5,321,830 | 4,528,204 | 6,016,724 | 0.48% |
| LINE | 59,572,100 | 52,388,102 | 59,874,266 | 4.76% |
| LTR | 17,695,841 | 14,253,031 | 17,788,511 | 1.41% |
| Low_complexity | 1,957,049 | 2,017,594 | 2,025,077 | 0.16% |
| Retroposon | 1,256 | - | 1,256 | 0.00% |
| SINE | 969,396 | 1,064,867 | 1,198,180 | 0.10% |
| Satellite | 442,417 | 151,797 | 468,838 | 0.04% |
| Simple_repeat | 9,072,263 | 10,318,708 | 10,336,793 | 0.82% |
| TandemRepeat | - | - | 21,521,428 | 1.71% |
| Unknown | 771,686 | 740,728 | 867,230 | 0.07% |
| Unspecified | 2,767,143 | - | 2,767,143 | 0.22% |
| Total_TE | 97,746,927 | 85,408,455 | 114,298,851 | 9.09% |

**b.** Oriental scops-owl

| Type | *Ab initio*  based (bp) | Homology  based (bp) | Total (bp) | Percentage of  genome (%) |
| --- | --- | --- | --- | --- |
| DNA | 5,263,514 | 4,327,304 | 5,927,333 | 0.48% |
| LINE | 56,913,491 | 49,993,098 | 57,258,859 | 4.65% |
| LTR | 19,759,573 | 15,127,553 | 19,852,639 | 1.61% |
| Low_complexity | 2,243,449 | 111,234 | 2,275,506 | 0.18% |
| Retroposon | 2,276 | - | 2,276 | 0.00% |
| SINE | 913,404 | 1,018,313 | 1,136,775 | 0.09% |
| Satellite | 548,057 | 271,720 | 596,703 | 0.05% |
| Simple_repeat | 9,177,656 | 1,474,546 | 9,732,434 | 0.79% |
| TandemRepeat | - | - | 13,019,635 | 1.06% |
| Unknown | 759,745 | 725,087 | 847,083 | 0.07% |
| Unspecified | 2,469,624 | - | 2,469,624 | 0.20% |
| Total_TE | 97,219,820 | 72,984,692 | 107,200,077 | 8.71% |

**c.** Eastern buzzard

| Type | *Ab initio*  based (bp) | Homology  based (bp) | Total (bp) | Percentage of  genome (%) |
| --- | --- | --- | --- | --- |
| DNA | 5,603,025 | 5,007,155 | 6,396,464 | 0.51% |
| LINE | 41,381,034 | 33,893,267 | 41,735,462 | 3.31% |
| LTR | 31,511,623 | 27,354,821 | 31,623,377 | 2.51% |
| Low_complexity | 2,133,269 | 2,190,607 | 2,197,521 | 0.17% |
| Retroposon | 1,156 | - | 1,156 | 0.00% |
| SINE | 1,060,451 | 1,185,014 | 1,319,029 | 0.10% |
| Satellite | 431,142 | 198,737 | 458,150 | 0.04% |
| Simple_repeat | 8,292,936 | 9,347,676 | 9,366,893 | 0.74% |
| TandemRepeat | - | - | 18,908,531 | 1.50% |
| Unknown | 813,988 | 783,932 | 917,221 | 0.07% |
| Unspecified | 2,408,856 | - | 2,408,856 | 0.19% |
| Total_TE | 92,877,476 | 79,864,568 | 107,687,623 | 8.55% |

**d.** Common kestrel

| Type | *Ab initio*  based (bp) | Homology  based (bp) | Total (bp) | Percentage of  genome (%) |
| --- | --- | --- | --- | --- |
| DNA | 4,257,324 | 3,739,434 | 5,142,337 | 0.44% |
| LINE | 43,063,180 | 37,529,708 | 43,510,124 | 3.69% |
| LTR | 15,362,220 | 12,819,987 | 15,428,962 | 1.31% |
| Low_complexity | 2,518,248 | 2,583,944 | 2,591,422 | 0.22% |
| Retroposon | 881 | - | 881 | 0.00% |
| SINE | 662,953 | 883,172 | 955,565 | 0.08% |
| Satellite | 207,775 | 102,354 | 232,366 | 0.02% |
| Simple_repeat | 9,804,775 | 10,734,936 | 10,751,267 | 0.91% |
| TandemRepeat | - | - | 8,284,031 | 0.70% |
| Unknown | 614,646 | 657,964 | 735,905 | 0.06% |
| Unspecified | 1,645,735 | - | 1,645,735 | 0.14% |
| Total_TE | 77,520,599 | 68,999,966 | 83,921,845 | 7.11% |

**Table S12. Whole genome sequencing and mapping statistics for the birds of prey and non-raptor birds**

| Species | Reference assembly | The number of sequence reads | The number of  mapped reads | Mapping rate (%) | Estimated sequencing depth from mapped reads (×) |
| --- | --- | --- | --- | --- | --- |
| Eurasian eagle-owl | Eurasian eagle-owl | 426,406,536 | 423,883,463 | 99.41 | 35.63 |
| Oriental scops-owl | Oriental scops-owl | 509,968,954 | 497,671,220 | 97.59 | 41.47 |
|  | Eurasian eagle-owl | 509,968,954 | 484,464,677 | 95.00 | 40.72 |
| Himalayan owl | Eurasian eagle-owl | 475,120,796 | 463,937,275 | 97.65 | 38.79 |
| Northern spotted owl | Northern spotted owl | 342,525,114 | 329,308,129 | 96.14 | 39.93 |
|  | Eurasian eagle-owl | 342,525,114 | 322,365,208 | 94.11 | 40.25 |
| Barred owl | Northern spotted owl | 254,293,316 | 253,085,927 | 99.53 | 30.69 |
|  | Eurasian eagle-owl | 254,293,316 | 249,849,100 | 98.25 | 31.19 |
| Northern boobook | Eurasian eagle-owl | 483,070,632 | 456,180,964 | 94.43 | 38.13 |
| Long-eared owl | Eurasian eagle-owl | 534,446,094 | 517,804,343 | 96.89 | 43.32 |
| Short-eared owl | Eurasian eagle-owl | 298,420,942 | 292,213,676 | 97.92 | 36.72 |
| Japanese scops-owl | Eurasian eagle-owl | 344,542,548 | 338,549,210 | 98.26 | 42.55 |
| Barn owl | Eurasian eagle-owl | 315,670,102 | 250,294,698 | 79.29 | 20.83 |
| Eastern buzzard | Eastern buzzard | 383,420,678 | 381,454,295 | 99.49 | 31.68 |
| Bald eagle | Bald eagle | 490,942,426 | 479,287,574 | 97.63 | 41.34 |
|  | Eastern buzzard | 490,942,426 | 482,644,359 | 98.31 | 39.69 |
| Golden eagle | Golden eagle | 408,914,924 | 403,943,772 | 98.78 | 34.23 |
|  | Eastern buzzard | 408,914,924 | 397,378,332 | 97.18 | 32.68 |
| Eurasian sparrowhawk | Eastern buzzard | 504,924,004 | 485,432,318 | 96.14 | 40.32 |
| Northern goshawk | Eastern buzzard | 508,628,750 | 491,544,168 | 96.64 | 40.82 |
| Cinereous vulture | Eastern buzzard | 472,000,000 | 463,442,066 | 98.19 | 38.11 |
| White-tailed eagle | Eastern buzzard | 295,742,950 | 292,462,759 | 98.89 | 36.31 |
| Oriental honey-buzzard | Eastern buzzard | 319,662,296 | 308,671,755 | 96.56 | 38.33 |
| Black kite | Eastern buzzard | 325,611,358 | 319,810,066 | 98.22 | 39.71 |
| Chinese sparrowhawk | Eastern buzzard | 267,398,406 | 262,592,596 | 98.20 | 32.61 |
| Turkey vulture | Eastern buzzard | 372,261,628 | 301,706,530 | 81.05 | 24.81 |
| Common kestrel | Common kestrel | 531,996,158 | 518,208,664 | 97.41 | 44.62 |
| Saker falcon | Saker falcon | 485,798,050 | 479,708,329 | 98.75 | 41.68 |
|  | Common kestrel | 485,798,050 | 472,692,978 | 97.30 | 40.30 |
| Peregrine falcon | Peregrine falcon | 494,055,728 | 490,597,776 | 99.30 | 42.53 |
|  | Common kestrel | 494,055,728 | 484,437,741 | 98.05 | 41.30 |
| Eurasian hobby | Common kestrel | 500,017,118 | 485,539,341 | 97.10 | 41.80 |
| American crow | Zebra finch | 738,841,632 | 527,764,698 | 71.43 | 43.16 |
| Medium ground-finch | Zebra finch | 448,092,142 | 356,172,483 | 79.49 | 29.13 |
| White-throated sparrow | Zebra finch | 356,989,660 | 296,542,972 | 83.07 | 24.25 |
| Common canary | Zebra finch | 357,144,420 | 261,161,690 | 73.12 | 21.57 |
| Collared flycatcher | Zebra finch | 372,990,014 | 303,709,377 | 81.43 | 24.84 |

**Table S13. Variant statistics for the birds of prey and non-raptor birds**

| Species | Reference assembly | All variant sites | Homozygous SNV sites | Heterozygous SNV sites | Indel sites |
| --- | --- | --- | --- | --- | --- |
| Eurasian eagle-owl | Eurasian eagle-owl | 2,007,402 | 27,807 | 1,808,718 | 170,877 |
| Oriental scops-owl | Oriental scops-owl | 8,644,348 | 19,715 | 7,880,483 | 744,150 |
|  | Eurasian eagle-owl | 45,212,978 | 34,196,869 | 8,871,144 | 2,144,965 |
| Himalayan owl | Eurasian eagle-owl | 30,233,090 | 27,005,757 | 1,908,218 | 1,319,115 |
| Northern spotted owl | Northern spotted owl | 380,227 | 55,064 | 255,118 | 70,045 |
|  | Eurasian eagle-owl | 29,463,889 | 25,445,792 | 1,015,331 | 3,002,766 |
| Barred owl | Northern spotted owl | 11,863,926 | 7,598,827 | 3,089,938 | 1,175,161 |
|  | Eurasian eagle-owl | 30,121,988 | 23,604,477 | 3,413,634 | 3,103,877 |
| Northern boobook | Eurasian eagle-owl | 52,462,194 | 47,733,068 | 2,518,712 | 2,210,414 |
| Long-eared owl | Eurasian eagle-owl | 35,670,717 | 30,317,154 | 3,709,009 | 1,644,554 |
| Short-eared owl | Eurasian eagle-owl | 37,742,200 | 29,588,416 | 6,187,653 | 1,966,131 |
| Japanese scops-owl | Eurasian eagle-owl | 43,590,517 | 37,348,510 | 4,329,977 | 1,912,030 |
| Barn owl | Eurasian eagle-owl | 77,562,229 | 73,075,569 | 2,010,995 | 2,475,665 |
| Eastern buzzard | Eastern buzzard | 2,283,626 | 14,293 | 2,088,450 | 180,883 |
| Bald eagle | Bald eagle | 1,010,177 | 97,023 | 807,219 | 105,935 |
|  | Eastern buzzard | 20,542,155 | 18,490,544 | 630,353 | 1,421,258 |
| Golden eagle | Golden eagle | 1,835,290 | 18,725 | 1,682,140 | 134,425 |
|  | Eastern buzzard | 38,382,222 | 34,751,436 | 2,177,793 | 1,452,993 |
| Eurasian sparrowhawk | Eastern buzzard | 42,577,883 | 37,628,360 | 3,174,952 | 1,774,571 |
| Northern goshawk | Eastern buzzard | 39,466,874 | 35,855,277 | 1,959,888 | 1,651,709 |
| Cinereous vulture | Eastern buzzard | 39,205,764 | 35,746,288 | 1,914,278 | 1,545,198 |
| White-tailed eagle | Eastern buzzard | 30,118,652 | 27,187,103 | 1,689,202 | 1,242,347 |
| Oriental honey-buzzard | Eastern buzzard | 58,016,817 | 51,403,339 | 4,390,175 | 2,223,303 |
| Black kite | Eastern buzzard | 32,182,132 | 28,728,716 | 2,132,391 | 1,321,025 |
| Chinese sparrowhawk | Eastern buzzard | 42,894,254 | 37,580,432 | 3,493,265 | 1,820,557 |
| Turkey vulture | Eastern buzzard | 75,410,762 | 70,610,796 | 2,356,688 | 2,443,278 |
| Common kestrel | Common kestrel | 5,698,946 | 17,843 | 5,152,212 | 528,891 |
| Saker falcon | Saker falcon | 1,740,466 | 104,111 | 1,453,643 | 182,712 |
|  | Common kestrel | 18,704,134 | 16,177,271 | 1,579,414 | 947,449 |
| Peregrine falcon | Peregrine falcon | 1,598,481 | 84,009 | 1,335,088 | 179,384 |
|  | Common kestrel | 18,644,180 | 16,241,792 | 1,467,194 | 935,194 |
| Eurasian hobby | Common kestrel | 20,292,235 | 16,229,518 | 2,927,043 | 1,135,674 |
| American crow | Zebra finch | 73,241,772 | 70,138,649 | 1,579,705 | 1,523,418 |
| Medium ground-finch | Zebra finch | 64,983,101 | 60,762,787 | 2,320,313 | 1,900,001 |
| White-throated sparrow | Zebra finch | 63,988,552 | 58,620,584 | 3,678,356 | 1,689,612 |
| Common canary | Zebra finch | 63,451,159 | 57,750,404 | 4,152,188 | 1,548,567 |
| Collared flycatcher | Zebra finch | 62,837,663 | 58,568,302 | 2,861,400 | 1,407,961 |

**Table S14. Blood transcriptome *de novo* assembly and mapping statistics.** RPKM is reads per kilobase per million mapped reads.

**a.** Transcriptome *de novo* assembly results

| Species | RNA reads | Assembled transcripts | Non-bacterial transcripts | Candidate unigenes | Non-redundant unigenes | Expressed unigenes (RPKM ≥ 1.0) |
| --- | --- | --- | --- | --- | --- | --- |
| Eurasian eagle-owl | 74,807,936 | 192,557 | 191,773 | 94,871 | 43,765 | 30,948 |
| Oriental scops-owl | 80,047,874 | 188,436 | 186,833 | 77,140 | 35,146 | 26,197 |
| Himalayan owl | 80,236,848 | 215,142 | 213,061 | 105,270 | 43,741 | 32,589 |
| Northern boobook | 73,698,344 | 226,633 | 225,926 | 113,910 | 52,541 | 40,005 |
| Long-eared owl | 72,575,180 | 178,865 | 176,769 | 92,214 | 36,725 | 28,065 |
| Eastern buzzard | 65,713,128 | 177,945 | 177,425 | 82,313 | 33,088 | 26,847 |
| Eurasian sparrowhawk | 64,419,598 | 166,403 | 166,145 | 75,790 | 30,635 | 25,904 |
| Northern goshawk | 105,925,434 | 268,641 | 267,370 | 136,735 | 53,835 | 39,255 |
| Cinereous vulture | 60,236,628 | 173,662 | 172,758 | 89,347 | 35,912 | 29,182 |
| Common kestrel | 64,250,348 | 181,982 | 181,774 | 81,852 | 34,577 | 27,956 |
| Saker falcon | 76,863,908 | 124,489 | 123,135 | 40,333 | 22,158 | 18,509 |
| Peregrine falcon | 79,571,270 | 117,666 | 106,738 | 36,892 | 21,002 | 17,750 |
| Eurasian hobby | 500,017,118 | 394,468 | 394,145 | 289,855 | 200,718 | 162,083 |
| Grey-headed woodpecker | 79,112,614 | 164,135 | 163,403 | 89,073 | 40,888 | 35,233 |
| Eurasian siskin | 41,742,212 | 72,078 | 72,030 | 17,846 | 12,775 | 12,619 |
| Little egret | 77,748,608 | 197,944 | 197,464 | 91,981 | 41,795 | 31,053 |
| Striated heron | 73,160,250 | 167,558 | 166,990 | 81,217 | 37,962 | 27,845 |
| Black-faced spoonbill | 82,694,972 | 179,098 | 178,488 | 94,107 | 40,848 | 29,368 |

**b.** Transcriptome sequences mapping results to the genome assembly

| Species | RNA reads | Mapped reads | Mapping rate (%) | Concordant pair  alignment rate (%) |
| --- | --- | --- | --- | --- |
| Eurasian eagle-owl | 74,807,936 | 58,030,942 | 77.6 | 68.7 |
| Oriental scops-owl | 80,047,874 | 67,101,568 | 83.8 | 74.9 |
| Eastern buzzard | 65,713,128 | 58,221,624 | 88.6 | 81.5 |
| Common kestrel | 64,250,348 | 55,612,832 | 86.6 | 80.2 |
| Saker falcon | 76,863,908 | 53,324,296 | 69.4 | 58.1 |
| Peregrine falcon | 79,571,270 | 53,987,662 | 67.8 | 59.5 |
| Little egret | 77,748,608 | 64,132,289 | 82.5 | 74.0 |

**Table S15. Genome assembly information for the 25 avian species used in this study.** Eurasian eagle-owl, oriental scops-owl, eastern buzzard, and common kestrel genome assemblies were generated from the present study, while the other avian genome assemblies were downloaded from the NCBI database.

| Species | Scaffold N50 | Assembly | Maximum  longevity  (yrs) | Specimen origin | Avg.  body mass  (g) | Nocturnality | Major diet | Ref.  source |
| --- | --- | --- | --- | --- | --- | --- | --- | --- |
| Eurasian  eagle-owl | 29.92 Mb | - | 27.8 | Wild | 2708 | Nocturnal | Carnivorous | 12,13 |
| Northern spotted owl | 3.98 Mb | Soccid_v01 | 25 | Wild | 588.7 | Nocturnal | Carnivorous | 13, 14 |
| Oriental  scops-owl | 18.54 Mb | - | N/A  (6.8 yrs for Eurasian scops-owl) | N/A  (wild) | 85 | Nocturnal | Carnivorous | 13,14 |
| Barn Owl | 52.8 Kb | ASM68720v1 | 34 | Wild | 580 | Nocturnal | Carnivorous | 13,14 |
| Downy woodpecker | 2.09 Mb | ASM69900v1 | 11.9 | Wild | 27.0 | - | Omnivorous | 13,14 |
| Golden eagle | 9.23 Mb | Aquila_  chrysaetos-1.0.2 | 48 | Captivity | 4421 | - | Carnivorous | 13,14 |
| Eastern buzzard | 7.47 Mb | - | N/A  (28.8 yrs for common buzzard) | N/A  (wild) | 731 | - | Carnivorous | 13,14 |
| Bald eagle | 9.15 Mb | Haliaeetus_  leucocephalus-4.0 | 48 | Captivity | 4650 | - | Carnivorous | 13,14 |
| Saker falcon | 4.15 Mb | F_cherrug_v1.0 | 15.9 | Wild | 997.5 | - | Carnivorous | 13,14 |
| Peregrine falcon | 3.94 Mb | F_peregrinus_  v1.0 | 25 | Unknown | 920 | - | Carnivorous | 13,14 |
| Common kestrel | 21.23 Mb | - | 23.8 | Wild | 214 | - | Carnivorous | 13,14 |
| Zebra Finch | 8.24 Mb | Taeniopygia_  guttata-3.2.4 | 12 | Unknown | 10 | - | Herbivorous | 13,14 |
| American crow | 6.95 Mb | ASM69197v1 | 20 | Unknown | 412.5 | - | Omnivorous | 13,14 |
| Budgerigar | 10.61 Mb | Melopsittacus_  undulatus_6.3 | 21 | Captivity | 27.5 | - | Herbivorous | 13,14 |
| Little egret | 3.07 Mb | ASM68718v1 | 22.3 | Wild | 495 | - | Carnivorous | 13,14,15 |
| Crested Ibis | 5.21 Mb | ASM70822v1 | 25.8 | Captivity | 1900 | - | Carnivorous | 13,14 |
| Hoatzin | 2.94 Mb | ASM69207v1 | 14 | Captivity | 797.5 | - | Herbivorous | 13,16 |
| Killdeer | 3.66 Mb | ASM70802v2 | 10.9 | Wild | 96.5 | - | Omnivorous | 13,14,17 |
| Chuck-will’s-widow | 46.3 Kb | ASM70074v1 | 14.8 | Wild | 116.3 | Nocturnal | Carnivorous | 13,14 |
| Anna’s hummingbird | 4.05 Mb | ASM69908v1 | 8.5 | Wild | 4.3 | - | Omnivorous | 13,14,17 |
| Common cuckoo | 2.99 Mb | ASM70932v1 | 12.9 | Wild | 115 | - | Carnivorous | 13,14 |
| Rock dove | 3.15 Mb | Cliv_1.0 | 6.0 | Wild | 270 | - | Herbivorous | 13,17 |
| Chicken | 6.38 Mb | Gallus_gallus-5.0 | 30 | Captivity | 914 | - | Omnivorous | 13,14 |
| Brown kiwi | 3.96 Mb | AptMant0 | 35 | Captivity | 2527.5 | Nocturnal | Omnivorous | 13,14 |
| Common ostrich | 3.59 Mb | ASM69896v1 | 50 | Captivity | 111000.0 | - | Herbivorous | 13, 14 |

**Table S16. Commonly enriched Gene Ontology (GO) categories of expanded gene families in the ancestral branches of Strigiformes, Accipitriformes, and Falconiformes.** GO categories also enriched in expanded gene families of brown kiwi were excluded. *P*-value was calculated by Fisher’s exact test with a 5% FDR criterion.

| GO term | Description | *P*-value in the ancestral branch of Strigiformes | *P*-value in the ancestral branch of Accipitriformes | *P*-value in the ancestral branch of Falconiformes |
| --- | --- | --- | --- | --- |
| GO:0051494 | negative regulation of cytoskeleton organization | 2.81E-02 | 1.46E-03 | 5.53E-03 |
| GO:0001725 | stress fiber | 4.31E-04 | 6.42E-04 | 1.77E-02 |
| GO:0051272 | positive regulation of cellular component movement | 3.27E-02 | 3.87E-04 | 1.23E-03 |
| GO:0051271 | negative regulation of cellular component movement | 2.74E-02 | 4.27E-05 | 6.59E-03 |
| GO:0051270 | regulation of cellular component movement | 5.35E-03 | 1.44E-05 | 1.79E-03 |
| GO:0040017 | positive regulation of locomotion | 4.08E-02 | 5.64E-04 | 1.61E-03 |
| GO:0007165 | signal transduction | 4.81E-03 | 6.91E-04 | 6.42E-03 |
| GO:0007166 | cell surface receptor signaling pathway | 1.72E-02 | 4.01E-03 | 3.26E-03 |
| GO:0005829 | cytosol | 3.98E-05 | 2.69E-06 | 1.22E-03 |
| GO:0051241 | negative regulation of multicellular organismal process | 1.55E-02 | 1.83E-08 | 2.04E-03 |
| GO:0070887 | cellular response to chemical stimulus | 2.81E-03 | 1.66E-05 | 7.00E-04 |
| GO:0005102 | receptor binding | 1.93E-02 | 9.49E-03 | 2.41E-03 |
| GO:0032432 | actin filament bundle | 5.93E-04 | 9.68E-04 | 2.06E-02 |
| GO:0070507 | regulation of microtubule cytoskeleton organization | 1.97E-02 | 2.51E-02 | 1.48E-02 |
| GO:0005856 | cytoskeleton | 1.41E-09 | 5.52E-08 | 6.87E-03 |
| GO:0050818 | regulation of coagulation | 4.97E-02 | 2.24E-02 | 2.32E-03 |
| GO:0006355 | regulation of transcription, DNA-templated | 1.34E-02 | 2.01E-02 | 4.82E-02 |
| GO:0060041 | retina development in camera-type eye | 3.49E-03 | 9.05E-03 | 1.76E-02 |
| GO:1902904 | negative regulation of supramolecular fiber organization | 2.40E-02 | 1.06E-03 | 4.39E-02 |
| GO:1902903 | regulation of supramolecular fiber organization | 6.37E-03 | 1.61E-02 | 1.20E-02 |
| GO:0071356 | cellular response to tumor necrosis factor | 3.06E-02 | 1.49E-02 | 2.29E-02 |
| GO:0051336 | regulation of hydrolase activity | 3.45E-02 | 2.81E-03 | 5.29E-03 |
| GO:0051049 | regulation of transport | 3.22E-02 | 9.88E-03 | 1.87E-02 |
| GO:0010633 | negative regulation of epithelial cell migration | 2.81E-02 | 1.03E-03 | 1.83E-02 |
| GO:0050878 | regulation of body fluid levels | 4.32E-03 | 7.08E-04 | 4.69E-03 |
| GO:0031589 | cell-substrate adhesion | 7.55E-04 | 3.60E-03 | 1.05E-03 |
| GO:0050909 | sensory perception of taste | 2.26E-02 | 1.54E-02 | 3.01E-03 |
| GO:0097517 | contractile actin filament bundle | 4.31E-04 | 6.42E-04 | 1.77E-02 |
| GO:0048522 | positive regulation of cellular process | 1.13E-02 | 4.74E-02 | 1.60E-02 |
| GO:0051056 | regulation of small GTPase mediated signal transduction | 1.37E-02 | 2.60E-04 | 4.48E-02 |
| GO:0033043 | regulation of organelle organization | 2.06E-05 | 5.47E-03 | 7.69E-03 |
| GO:0032886 | regulation of microtubule-based process | 9.90E-03 | 9.24E-03 | 2.14E-02 |
| GO:0010591 | regulation of lamellipodium assembly | 2.91E-02 | 2.19E-02 | 1.28E-04 |
| GO:0040013 | negative regulation of locomotion | 2.39E-03 | 5.10E-05 | 7.28E-03 |
| GO:0040012 | regulation of locomotion | 2.11E-03 | 1.96E-04 | 1.95E-03 |
| GO:0030029 | actin filament-based process | 4.13E-05 | 4.43E-04 | 4.04E-03 |
| GO:0061035 | regulation of cartilage development | 7.34E-03 | 3.25E-03 | 2.70E-02 |
| GO:0009966 | regulation of signal transduction | 2.29E-02 | 7.31E-04 | 5.23E-03 |
| GO:0048518 | positive regulation of biological process | 5.98E-03 | 3.01E-03 | 5.35E-03 |
| GO:0010243 | response to organonitrogen compound | 1.04E-02 | 1.56E-05 | 1.02E-03 |
| GO:0090066 | regulation of anatomical structure size | 9.29E-05 | 3.18E-02 | 3.97E-03 |
| GO:0008092 | cytoskeletal protein binding | 7.48E-05 | 1.41E-05 | 2.11E-02 |
| GO:0045666 | positive regulation of neuron differentiation | 4.34E-02 | 3.64E-02 | 8.47E-03 |
| GO:0032879 | regulation of localization | 1.71E-02 | 2.27E-04 | 2.24E-03 |
| GO:0015629 | actin cytoskeleton | 2.46E-07 | 6.74E-05 | 2.81E-02 |
| GO:0000226 | microtubule cytoskeleton organization | 1.96E-04 | 3.79E-04 | 1.89E-02 |
| GO:0065009 | regulation of molecular function | 3.82E-02 | 4.33E-03 | 4.58E-02 |
| GO:0045596 | negative regulation of cell differentiation | 1.12E-02 | 9.58E-08 | 1.01E-02 |
| GO:0051093 | negative regulation of developmental process | 3.79E-03 | 9.25E-08 | 2.96E-03 |
| GO:0051092 | positive regulation of NF-kappaB transcription factor activity | 1.28E-02 | 3.05E-02 | 3.43E-03 |
| GO:0005884 | actin filament | 3.27E-05 | 1.04E-03 | 1.63E-02 |
| GO:0007015 | actin filament organization | 1.31E-03 | 1.53E-02 | 2.76E-02 |
| GO:0007017 | microtubule-based process | 1.93E-04 | 1.98E-02 | 3.59E-02 |
| GO:0044087 | regulation of cellular component biogenesis | 2.24E-04 | 2.67E-06 | 1.33E-04 |
| GO:0044089 | positive regulation of cellular component biogenesis | 1.04E-02 | 4.72E-03 | 1.38E-02 |
| GO:0043207 | response to external biotic stimulus | 4.87E-02 | 4.07E-02 | 1.17E-02 |
| GO:0120032 | regulation of plasma membrane bounded cell projection assembly | 2.76E-03 | 3.60E-03 | 1.25E-03 |
| GO:0003779 | actin binding | 8.12E-04 | 1.72E-03 | 7.93E-03 |
| GO:0050794 | regulation of cellular process | 1.46E-03 | 2.32E-02 | 4.16E-02 |
| GO:0097305 | response to alcohol | 6.62E-03 | 1.72E-02 | 2.78E-02 |
| GO:0042641 | actomyosin | 1.12E-04 | 8.32E-05 | 2.13E-02 |
| GO:2000181 | negative regulation of blood vessel morphogenesis | 4.65E-02 | 2.27E-03 | 2.29E-02 |
| GO:1901700 | response to oxygen-containing compound | 9.98E-03 | 1.15E-06 | 3.47E-05 |
| GO:1901701 | cellular response to oxygen-containing compound | 9.88E-03 | 3.84E-06 | 1.84E-04 |
| GO:0034110 | regulation of homotypic cell-cell adhesion | 2.91E-02 | 9.99E-03 | 2.13E-03 |
| GO:0044093 | positive regulation of molecular function | 4.14E-02 | 2.18E-03 | 7.03E-04 |
| GO:0060491 | regulation of cell projection assembly | 2.99E-03 | 3.90E-03 | 1.33E-03 |
| GO:0044430 | cytoskeletal part | 8.28E-06 | 1.17E-07 | 9.71E-03 |
| GO:0120025 | plasma membrane bounded cell projection | 3.45E-04 | 9.54E-05 | 1.19E-02 |
| GO:0050890 | cognition | 1.55E-04 | 2.23E-03 | 1.56E-04 |
| GO:0050769 | positive regulation of neurogenesis | 5.74E-03 | 2.47E-03 | 3.97E-03 |
| GO:1903034 | regulation of response to wounding | 4.31E-02 | 1.87E-02 | 1.08E-03 |
| GO:0005515 | protein binding | 6.72E-04 | 2.58E-03 | 3.96E-03 |
| GO:0032535 | regulation of cellular component size | 4.88E-04 | 2.41E-02 | 5.95E-03 |
| GO:0043226 | organelle | 3.13E-02 | 5.49E-03 | 2.50E-02 |
| GO:1900046 | regulation of hemostasis | 4.18E-02 | 1.88E-02 | 1.91E-03 |
| GO:0034097 | response to cytokine | 6.59E-04 | 3.08E-04 | 2.99E-02 |
| GO:0050954 | sensory perception of mechanical stimulus | 1.03E-04 | 1.07E-07 | 1.08E-03 |
| GO:0097435 | supramolecular fiber organization | 2.21E-04 | 1.58E-04 | 5.11E-03 |
| GO:0043005 | neuron projection | 2.49E-02 | 3.30E-04 | 3.24E-02 |
| GO:0006508 | proteolysis | 1.23E-03 | 5.92E-03 | 3.69E-03 |
| GO:0090330 | regulation of platelet aggregation | 1.34E-02 | 3.89E-03 | 8.12E-04 |
| GO:0022839 | ion gated channel activity | 6.00E-03 | 2.60E-02 | 4.57E-03 |
| GO:0010543 | regulation of platelet activation | 4.39E-02 | 1.69E-02 | 3.67E-03 |
| GO:1901698 | response to nitrogen compound | 1.51E-02 | 3.08E-05 | 1.47E-03 |
| GO:1901699 | cellular response to nitrogen compound | 8.04E-04 | 7.74E-05 | 2.60E-03 |
| GO:1902743 | regulation of lamellipodium organization | 6.86E-03 | 3.96E-02 | 3.17E-04 |
| GO:0099513 | polymeric cytoskeletal fiber | 4.15E-06 | 5.47E-06 | 9.83E-03 |
| GO:0009607 | response to biotic stimulus | 2.91E-02 | 2.21E-02 | 1.55E-02 |
| GO:0032956 | regulation of actin cytoskeleton organization | 4.97E-04 | 1.06E-03 | 1.95E-03 |
| GO:0038083 | peptidyl-tyrosine autophosphorylation | 3.27E-03 | 1.69E-02 | 3.01E-03 |
| GO:0016525 | negative regulation of angiogenesis | 4.33E-02 | 1.86E-03 | 2.13E-02 |
| GO:0060341 | regulation of cellular localization | 2.01E-02 | 2.79E-02 | 1.25E-02 |
| GO:0019904 | protein domain specific binding | 9.05E-03 | 2.44E-02 | 1.11E-02 |
| GO:0007611 | learning or memory | 1.40E-03 | 1.10E-03 | 7.64E-05 |
| GO:0042995 | cell projection | 8.63E-05 | 1.33E-05 | 4.69E-03 |
| GO:0010033 | response to organic substance | 6.80E-04 | 3.84E-07 | 1.11E-03 |
| GO:2000145 | regulation of cell motility | 5.92E-03 | 7.85E-05 | 1.10E-03 |
| GO:2000147 | positive regulation of cell motility | 2.71E-02 | 3.00E-04 | 1.06E-03 |
| GO:2000146 | negative regulation of cell motility | 2.07E-02 | 2.95E-05 | 5.47E-03 |
| GO:0032101 | regulation of response to external stimulus | 6.21E-03 | 1.72E-02 | 7.51E-03 |
| GO:0051240 | positive regulation of multicellular organismal process | 4.42E-04 | 2.53E-06 | 1.23E-04 |
| GO:0030326 | embryonic limb morphogenesis | 6.35E-04 | 9.87E-05 | 4.81E-02 |
| GO:0035113 | embryonic appendage morphogenesis | 6.71E-04 | 1.06E-04 | 4.92E-02 |
| GO:0042221 | response to chemical | 1.13E-03 | 4.13E-06 | 2.59E-04 |
| GO:0005604 | basement membrane | 2.42E-04 | 1.70E-04 | 2.21E-02 |
| GO:0048583 | regulation of response to stimulus | 1.20E-02 | 1.69E-03 | 7.41E-03 |
| GO:0007605 | sensory perception of sound | 7.38E-05 | 7.59E-08 | 9.38E-04 |
| GO:0022607 | cellular component assembly | 2.17E-02 | 2.60E-03 | 3.11E-02 |
| GO:0006952 | defense response | 2.40E-02 | 3.61E-02 | 2.21E-04 |
| GO:0010799 | regulation of peptidyl-threonine phosphorylation | 3.14E-02 | 1.68E-03 | 1.28E-04 |
| GO:0009719 | response to endogenous stimulus | 9.31E-04 | 1.67E-06 | 2.96E-03 |
| GO:0071229 | cellular response to acid chemical | 4.42E-02 | 2.05E-03 | 7.58E-03 |
| GO:0071310 | cellular response to organic substance | 2.82E-04 | 3.03E-06 | 2.50E-03 |
| GO:0030336 | negative regulation of cell migration | 1.63E-02 | 1.89E-05 | 4.38E-03 |
| GO:0030335 | positive regulation of cell migration | 2.38E-02 | 2.44E-04 | 9.10E-04 |
| GO:0030334 | regulation of cell migration | 4.06E-03 | 4.82E-05 | 7.93E-04 |
| GO:0034612 | response to tumor necrosis factor | 9.81E-03 | 5.07E-04 | 3.60E-02 |
| GO:0030193 | regulation of blood coagulation | 4.18E-02 | 1.88E-02 | 1.91E-03 |
| GO:0005737 | cytoplasm | 6.06E-04 | 1.19E-06 | 9.87E-03 |
| GO:0008017 | microtubule binding | 1.52E-02 | 1.09E-02 | 4.70E-02 |
| GO:0014910 | regulation of smooth muscle cell migration | 2.48E-04 | 2.54E-02 | 5.61E-03 |
| GO:0014912 | negative regulation of smooth muscle cell migration | 1.03E-02 | 7.67E-03 | 1.18E-03 |

**Table S17. Commonly enriched KEGG pathways of expanded gene families in the ancestral branches of Strigiformes, Accipitriformes, and Falconiformes.** KEGG pathways also enriched in expanded gene families of brown kiwi were excluded. *P*-value was calculated by Fisher’s exact test with a 5% FDR criterion.

| KEGG | Description | *P*-value in the ancestral branch of Strigiformes | *P*-value in the ancestral branch of Accipitriformes | *P*-value in the ancestral branch of Falconiformes |
| --- | --- | --- | --- | --- |
| ko05410 | Hypertrophic cardiomyopathy (HCM) | 9.52E-03 | 4.82E-04 | 4.04E-02 |
| ko05414 | Dilated cardiomyopathy (DCM) | 7.97E-08 | 1.05E-03 | 4.98E-02 |
| ko04750 | Inflammatory mediator regulation of TRP channels | 7.09E-09 | 7.71E-03 | 5.57E-04 |
| ko04261 | Adrenergic signaling in cardiomyocytes | 9.14E-05 | 9.21E-03 | 1.41E-02 |
| ko04015 | Rap1 signaling pathway | 1.79E-03 | 1.86E-04 | 3.13E-06 |

**Table S18. Gene Ontology (GO) enrichment of gene families that were expanded in the present birds of prey.** *P*-value was calculated by Fisher’s exact test with a 5% FDR criterion.

| GO term | Description | Fold-change of Avg. gene number | *P*-value |
| --- | --- | --- | --- |
| GO:0044403 | symbiosis, encompassing mutualism through parasitism | 2.03 | 5.89E-07 |
| GO:0044419 | interspecies interaction between organisms | 2.02 | 8.08E-06 |
| GO:0002223 | stimulatory C-type lectin receptor signaling pathway | 3.15 | 1.23E-04 |
| GO:0002220 | innate immune response activating cell surface receptor signaling pathway | 3.37 | 2.39E-04 |
| GO:0034377 | plasma lipoprotein particle assembly | 2.31 | 9.75E-04 |
| GO:1900026 | positive regulation of substrate adhesion-dependent cell spreading | 2.07 | 1.08E-03 |
| GO:0048295 | positive regulation of isotype switching to IgE isotypes | 2.27 | 1.12E-03 |
| GO:1900024 | regulation of substrate adhesion-dependent cell spreading | 2.05 | 1.64E-03 |
| GO:0002478 | antigen processing and presentation of exogenous peptide antigen | 2.12 | 1.70E-03 |
| GO:0019068 | virion assembly | 2.67 | 2.10E-03 |
| GO:2001044 | regulation of integrin-mediated signaling pathway | 2.19 | 2.10E-03 |
| GO:0048002 | antigen processing and presentation of peptide antigen | 2.14 | 2.28E-03 |
| GO:0051569 | regulation of histone H3-K4 methylation | 2.06 | 3.32E-03 |
| GO:0019884 | antigen processing and presentation of exogenous antigen | 2.17 | 3.49E-03 |
| GO:2001046 | positive regulation of integrin-mediated signaling pathway | 2.62 | 3.96E-03 |
| GO:0065005 | protein-lipid complex assembly | 2.91 | 4.02E-03 |
| GO:0070093 | negative regulation of glucagon secretion | 2.51 | 4.12E-03 |
| GO:1901381 | positive regulation of potassium ion transmembrane transport | 2.19 | 4.20E-03 |
| GO:0019886 | antigen processing and presentation of exogenous peptide antigen via MHC class II | 2.15 | 4.41E-03 |
| GO:0022832 | voltage-gated channel activity | 2.01 | 4.92E-03 |
| GO:0030206 | chondroitin sulfate biosynthetic process | 4.38 | 5.08E-03 |
| GO:1902495 | transmembrane transporter complex | 2.05 | 5.27E-03 |
| GO:0008331 | high voltage-gated calcium channel activity | 2.05 | 5.27E-03 |
| GO:0002495 | antigen processing and presentation of peptide antigen via MHC class II | 2.13 | 5.39E-03 |
| GO:0071827 | plasma lipoprotein particle organization | 3.09 | 5.39E-03 |
| GO:1902170 | cellular response to reactive nitrogen species | 2.14 | 5.74E-03 |
| GO:1902855 | regulation of non-motile cilium assembly | 2.11 | 5.74E-03 |
| GO:0034702 | ion channel complex | 2.08 | 6.18E-03 |
| GO:0002504 | antigen processing and presentation of peptide or polysaccharide antigen via MHC class II | 2.1 | 6.53E-03 |
| GO:1905476 | negative regulation of protein localization to membrane | 2.23 | 7.76E-03 |
| GO:0038095 | Fc-epsilon receptor signaling pathway | 2.93 | 7.84E-03 |
| GO:0043312 | neutrophil degranulation | 2.11 | 8.81E-03 |
| GO:0003846 | 2-acylglycerol O-acyltransferase activity | 10.89 | 9.10E-03 |
| GO:0042590 | antigen processing and presentation of exogenous peptide antigen via MHC class I | 4.24 | 9.50E-03 |
| GO:0048293 | regulation of isotype switching to IgE isotypes | 2.51 | 9.50E-03 |
| GO:0031523 | Myb complex | 2.88 | 1.00E-02 |
| GO:0051120 | hepoxilin A3 synthase activity | 10.89 | 1.05E-02 |
| GO:0050252 | retinol O-fatty-acyltransferase activity | 12.44 | 1.05E-02 |
| GO:1990189 | peptide-serine-N-acetyltransferase activity | 3.11 | 1.05E-02 |
| GO:0047323 | [3-methyl-2-oxobutanoate dehydrogenase (acetyl-transferring)] kinase activity | 5.29 | 1.05E-02 |
| GO:1900121 | negative regulation of receptor binding | 2.21 | 1.08E-02 |
| GO:0002296 | T-helper 1 cell lineage commitment | 2.77 | 1.08E-02 |
| GO:0036155 | acylglycerol acyl-chain remodeling | 14 | 1.08E-02 |
| GO:0034178 | toll-like receptor 13 signaling pathway | 2.59 | 1.08E-02 |
| GO:1901731 | positive regulation of platelet aggregation | 2.31 | 1.08E-02 |
| GO:1902044 | regulation of Fas signaling pathway | 2.18 | 1.08E-02 |
| GO:0001844 | protein insertion into mitochondrial membrane involved in apoptotic signaling pathway | 2.33 | 1.08E-02 |
| GO:0034465 | response to carbon monoxide | 2.09 | 1.08E-02 |
| GO:0036149 | phosphatidylinositol acyl-chain remodeling | 4.04 | 1.08E-02 |
| GO:1902045 | negative regulation of Fas signaling pathway | 2.8 | 1.08E-02 |
| GO:0008615 | pyridoxine biosynthetic process | 2.07 | 1.08E-02 |
| GO:0008614 | pyridoxine metabolic process | 2.42 | 1.08E-02 |
| GO:0001796 | regulation of type IIa hypersensitivity | 2.16 | 1.08E-02 |
| GO:1905453 | regulation of myeloid progenitor cell differentiation | 7 | 1.08E-02 |
| GO:0035579 | specific granule membrane | 4.77 | 1.10E-02 |
| GO:0071825 | protein-lipid complex subunit organization | 3.4 | 1.11E-02 |
| GO:0000789 | cytoplasmic chromatin | 2.59 | 1.11E-02 |
| GO:0002283 | neutrophil activation involved in immune response | 2.61 | 1.16E-02 |
| GO:0043268 | positive regulation of potassium ion transport | 2.16 | 1.16E-02 |
| GO:0033137 | negative regulation of peptidyl-serine phosphorylation | 2.08 | 1.18E-02 |
| GO:0005249 | voltage-gated potassium channel activity | 2.11 | 1.33E-02 |

**Table S19. List of genes showing accelerated *d_N_*/*d_S_* in the three raptor orders**

| Genes | ω Three raptor branches | | | | ω outgroups |
| --- | --- | --- | --- | --- | --- |
|  | Strigiformes | Accipitriformes | Falconiformes | Merged |  |
| *LOC422316* | 0.7250 | 0.2188 | 0.3676 | 0.4146 | 0.1253 |
| *RNH1* | 0.5599 | 0.3043 | 0.5248 | 0.4822 | 0.1340 |
| *RIF1* | 0.4231 | 0.5985 | 0.6027 | 0.5556 | 0.3007 |
| *VLDLR* | 0.4052 | 0.5618 | 0.4375 | 0.4731 | 0.1466 |
| *IL4I1* | 1.3922 | 0.2168 | 0.3847 | 0.4886 | 0.1801 |
| *SKP2* | 1.3782 | 0.6191 | 0.7180 | 0.9569 | 0.2997 |
| *SMC4* | 0.4212 | 0.4064 | 0.4548 | 0.4350 | 0.2166 |
| *MUM1* | 0.9040 | 0.5793 | 0.6939 | 0.7164 | 0.3299 |
| *ORC1* | 0.5132 | 0.4786 | 1.0003 | 0.7373 | 0.3508 |
| *DNAJC25* | 1.1660 | 0.1189 | 0.4075 | 0.3706 | 0.0647 |
| *COL3A1* | 0.5096 | 0.2897 | 0.2788 | 0.3436 | 0.1746 |
| *GMPS* | 0.1401 | 0.0521 | 0.1685 | 0.1259 | 0.0166 |
| *ITFG1* | 0.4411 | 0.4819 | 0.5602 | 0.5029 | 0.1608 |
| *LMNA* | 1.0153 | 0.9055 | 0.8907 | 0.9214 | 0.4318 |
| *DNAJC21* | 0.5231 | 0.8603 | 0.4373 | 0.6237 | 0.2616 |
| *HELB* | 0.7658 | 1.1112 | 0.4070 | 0.7014 | 0.3765 |
| *C5* | 0.4088 | 0.4190 | 0.4032 | 0.4098 | 0.2447 |
| *LOC107048990* | 0.7351 | 0.2973 | 0.3189 | 0.4411 | 0.2321 |
| *MRPL37* | 0.7049 | 0.9437 | 0.9591 | 0.8887 | 0.2786 |
| *NCAPG* | 0.6532 | 0.5192 | 0.6578 | 0.6243 | 0.3423 |
| *TSPO2* | 4.1893 | 0.4787 | 1.2710 | 1.2814 | 0.1887 |
| *RAD50* | 0.1508 | 0.2132 | 0.2709 | 0.2286 | 0.1179 |
| *SLC22A7* | 0.4227 | 0.4236 | 0.4662 | 0.4376 | 0.2099 |
| *SMC6* | 0.6001 | 0.3487 | 0.5704 | 0.5198 | 0.3019 |
| *ETV4* | 0.3241 | 0.7722 | 0.5582 | 0.5281 | 0.2409 |
| *LEMD2* | 0.5947 | 0.4558 | 0.4652 | 0.4546 | 0.2083 |
| *KCNQ1* | 0.2666 | 0.0690 | 0.2292 | 0.1870 | 0.0644 |
| *CENPC* | 0.9169 | 1.3188 | 1.3028 | 1.1991 | 0.7251 |
| *SPCS1* | 0.1741 | 3.2942 | 0.6508 | 1.3882 | 0.1456 |
| *ELAVL2* | 0.2034 | 0.0607 | 0.3010 | 0.1946 | 0.0587 |
| *KNTC1* | 0.3165 | 0.2435 | 0.4318 | 0.3490 | 0.2371 |
| *KIAA1551* | 1.1546 | 1.3174 | 0.7757 | 0.9800 | 0.6994 |
| *CENPL* | 0.7673 | 0.5784 | 0.4658 | 0.5660 | 0.2346 |
| *DIS3L* | 0.1537 | 0.4051 | 0.1708 | 0.2379 | 0.1297 |
| *NSMCE4A* | 1.1903 | 0.4201 | 0.8415 | 0.6702 | 0.2963 |
| *GLIS3* | 0.9297 | 0.6221 | 0.8195 | 0.7836 | 0.4803 |
| *RHCE* | 1.3586 | 1.6996 | 1.0154 | 1.2200 | 0.5970 |
| *SNAP47* | 0.3797 | 0.1633 | 0.3045 | 0.2815 | 0.1036 |
| *TCFL5* | 0.8234 | 0.4243 | 0.5573 | 0.6004 | 0.3415 |
| *GFM2* | 0.3309 | 0.2564 | 0.4258 | 0.3401 | 0.1830 |
| *AP1AR* | 0.4730 | 0.2743 | 0.3456 | 0.3500 | 0.0881 |
| *ORC2* | 0.8864 | 0.4046 | 0.6894 | 0.6752 | 0.3308 |
| *NANOG* | 0.7016 | 0.3990 | 0.9100 | 0.6286 | 0.2764 |
| *DNAJC8* | 0.4367 | 0.8052 | 0.0335 | 0.1867 | 0.0286 |
| *PRPH2* | 0.4387 | 0.1431 | 0.2196 | 0.2403 | 0.0870 |
| *LOC770429* | 0.5131 | 0.2316 | 0.3097 | 0.3657 | 0.1871 |
| *FOCAD* | 0.4374 | 0.3309 | 0.4922 | 0.4310 | 0.3004 |
| *LIG4* | 0.1824 | 0.2141 | 0.3160 | 0.2448 | 0.1378 |
| *POLRMT* | 0.2583 | 0.2572 | 0.3023 | 0.2782 | 0.1846 |
| *SS18L1* | 0.8364 | 0.1186 | 0.0581 | 0.1452 | 0.0255 |
| *KBTBD2* | 0.0436 | 0.0154 | 0.0575 | 0.0432 | 0.0099 |
| *KPNA7* | 0.2822 | 0.3663 | 0.6127 | 0.4376 | 0.2511 |
| *MMP1* | 0.4442 | 0.9045 | 0.4847 | 0.5787 | 0.2872 |
| *RAD21L1* | 0.6515 | 0.4188 | 0.7522 | 0.6250 | 0.3461 |
| *MIS12* | 0.4099 | 0.2780 | 0.3949 | 0.3795 | 0.1602 |
| *TBPL2* | 0.5624 | 0.3218 | 0.6313 | 0.4961 | 0.2344 |
| *LRWD1* | 0.2260 | 0.2716 | 0.4269 | 0.3063 | 0.1751 |
| *POLR2D* | 0.0427 | 0.3664 | 0.2344 | 0.1768 | 0.0251 |
| *KIAA1033* | 0.1552 | 0.1328 | 0.1589 | 0.1366 | 0.0712 |
| *UBE2G2* | 0.1511 | 0.0872 | 0.2136 | 0.1675 | 0.0233 |
| *TMEM116* | 0.3217 | 0.3364 | 0.3495 | 0.3190 | 0.1477 |
| *STX8* | 0.7933 | 0.1897 | 0.1626 | 0.3038 | 0.1201 |
| *CNTD1* | 1.2119 | 0.2612 | 0.5072 | 0.4879 | 0.2434 |
| *NUP93* | 0.2211 | 0.1128 | 0.1113 | 0.1364 | 0.0570 |
| *OVOL2* | 0.1540 | 0.0696 | 0.3300 | 0.1984 | 0.0502 |
| *DBF4* | 0.4026 | 0.6240 | 0.6836 | 0.5856 | 0.3440 |
| *TECPR1* | 0.2763 | 0.2596 | 0.1825 | 0.2300 | 0.1396 |
| *MMP10* | 0.4672 | 0.3015 | 0.4835 | 0.4404 | 0.2370 |
| *LOC100859557* | 0.5256 | 0.6479 | 0.4993 | 0.5387 | 0.3572 |
| *MEIOB* | 0.4184 | 0.2747 | 0.3330 | 0.3513 | 0.1727 |
| *ALKBH4* | 0.4927 | 0.4464 | 0.5723 | 0.5166 | 0.1860 |
| *ING3* | 0.1429 | 0.0374 | 0.2016 | 0.1333 | 0.0313 |
| *TOPORS* | 0.4527 | 0.4135 | 0.4088 | 0.4233 | 0.2882 |
| *TGM6L* | 0.3119 | 0.2491 | 0.3588 | 0.3217 | 0.1732 |
| *ZDHHC12* | 0.8205 | 0.6576 | 0.5786 | 0.6555 | 0.2401 |
| *HP1BP3* | 0.2248 | 0.1323 | 0.2448 | 0.2165 | 0.0939 |
| *SLC24A1* | 0.5810 | 0.4383 | 0.3935 | 0.4639 | 0.2598 |
| *SRP68* | 0.1117 | 0.1310 | 0.0533 | 0.0921 | 0.0299 |
| *TMEM175* | 0.4089 | 0.2286 | 0.3711 | 0.3423 | 0.1590 |
| *SPRED2* | 0.1132 | 0.1351 | 0.0933 | 0.1109 | 0.0270 |
| *TMEM9B* | 0.0789 | 1.9266 | 0.0842 | 0.2112 | 0.0232 |
| *WDR76* | 0.2154 | 0.3973 | 0.3036 | 0.3042 | 0.1728 |
| *RHAG* | 0.5409 | 0.9078 | 0.9742 | 0.8175 | 0.4571 |
| *CPLX4* | 0.7597 | 0.0882 | 0.1211 | 0.2543 | 0.0574 |
| *A1CF* | 0.1122 | 0.1163 | 0.0611 | 0.0928 | 0.0419 |
| *MYEF2* | 0.3063 | 0.3647 | 0.2974 | 0.3117 | 0.1458 |
| *TMEM136-2* | 0.2489 | 0.1882 | 0.2055 | 0.1874 | 0.0834 |
| *NDUFS3* | 0.3365 | 0.2764 | 0.2446 | 0.2422 | 0.0787 |
| *ANO5* | 0.2622 | 0.0901 | 0.0808 | 0.1343 | 0.0784 |
| *FUT11* | 0.2902 | 0.3001 | 0.1422 | 0.2026 | 0.1027 |
| *L2HGDH* | 0.2944 | 0.1769 | 0.2452 | 0.2524 | 0.1130 |
| *TRIM25* | 0.8545 | 0.9449 | 0.6694 | 0.7773 | 0.4986 |
| *GGT5* | 0.4894 | 0.6780 | 0.4471 | 0.5422 | 0.3287 |
| *FLI1* | 0.0711 | 0.2944 | 0.0851 | 0.1456 | 0.0404 |
| *MAK16* | 0.1454 | 0.3380 | 0.0665 | 0.1538 | 0.0638 |
| *UVSSA* | 0.4039 | 0.3080 | 0.4887 | 0.4105 | 0.2612 |
| *AAGAB* | 0.2813 | 0.2481 | 0.6144 | 0.3637 | 0.1667 |
| *SNAPC3* | 0.6118 | 0.1719 | 0.4173 | 0.4124 | 0.1666 |
| *SOD1* | 0.7430 | 1.1950 | 0.7399 | 0.8194 | 0.2546 |
| *MUC4* | 0.5334 | 0.5018 | 0.3805 | 0.4498 | 0.3563 |
| *TRMT10C* | 1.7110 | 0.3671 | 0.4010 | 0.5465 | 0.3053 |
| *SURF6* | 0.4193 | 0.3794 | 0.3994 | 0.4062 | 0.2039 |
| *CDC37L1* | 0.7698 | 0.4973 | 0.3091 | 0.4928 | 0.2516 |
| *RMDN3* | 0.3245 | 0.2037 | 0.2693 | 0.2695 | 0.1439 |
| *FUBP3* | 0.5092 | 0.2864 | 0.1051 | 0.1954 | 0.0688 |
| *AREG* | 0.4658 | 0.2970 | 0.6075 | 0.4470 | 0.1661 |
| *MED7* | 0.0504 | 0.1272 | 0.2549 | 0.1925 | 0.0479 |
| *TPX2* | 0.3119 | 0.5454 | 0.4831 | 0.4414 | 0.2940 |
| *N4BP2L2* | 1.0204 | 0.6232 | 1.1135 | 0.8609 | 0.5532 |
| *MRPS11* | 2.3194 | 0.5127 | 0.4985 | 0.6119 | 0.1897 |
| *ORC3* | 0.7933 | 0.4183 | 0.5177 | 0.5589 | 0.3409 |
| *APITD1* | 1.1615 | 0.3422 | 0.3609 | 0.5579 | 0.2415 |
| *CDC20* | 0.2017 | 0.2089 | 0.2715 | 0.2410 | 0.1270 |
| *CD14* | 0.4434 | 0.4728 | 0.4999 | 0.4836 | 0.2875 |
| *MMD* | 0.0797 | 0.3431 | 0.0983 | 0.1359 | 0.0378 |
| *RPAIN* | 0.5969 | 0.4616 | 0.3522 | 0.4030 | 0.1821 |
| *CHODL* | 0.2043 | 0.2240 | 0.4596 | 0.3084 | 0.1409 |
| *CEP63* | 0.5881 | 0.6250 | 0.5379 | 0.5697 | 0.3658 |
| *MTHFS* | 0.7633 | 1.1967 | 0.3747 | 0.8158 | 0.1910 |
| *DCAF5* | 0.1936 | 0.3209 | 0.1686 | 0.2083 | 0.1221 |
| *EIF2AK4* | 0.1558 | 0.2401 | 0.1963 | 0.1998 | 0.1384 |
| *MARS2* | 0.1934 | 0.2755 | 0.2914 | 0.2675 | 0.1497 |
| *USP30* | 0.2130 | 0.2193 | 0.1992 | 0.2153 | 0.1030 |
| *RD3* | 0.6943 | 0.2254 | 0.2965 | 0.3380 | 0.1105 |
| *MGP* | 0.4539 | 0.6842 | 1.3075 | 0.8335 | 0.3609 |
| *SLMO1* | 0.1772 | 0.5650 | 0.1384 | 0.2609 | 0.0923 |
| *C3H2orf71* | 0.6408 | 0.4491 | 0.4959 | 0.5177 | 0.3859 |
| *UTP18* | 0.3422 | 0.3808 | 0.2709 | 0.3220 | 0.1941 |
| *NUDT19* | 0.8512 | 0.4439 | 0.3953 | 0.5564 | 0.2914 |
| *OCSTAMP* | 0.5514 | 0.4786 | 0.6797 | 0.6036 | 0.3782 |
| *PLSCR5* | 0.3566 | 0.2182 | 0.4803 | 0.3904 | 0.1749 |
| *OPN4-1* | 0.3445 | 0.2404 | 0.4861 | 0.3658 | 0.2214 |
| *SLC30A5* | 0.1695 | 0.1483 | 0.0900 | 0.1201 | 0.0584 |
| *RALGAPB* | 0.0561 | 0.0569 | 0.0802 | 0.0673 | 0.0372 |
| *RLF* | 0.1986 | 0.1728 | 0.2291 | 0.2093 | 0.1540 |
| *ABCB7* | 0.1412 | 0.2699 | 0.1560 | 0.1740 | 0.0917 |
| *PPARA* | 0.0758 | 0.0634 | 0.1495 | 0.1101 | 0.0358 |
| *ACRC* | 0.4677 | 0.4080 | 0.7583 | 0.5544 | 0.3506 |
| *LOC428778* | 0.5008 | 0.4180 | 0.4442 | 0.4568 | 0.2842 |
| *LRRC57* | 0.1208 | 0.3721 | 0.1769 | 0.2066 | 0.0670 |
| *N4BP1* | 0.5323 | 0.3726 | 0.6144 | 0.5338 | 0.3709 |
| *MRPL45* | 0.2253 | 0.2366 | 0.4585 | 0.3219 | 0.1530 |
| *METTL6* | 0.3089 | 0.3451 | 0.5078 | 0.3894 | 0.1765 |
| *ORC6* | 0.9626 | 0.3750 | 0.6522 | 0.5851 | 0.3124 |
| *PARL* | 0.2813 | 0.4540 | 0.1711 | 0.2587 | 0.1026 |
| *RP5-1028K7.3* | 0.7597 | 1.6145 | 0.2441 | 0.4241 | 0.1928 |
| *SMG8* | 0.1183 | 0.0688 | 0.0875 | 0.0894 | 0.0511 |
| *CCNA2* | 0.2925 | 0.4920 | 0.3058 | 0.3211 | 0.1584 |

**Table S20. Selection constraints of beak development associated genes in the ancestral branches of birds of prey.** Genes showing accelerated *d_N_*/*d_S_* in the ancestral branches of the three raptor orders are highlighted in bold.

| Gene | ω Three raptor branches | | | | ω outgroups |
| --- | --- | --- | --- | --- | --- |
|  | Strigiformes | Accipitriformes | Falconiformes | Merged |  |
| *GDF9* | **0.3145** | **0.3211** | **0.1781** | **0.2365** | **0.1730** |
| *GJB5* | **0.2690** | **0.2442** | **5.5951** | **0.2540** | **0.1154** |
| *NAB1* | **0.1287** | **0.0698** | **0.0608** | **0.0653** | **0.0439** |
| *TRIP11* | **0.2669** | **0.2365** | **0.2437** | **0.2482** | **0.2115** |
| *BMP10* | 0.0524 | **0.1051** | **0.0981** | **0.0961** | 0.0587 |
| *BMP3* | 0.1085 | **0.3696** | **0.3627** | **0.3424** | 0.2367 |
| *GDF5* | 0.0728 | **0.1110** | **0.1349** | **0.1134** | 0.0780 |
| *GPRC5C* | **0.0869** | 0.0812 | **0.1314** | **0.1104** | 0.0817 |
| *HIC1* | **1.0294** | 0.0001 | **0.0264** | **0.0209** | 0.0154 |
| *WNT6* | **0.0367** | 0.0058 | **0.0174** | **0.0126** | 0.0093 |
| *TAF3* | **0.1182** | 0.0591 | **0.1375** | 0.1076 | 0.1180 |
| *BMP15* | 0.1590 | 0.1672 | **0.2764** | **0.2244** | 0.1714 |
| *MFNG* | 0.0001 | 0.1409 | **0.3921** | **0.2468** | 0.2100 |
| *RARB* | 0.0286 | 0.0266 | **0.1642** | **0.0840** | 0.0717 |
| *SATB2* | 0.0001 | 0.0001 | **0.0498** | **0.0363** | 0.0311 |
| *DLX5* | 0.0001 | 0.0001 | **0.2119** | 0.0974 | 0.1458 |
| *RREB1* | 0.1094 | 0.1496 | **0.2076** | 0.1626 | 0.1937 |
| *TGFB2* | 0.0001 | 0.0001 | **0.0220** | 0.0121 | 0.0201 |
| *GATA6* | **0.1612** | **0.1554** | 0.0278 | 0.0894 | 0.0994 |
| *KLF13* | **999** | **0.1993** | 0.0407 | 0.1231 | 0.1588 |
| *PITX2* | **0.0879** | **0.1091** | 0.0001 | 0.0001 | 0.0684 |
| *CDKN3* | 0.2747 | **0.5323** | 0.2549 | **0.3203** | 0.2768 |
| *NR3C2* | 0.0527 | **0.2807** | 0.0634 | **0.0877** | 0.0789 |
| *PTTG1IP* | 0.1018 | **0.3244** | 0.0654 | **0.1771** | 0.1138 |
| *THBS3* | 0.0155 | **0.1662** | 0.0231 | **0.0575** | 0.0401 |
| *FGF10* | 0.0001 | **0.1811** | 0.0001 | 0.0472 | 0.0591 |
| *PAX9* | 0.0001 | **0.0854** | 0.0001 | 0.0244 | 0.0816 |
| *SCML4* | 0.1783 | **0.2184** | 0.1186 | 0.1465 | 0.1898 |
| *CAMK2A* | **0.1514** | 0.0001 | 0.0001 | **0.0532** | 0.0136 |
| *FGF8* | **0.0720** | 0.0001 | 0.0001 | **0.0331** | 0.0263 |
| *FZD1* | **0.0745** | 0.0001 | 0.0001 | 0.0001 | 0.0118 |
| *NFE2L2* | **0.5749** | 0.1687 | 0.2810 | 0.2607 | 0.2974 |
| *PFKL* | **0.0305** | 0.0001 | 0.0036 | 0.0125 | 0.0146 |
| *TGFBI* | **0.2108** | 0.0566 | 0.0339 | 0.0609 | 0.0715 |
| *BMP1* | 0.0199 | 0.0264 | 0.0127 | 0.0194 | 0.0386 |
| *BMP4* | 0.0001 | 0.0001 | 0.0130 | 0.0083 | 0.0635 |
| *DLX1* | 0.0381 | 0.0001 | 0.0001 | 0.0001 | 0.0985 |
| *FGF13* | 0.0001 | 0.0001 | 0.0001 | 0.0001 | 0.0726 |
| *FGF2* | 0.0001 | 0.0001 | 0.0001 | 0.0001 | 0.1616 |
| *IRX2* | 0.0533 | 0.0848 | 0.0610 | 0.0651 | 0.0953 |
| *LZTR1* | 0.0188 | 0.0238 | 0.0001 | 0.0093 | 0.0251 |
| *MSX1* | 0.0001 | 0.0001 | 0.0001 | 0.0001 | 0.0454 |
| *MSX2* | 0.0001 | 0.0001 | 0.0001 | 0.0001 | 0.0718 |
| *MYCBPAP* | 0.4091 | 0.5645 | 0.4794 | 0.4740 | 0.7320 |
| *PAX6* | 0.0001 | 0.0001 | 0.0001 | 0.0001 | 0.0309 |
| *PRDM5* | 0.0001 | 0.0218 | 0.0311 | 0.0239 | 0.0753 |
| *PTCH2* | 0.0443 | 0.0302 | 0.0352 | 0.0360 | 0.0482 |
| *TCF20* | 0.0470 | 0.1163 | 0.1149 | 0.1139 | 0.1261 |
| *TLX1* | 0.0001 | 0.0001 | 0.0001 | 0.0001 | 0.0193 |
| *TRIM9* | 0.0001 | 0.0001 | 0.0084 | 0.0040 | 0.0164 |

**Table S21. Gene Ontology (GO) enrichment of genes that have a high level of GC3 biases in the bird of prey genomes.** *P*-value was calculated by Fisher’s exact test. Only GO categories with *P* < 5.00E-03 are shown.

| GO term | Description | *P*-value | FDR |
| --- | --- | --- | --- |
| GO:0006928 | movement of cell or subcellular component | 2.43E-04 | 9.12E-02 |
| GO:0070309 | lens fiber cell morphogenesis | 4.04E-04 | 9.12E-02 |
| GO:0033077 | T cell differentiation in thymus | 6.14E-04 | 9.12E-02 |
| GO:0033151 | V(D)J recombination | 6.14E-04 | 9.12E-02 |
| GO:0040011 | locomotion | 9.58E-04 | 9.12E-02 |
| GO:0002568 | somatic diversification of T cell receptor genes | 1.20E-03 | 9.12E-02 |
| GO:0033153 | T cell receptor V(D)J recombination | 1.20E-03 | 9.12E-02 |
| GO:0002681 | somatic recombination of T cell receptor gene segments | 1.20E-03 | 9.12E-02 |
| GO:0048538 | thymus development | 2.47E-03 | 9.12E-02 |
| GO:0051965 | positive regulation of synapse assembly | 2.47E-03 | 9.12E-02 |
| GO:0021953 | central nervous system neuron differentiation | 3.05E-03 | 9.12E-02 |
| GO:0030217 | T cell differentiation | 3.23E-03 | 9.12E-02 |
| GO:0002200 | somatic diversification of immune receptors | 3.69E-03 | 9.12E-02 |
| GO:0016444 | somatic cell DNA recombination | 3.69E-03 | 9.12E-02 |
| GO:0002562 | somatic diversification of immune receptors via germline recombination within a single locus | 3.69E-03 | 9.12E-02 |
| GO:0002089 | lens morphogenesis in camera-type eye | 3.88E-03 | 9.12E-02 |
| GO:0048870 | cell motility | 5.00E-03 | 9.12E-02 |
| GO:0051963 | regulation of synapse assembly | 5.23E-03 | 9.12E-02 |
| GO:0060561 | apoptotic process involved in morphogenesis | 5.75E-03 | 9.12E-02 |
| GO:1902742 | apoptotic process involved in development | 5.75E-03 | 9.12E-02 |
| GO:0005539 | glycosaminoglycan binding | 5.95E-03 | 9.12E-02 |
| GO:0097485 | neuron projection guidance | 6.18E-03 | 9.12E-02 |
| GO:0007411 | axon guidance | 6.18E-03 | 9.12E-02 |
| GO:0008283 | cell proliferation | 6.38E-03 | 9.12E-02 |
| GO:0051960 | regulation of nervous system development | 6.58E-03 | 9.12E-02 |
| GO:0072376 | protein activation cascade | 7.94E-03 | 9.12E-02 |
| GO:0000904 | cell morphogenesis involved in differentiation | 8.16E-03 | 9.12E-02 |
| GO:0016477 | cell migration | 8.76E-03 | 9.12E-02 |
| GO:0008219 | cell death | 9.18E-03 | 9.12E-02 |
| GO:0050807 | regulation of synapse organization | 9.31E-03 | 9.12E-02 |
| GO:0045165 | cell fate commitment | 9.31E-03 | 9.12E-02 |
| GO:0030154 | cell differentiation | 9.61E-03 | 9.12E-02 |

**Table S22. Statistics regarding highly conserved regions in Strigiformes, Accipitriformes, Falconiformes, and Passeriformes**

**a.** Identification of highly conserved regions (HCRs) in the four avian orders

| Orders | Reference  genome size | The number of windows (100Kb window,  >80% of sufficiently covered) | | Highly conserved windows (100Kb window,  Adjusted *P*-value < 0.0001) | | |
| --- | --- | --- | --- | --- | --- | --- |
|  |  | Window count | Non-overlapped  length (bp) | Window count | Non-overlapped  length (bp) | Percentage |
| Strigiformes | 1,258,075,470 | 117,593 | 1,211,056,698 | 31,477 | 509,623,240 | 42.08% |
| Accipitriformes | 1,259,752,395 | 117,508 | 1,215,179,433 | 35,908 | 579,312,028 | 47.67% |
| Falconiformes | 1,179,916,286 | 113,047 | 1,148,371,940 | 28,695 | 454,032,712 | 39.54% |
| Passeriformes | 1,232,135,591 | 103,837 | 1,046,118,050 | 27,338 | 420,336,735 | 40.18% |

**b.** Statistics of genes present in the HCRs

| Species | Total number of genes  in HCR regions  (with 50% coverage of gene) | # of specific  genes | # of exclusively shared genes  among the three bird of prey orders | # of shared genes  among the four orders |
| --- | --- | --- | --- | --- |
| Strigiformes | 3,423 | 733 | 765 | 420 |
| Accipitriformes | 3,897 | 1,122 |  |  |
| Falconiformes | 2,933 | 736 |  |  |
| Passeriformes | 4,694 | 1,478 | - |  |

**Table S23. Commonly enriched Gene Ontology (GO) categories of genes in the highly conserved genomic regions (HCRs) of Strigiformes, Accipitriformes, and Falconiformes.** GO categories also enriched in the HCRs of Passeriformes were excluded. *P*-value was calculated by Fisher’s exact test with a 5% FDR criterion.

| GO term | Description | *P*-value in Strigiformes | *P*-value in Accipitriformes | *P*-value in Falconiformes |
| --- | --- | --- | --- | --- |
| GO:0048869 | cellular developmental process | 8.72E-13 | 1.78E-10 | 1.75E-24 |
| GO:0051239 | regulation of multicellular organismal process | 2.87E-11 | 1.55E-04 | 2.10E-15 |
| GO:0050794 | regulation of cellular process | 5.87E-11 | 1.79E-08 | 1.22E-09 |
| GO:0050789 | regulation of biological process | 3.83E-10 | 2.53E-08 | 4.83E-10 |
| GO:0048646 | anatomical structure formation involved in morphogenesis | 7.78E-10 | 7.24E-06 | 9.14E-14 |
| GO:0008283 | cell proliferation | 9.24E-10 | 1.40E-05 | 3.21E-11 |
| GO:0065007 | biological regulation | 2.98E-09 | 2.02E-06 | 1.56E-09 |
| GO:0001763 | morphogenesis of a branching structure | 9.10E-09 | 2.34E-06 | 6.54E-12 |
| GO:0051241 | negative regulation of multicellular organismal process | 1.22E-08 | 1.68E-04 | 3.67E-10 |
| GO:0009888 | tissue development | 1.33E-08 | 7.38E-07 | 1.04E-14 |
| GO:0010941 | regulation of cell death | 1.80E-08 | 1.29E-03 | 2.82E-05 |
| GO:0061138 | morphogenesis of a branching epithelium | 4.63E-08 | 8.33E-06 | 1.04E-11 |
| GO:0030278 | regulation of ossification | 6.19E-08 | 1.45E-05 | 9.42E-08 |
| GO:0048754 | branching morphogenesis of an epithelial tube | 9.44E-08 | 1.34E-05 | 8.74E-13 |
| GO:0048514 | blood vessel morphogenesis | 2.48E-07 | 1.13E-04 | 3.05E-08 |
| GO:0090287 | regulation of cellular response to growth factor stimulus | 8.73E-07 | 5.94E-04 | 2.74E-07 |
| GO:0032501 | multicellular organismal process | 9.60E-07 | 2.06E-06 | 2.20E-19 |
| GO:0035295 | tube development | 1.44E-06 | 5.42E-05 | 2.37E-11 |
| GO:0060429 | epithelium development | 1.53E-06 | 2.58E-05 | 8.96E-13 |
| GO:0060560 | developmental growth involved in morphogenesis | 1.61E-06 | 5.95E-04 | 4.43E-06 |
| GO:0048589 | developmental growth | 1.77E-06 | 2.33E-05 | 4.98E-09 |
| GO:0048468 | cell development | 1.91E-06 | 8.23E-07 | 1.82E-13 |
| GO:0045778 | positive regulation of ossification | 2.30E-06 | 1.45E-03 | 1.20E-04 |
| GO:0033077 | T cell differentiation in thymus | 3.28E-06 | 1.24E-03 | 8.37E-06 |
| GO:0045667 | regulation of osteoblast differentiation | 3.73E-06 | 6.06E-04 | 1.80E-04 |
| GO:0035115 | embryonic forelimb morphogenesis | 4.41E-06 | 2.01E-05 | 5.27E-09 |
| GO:0007423 | sensory organ development | 4.59E-06 | 2.18E-06 | 4.85E-11 |
| GO:0035136 | forelimb morphogenesis | 5.16E-06 | 1.21E-04 | 7.83E-09 |
| GO:0030217 | T cell differentiation | 5.56E-06 | 1.44E-03 | 3.53E-05 |
| GO:0040007 | growth | 6.20E-06 | 3.40E-05 | 2.89E-08 |
| GO:0035148 | tube formation | 6.38E-06 | 1.78E-05 | 4.57E-07 |
| GO:0022603 | regulation of anatomical structure morphogenesis | 1.13E-05 | 7.73E-04 | 9.21E-11 |
| GO:0001822 | kidney development | 1.28E-05 | 4.28E-06 | 1.27E-10 |
| GO:0060070 | canonical Wnt signaling pathway | 2.71E-05 | 7.43E-04 | 3.47E-03 |
| GO:0003006 | developmental process involved in reproduction | 2.88E-05 | 5.77E-05 | 8.53E-05 |
| GO:0072164 | mesonephric tubule development | 4.37E-05 | 4.38E-04 | 4.00E-07 |
| GO:0001654 | eye development | 5.04E-05 | 1.12E-06 | 4.70E-08 |
| GO:0010648 | negative regulation of cell communication | 7.10E-05 | 1.64E-04 | 2.88E-06 |
| GO:0072163 | mesonephric epithelium development | 7.39E-05 | 6.93E-04 | 1.27E-07 |
| GO:0051271 | negative regulation of cellular component movement | 7.43E-05 | 1.41E-03 | 3.29E-05 |
| GO:0001656 | metanephros development | 8.07E-05 | 1.62E-05 | 2.59E-06 |
| GO:0023057 | negative regulation of signaling | 8.57E-05 | 1.41E-04 | 2.48E-06 |
| GO:0072073 | kidney epithelium development | 8.87E-05 | 9.14E-04 | 9.66E-10 |
| GO:0030111 | regulation of Wnt signaling pathway | 1.25E-04 | 5.18E-06 | 1.93E-03 |
| GO:0048812 | neuron projection morphogenesis | 1.27E-04 | 3.34E-05 | 8.00E-04 |
| GO:0042733 | embryonic digit morphogenesis | 1.35E-04 | 5.05E-05 | 3.91E-10 |
| GO:0032989 | cellular component morphogenesis | 1.41E-04 | 1.61E-06 | 7.41E-05 |
| GO:0048858 | cell projection morphogenesis | 1.42E-04 | 1.91E-05 | 7.95E-04 |
| GO:0007178 | transmembrane receptor protein serine/threonine kinase signaling pathway | 1.49E-04 | 1.02E-03 | 7.00E-05 |
| GO:0001823 | mesonephros development | 1.56E-04 | 4.17E-04 | 6.85E-04 |
| GO:0022414 | reproductive process | 1.61E-04 | 3.28E-04 | 2.26E-04 |
| GO:0009880 | embryonic pattern specification | 1.66E-04 | 4.97E-05 | 3.22E-05 |
| GO:0048568 | embryonic organ development | 1.77E-04 | 5.68E-04 | 5.48E-04 |
| GO:0120039 | plasma membrane bounded cell projection morphogenesis | 1.96E-04 | 5.65E-05 | 5.61E-04 |
| GO:0048588 | developmental cell growth | 1.98E-04 | 4.92E-04 | 2.49E-03 |
| GO:0007409 | axonogenesis | 2.34E-04 | 4.54E-04 | 2.24E-05 |
| GO:0006928 | movement of cell or subcellular component | 2.47E-04 | 7.94E-07 | 1.34E-11 |
| GO:0002062 | chondrocyte differentiation | 2.49E-04 | 1.12E-03 | 3.22E-05 |
| GO:0009968 | negative regulation of signal transduction | 3.29E-04 | 1.13E-04 | 2.15E-05 |
| GO:2000027 | regulation of organ morphogenesis | 3.37E-04 | 1.49E-03 | 7.59E-14 |
| GO:0048585 | negative regulation of response to stimulus | 3.59E-04 | 6.32E-05 | 3.54E-05 |
| GO:0003279 | cardiac septum development | 4.12E-04 | 4.96E-04 | 9.07E-05 |
| GO:0010718 | positive regulation of epithelial to mesenchymal transition | 4.12E-04 | 3.65E-04 | 6.13E-06 |
| GO:0010463 | mesenchymal cell proliferation | 4.62E-04 | 3.93E-04 | 3.17E-06 |
| GO:0048762 | mesenchymal cell differentiation | 5.32E-04 | 1.45E-03 | 3.58E-06 |
| GO:0043010 | camera-type eye development | 5.66E-04 | 1.80E-04 | 1.57E-05 |
| GO:0060272 | embryonic skeletal joint morphogenesis | 5.75E-04 | 1.07E-04 | 3.67E-07 |
| GO:0060828 | regulation of canonical Wnt signaling pathway | 6.59E-04 | 3.20E-05 | 4.08E-03 |
| GO:0048736 | appendage development | 6.78E-04 | 6.47E-05 | 7.24E-04 |
| GO:0060173 | limb development | 6.78E-04 | 6.47E-05 | 7.24E-04 |
| GO:1901214 | regulation of neuron death | 1.20E-03 | 7.53E-04 | 1.25E-04 |
| GO:0030279 | negative regulation of ossification | 1.38E-03 | 2.72E-04 | 7.56E-05 |
| GO:0046483 | heterocycle metabolic process | 1.42E-03 | 1.37E-03 | 1.45E-04 |
| GO:0010721 | negative regulation of cell development | 1.52E-03 | 3.83E-04 | 2.99E-06 |
| GO:0060441 | epithelial tube branching involved in lung morphogenesis | 1.59E-03 | 7.97E-04 | 5.91E-04 |
| GO:0050679 | positive regulation of epithelial cell proliferation | 2.16E-03 | 3.38E-04 | 2.24E-05 |
| GO:0030178 | negative regulation of Wnt signaling pathway | 2.32E-03 | 4.18E-04 | 9.01E-05 |
| GO:0072111 | cell proliferation involved in kidney development | 2.50E-03 | 2.45E-04 | 5.15E-05 |
| GO:0051961 | negative regulation of nervous system development | 3.19E-03 | 1.12E-03 | 7.88E-04 |

**Table S24. Commonly enriched KEGG pathways of genes in the highly conserved genomic regions (HCRs) of Strigiformes, Accipitriformes, and Falconiformes.** KEGG pathways also enriched in the HCRs of Passeriformes were excluded. *P*-value was calculated by Fisher’s exact test with a 5% FDR criterion.

| KEGG | Description | *P*-value in Strigiformes | *P*-value in Accipitriformes | *P*-value in Falconiformes |
| --- | --- | --- | --- | --- |
| ko04360 | Axon guidance | 1.88E-06 | 2.20E-05 | 1.86E-05 |
| ko04550 | Signaling pathways regulating pluripotency of stem cells | 5.11E-06 | 2.18E-05 | 3.47E-05 |
| ko05224 | Breast cancer | 9.93E-06 | 3.16E-04 | 1.93E-05 |
| ko04310 | Wnt signaling pathway | 1.92E-05 | 2.18E-05 | 1.06E-04 |
| ko05202 | Transcriptional misregulation in cancer | 2.39E-05 | 3.15E-04 | 2.35E-07 |
| ko05225 | Hepatocellular carcinoma | 3.45E-05 | 6.22E-06 | 3.21E-06 |
| ko05226 | Gastric cancer | 3.60E-05 | 3.33E-05 | 1.44E-05 |
| ko05165 | Human papillomavirus infection | 4.28E-05 | 3.30E-04 | 8.17E-04 |
| ko05217 | Basal cell carcinoma | 6.95E-05 | 8.15E-04 | 3.11E-06 |
| ko05205 | Proteoglycans in cancer | 1.25E-03 | 1.67E-03 | 6.08E-05 |
| ko05221 | Acute myeloid leukemia | 2.79E-03 | 2.29E-04 | 1.19E-03 |
| ko04320 | Dorso-ventral axis formation | 3.06E-03 | 2.62E-04 | 1.18E-03 |

**Table S25. Strigiformes specific GO enrichment of genes in the highly conserved genomic regions (HCRs).** *P*-value was calculated by Fisher’s exact test.

| GO term | Description | *P*-value | FDR |
| --- | --- | --- | --- |
| GO:0032270 | positive regulation of cellular protein metabolic process | 8.18E-07 | 4.47E-05 |
| GO:0051247 | positive regulation of protein metabolic process | 1.96E-06 | 9.49E-05 |
| GO:0032268 | regulation of cellular protein metabolic process | 1.07E-05 | 4.36E-04 |
| GO:0051246 | regulation of protein metabolic process | 1.34E-05 | 5.32E-04 |
| GO:0031401 | positive regulation of protein modification process | 2.12E-05 | 8.12E-04 |
| GO:0007498 | mesoderm development | 4.28E-05 | 1.55E-03 |
| GO:0034483 | heparan sulfate sulfotransferase activity | 4.79E-05 | 1.72E-03 |
| GO:0046651 | lymphocyte proliferation | 4.93E-05 | 1.74E-03 |
| GO:0032943 | mononuclear cell proliferation | 4.93E-05 | 1.74E-03 |
| GO:0010942 | positive regulation of cell death | 6.09E-05 | 2.08E-03 |
| GO:0043068 | positive regulation of programmed cell death | 1.24E-04 | 3.87E-03 |
| GO:0030501 | positive regulation of bone mineralization | 1.45E-04 | 4.34E-03 |
| GO:0043065 | positive regulation of apoptotic process | 1.60E-04 | 4.68E-03 |
| GO:0070169 | positive regulation of biomineral tissue development | 2.49E-04 | 6.82E-03 |
| GO:0070372 | regulation of ERK1 and ERK2 cascade | 2.93E-04 | 7.92E-03 |
| GO:0043408 | regulation of MAPK cascade | 3.01E-04 | 8.10E-03 |
| GO:0001047 | core promoter binding | 3.36E-04 | 8.83E-03 |
| GO:0048645 | animal organ formation | 3.50E-04 | 9.04E-03 |
| GO:0010894 | negative regulation of steroid biosynthetic process | 3.50E-04 | 9.04E-03 |
| GO:0045939 | negative regulation of steroid metabolic process | 3.50E-04 | 9.04E-03 |
| GO:1905208 | negative regulation of cardiocyte differentiation | 5.09E-04 | 1.23E-02 |
| GO:0043589 | skin morphogenesis | 5.09E-04 | 1.23E-02 |
| GO:0010596 | negative regulation of endothelial cell migration | 5.63E-04 | 1.35E-02 |
| GO:0045668 | negative regulation of osteoblast differentiation | 5.63E-04 | 1.35E-02 |
| GO:0031399 | regulation of protein modification process | 5.77E-04 | 1.37E-02 |
| GO:0045669 | positive regulation of osteoblast differentiation | 6.51E-04 | 1.50E-02 |
| GO:1990138 | neuron projection extension | 6.78E-04 | 1.54E-02 |
| GO:0010633 | negative regulation of epithelial cell migration | 6.78E-04 | 1.54E-02 |
| GO:0008201 | heparin binding | 6.94E-04 | 1.56E-02 |
| GO:1901343 | negative regulation of vasculature development | 7.80E-04 | 1.73E-02 |
| GO:0060324 | face development | 7.88E-04 | 1.73E-02 |
| GO:0045687 | positive regulation of glial cell differentiation | 7.88E-04 | 1.73E-02 |
| GO:0060740 | prostate gland epithelium morphogenesis | 7.88E-04 | 1.73E-02 |
| GO:0006090 | pyruvate metabolic process | 8.47E-04 | 1.84E-02 |
| GO:0030010 | establishment of cell polarity | 8.83E-04 | 1.89E-02 |
| GO:0000983 | transcription factor activity, RNA polymerase II core promoter sequence-specific DNA binding | 8.97E-04 | 1.91E-02 |
| GO:0001889 | liver development | 9.70E-04 | 2.04E-02 |
| GO:0051101 | regulation of DNA binding | 9.81E-04 | 2.05E-02 |
| GO:0010975 | regulation of neuron projection development | 1.05E-03 | 2.17E-02 |
| GO:0071542 | dopaminergic neuron differentiation | 1.07E-03 | 2.20E-02 |
| GO:0044451 | nucleoplasm part | 1.17E-03 | 2.37E-02 |
| GO:0016049 | cell growth | 1.17E-03 | 2.37E-02 |
| GO:0019216 | regulation of lipid metabolic process | 1.21E-03 | 2.44E-02 |
| GO:2001234 | negative regulation of apoptotic signaling pathway | 1.41E-03 | 2.78E-02 |
| GO:0050919 | negative chemotaxis | 1.42E-03 | 2.78E-02 |
| GO:0035282 | segmentation | 1.50E-03 | 2.87E-02 |
| GO:0045862 | positive regulation of proteolysis | 1.50E-03 | 2.87E-02 |
| GO:0002718 | regulation of cytokine production involved in immune response | 1.50E-03 | 2.87E-02 |
| GO:0007369 | gastrulation | 1.54E-03 | 2.92E-02 |
| GO:0048640 | negative regulation of developmental growth | 1.55E-03 | 2.92E-02 |
| GO:0001655 | urogenital system development | 1.55E-03 | 2.92E-02 |
| GO:0000790 | nuclear chromatin | 1.60E-03 | 2.98E-02 |
| GO:0003712 | transcription cofactor activity | 1.67E-03 | 3.09E-02 |
| GO:0004714 | transmembrane receptor protein tyrosine kinase activity | 1.69E-03 | 3.11E-02 |
| GO:2000116 | regulation of cysteine-type endopeptidase activity | 1.69E-03 | 3.11E-02 |
| GO:0043281 | regulation of cysteine-type endopeptidase activity involved in apoptotic process | 1.72E-03 | 3.16E-02 |
| GO:0007165 | signal transduction | 1.91E-03 | 3.48E-02 |
| GO:1903322 | positive regulation of protein modification by small protein conjugation or removal | 1.91E-03 | 3.48E-02 |
| GO:0042326 | negative regulation of phosphorylation | 1.98E-03 | 3.59E-02 |
| GO:0045765 | regulation of angiogenesis | 1.99E-03 | 3.60E-02 |
| GO:0022602 | ovulation cycle process | 2.15E-03 | 3.82E-02 |
| GO:0003179 | heart valve morphogenesis | 2.16E-03 | 3.82E-02 |
| GO:0014015 | positive regulation of gliogenesis | 2.16E-03 | 3.82E-02 |
| GO:0060393 | regulation of pathway-restricted SMAD protein phosphorylation | 2.24E-03 | 3.95E-02 |
| GO:0050798 | activated T cell proliferation | 2.32E-03 | 3.95E-02 |
| GO:1902285 | semaphorin-plexin signaling pathway involved in neuron projection guidance | 2.32E-03 | 3.95E-02 |
| GO:0021855 | hypothalamus cell migration | 2.32E-03 | 3.95E-02 |
| GO:0046880 | regulation of follicle-stimulating hormone secretion | 2.32E-03 | 3.95E-02 |
| GO:1901166 | neural crest cell migration involved in autonomic nervous system development | 2.32E-03 | 3.95E-02 |
| GO:0045661 | regulation of myoblast differentiation | 2.32E-03 | 3.95E-02 |
| GO:0061314 | Notch signaling involved in heart development | 2.32E-03 | 3.95E-02 |
| GO:0015015 | heparan sulfate proteoglycan biosynthetic process, enzymatic modification | 2.32E-03 | 3.95E-02 |
| GO:0006189 | 'de novo' IMP biosynthetic process | 2.32E-03 | 3.95E-02 |
| GO:2000726 | negative regulation of cardiac muscle cell differentiation | 2.32E-03 | 3.95E-02 |
| GO:0048714 | positive regulation of oligodendrocyte differentiation | 2.32E-03 | 3.95E-02 |
| GO:0002043 | blood vessel endothelial cell proliferation involved in sprouting angiogenesis | 2.32E-03 | 3.95E-02 |
| GO:0033151 | V(D)J recombination | 2.35E-03 | 3.95E-02 |
| GO:0021675 | nerve development | 2.35E-03 | 3.95E-02 |
| GO:0042100 | B cell proliferation | 2.35E-03 | 3.95E-02 |
| GO:0045577 | regulation of B cell differentiation | 2.35E-03 | 3.95E-02 |
| GO:0001934 | positive regulation of protein phosphorylation | 2.39E-03 | 4.00E-02 |
| GO:0043409 | negative regulation of MAPK cascade | 2.41E-03 | 4.01E-02 |
| GO:0030099 | myeloid cell differentiation | 2.41E-03 | 4.01E-02 |
| GO:0000796 | condensin complex | 2.42E-03 | 4.02E-02 |
| GO:0034770 | histone H4-K20 methylation | 2.50E-03 | 4.02E-02 |
| GO:0021517 | ventral spinal cord development | 2.50E-03 | 4.02E-02 |
| GO:0007221 | positive regulation of transcription of Notch receptor target | 2.50E-03 | 4.02E-02 |
| GO:0034091 | regulation of maintenance of sister chromatid cohesion | 2.50E-03 | 4.02E-02 |
| GO:0003184 | pulmonary valve morphogenesis | 2.50E-03 | 4.02E-02 |
| GO:0021800 | cerebral cortex tangential migration | 2.50E-03 | 4.02E-02 |
| GO:0048617 | embryonic foregut morphogenesis | 2.50E-03 | 4.02E-02 |
| GO:0034182 | regulation of maintenance of mitotic sister chromatid cohesion | 2.50E-03 | 4.02E-02 |
| GO:0003149 | membranous septum morphogenesis | 2.50E-03 | 4.02E-02 |
| GO:0032276 | regulation of gonadotropin secretion | 2.50E-03 | 4.02E-02 |
| GO:0042325 | regulation of phosphorylation | 2.53E-03 | 4.06E-02 |
| GO:0001933 | negative regulation of protein phosphorylation | 2.63E-03 | 4.21E-02 |
| GO:0005021 | vascular endothelial growth factor-activated receptor activity | 2.70E-03 | 4.30E-02 |
| GO:0048511 | rhythmic process | 2.70E-03 | 4.30E-02 |
| GO:0042327 | positive regulation of phosphorylation | 2.71E-03 | 4.30E-02 |
| GO:1905114 | cell surface receptor signaling pathway involved in cell-cell signaling | 2.73E-03 | 4.32E-02 |
| GO:0001932 | regulation of protein phosphorylation | 2.84E-03 | 4.47E-02 |
| GO:0055006 | cardiac cell development | 2.92E-03 | 4.56E-02 |
| GO:0010719 | negative regulation of epithelial to mesenchymal transition | 2.92E-03 | 4.56E-02 |
| GO:0008585 | female gonad development | 2.92E-03 | 4.56E-02 |
| GO:0035265 | organ growth | 2.92E-03 | 4.56E-02 |
| GO:0001756 | somitogenesis | 3.21E-03 | 4.94E-02 |

**Table S26. Strigiformes specifically enriched KEGG pathways of genes in the highly conserved genomic regions (HCRs).** *P*-value was calculated by Fisher’s exact test.

| KEGG | Description | *P*-value | FDR |
| --- | --- | --- | --- |
| ko05210 | Colorectal cancer | 1.01E-03 | 1.85E-02 |
| ko01522 | Endocrine resistance | 3.03E-03 | 4.00E-02 |
| ko05215 | Prostate cancer | 1.19E-03 | 2.00E-02 |
| ko05213 | Endometrial cancer | 1.95E-04 | 4.59E-03 |
| ko04510 | Focal adhesion | 8.97E-04 | 1.85E-02 |
| ko04213 | Longevity regulating pathway - multiple species | 3.91E-03 | 4.76E-02 |
| ko00534 | Glycosaminoglycan biosynthesis - heparan sulfate / heparin | 1.13E-04 | 3.00E-03 |
| ko04916 | Melanogenesis | 6.97E-04 | 1.54E-02 |
| ko04391 | Hippo signaling pathway - fly | 2.77E-03 | 3.94E-02 |
| ko04520 | Adherens junction | 4.17E-05 | 1.37E-03 |

**Table S27. Accipitriformes specific GO enrichment of genes in the highly conserved genomic regions (HCRs).** *P*-value was calculated by Fisher’s exact test.

| GO term | Description | *P*-value | FDR |
| --- | --- | --- | --- |
| GO:0048813 | dendrite morphogenesis | 8.55E-05 | 4.67E-03 |
| GO:0060976 | coronary vasculature development | 1.15E-04 | 5.95E-03 |
| GO:0005515 | protein binding | 1.70E-04 | 8.36E-03 |
| GO:0019321 | pentose metabolic process | 2.45E-04 | 1.15E-02 |
| GO:0046548 | retinal rod cell development | 2.45E-04 | 1.15E-02 |
| GO:0009755 | hormone-mediated signaling pathway | 3.39E-04 | 1.54E-02 |
| GO:0050890 | cognition | 3.40E-04 | 1.54E-02 |
| GO:0007611 | learning or memory | 3.43E-04 | 1.54E-02 |
| GO:0071840 | cellular component organization or biogenesis | 3.52E-04 | 1.57E-02 |
| GO:0005930 | axoneme | 3.54E-04 | 1.57E-02 |
| GO:0043167 | ion binding | 3.64E-04 | 1.61E-02 |
| GO:0046040 | IMP metabolic process | 3.83E-04 | 1.66E-02 |
| GO:0006188 | IMP biosynthetic process | 3.83E-04 | 1.66E-02 |
| GO:0016043 | cellular component organization | 4.21E-04 | 1.78E-02 |
| GO:0030030 | cell projection organization | 7.09E-04 | 2.78E-02 |
| GO:0051056 | regulation of small GTPase mediated signal transduction | 9.69E-04 | 3.65E-02 |
| GO:0014069 | postsynaptic density | 1.22E-03 | 4.37E-02 |
| GO:0099572 | postsynaptic specialization | 1.22E-03 | 4.37E-02 |
| GO:0042462 | eye photoreceptor cell development | 1.25E-03 | 4.43E-02 |
| GO:0042461 | photoreceptor cell development | 1.25E-03 | 4.43E-02 |
| GO:0009168 | purine ribonucleoside monophosphate biosynthetic process | 1.30E-03 | 4.57E-02 |
| GO:0061549 | sympathetic ganglion development | 1.35E-03 | 4.61E-02 |
| GO:0030323 | respiratory tube development | 1.35E-03 | 4.61E-02 |
| GO:0044030 | regulation of DNA methylation | 1.35E-03 | 4.61E-02 |
| GO:0043101 | purine-containing compound salvage | 1.35E-03 | 4.61E-02 |
| GO:0043401 | steroid hormone mediated signaling pathway | 1.45E-03 | 4.79E-02 |

**Table S28. Accipitriformes specifically enriched KEGG pathways of genes in the highly conserved genomic regions (HCRs).** *P*-value was calculated by Fisher’s exact test.

| KEGG | Description | *P*-value | FDR |
| --- | --- | --- | --- |
| ko05220 | Chronic myeloid leukemia | 7.94E-04 | 1.60E-02 |
| ko04152 | AMPK signaling pathway | 2.00E-03 | 3.54E-02 |
| ko00280 | Valine, leucine and isoleucine degradation | 2.29E-03 | 3.86E-02 |
| ko05211 | Renal cell carcinoma | 2.58E-03 | 4.15E-02 |

**Table S29. Falconiformes specific GO enrichment of genes in the highly conserved genomic regions (HCRs).** *P*-value was calculated by Fisher’s exact test. Only GO categories with *P* < 1.00E-03 are shown.

| GO term | Description | *P*-value | FDR |
| --- | --- | --- | --- |
| GO:0051216 | cartilage development | 2.27E-08 | 1.01E-06 |
| GO:0048863 | stem cell differentiation | 2.90E-07 | 1.12E-05 |
| GO:0040008 | regulation of growth | 6.86E-07 | 2.49E-05 |
| GO:0003156 | regulation of animal organ formation | 7.88E-07 | 2.82E-05 |
| GO:2000826 | regulation of heart morphogenesis | 1.14E-06 | 3.99E-05 |
| GO:2001053 | regulation of mesenchymal cell apoptotic process | 1.91E-06 | 6.55E-05 |
| GO:0030856 | regulation of epithelial cell differentiation | 2.78E-06 | 9.14E-05 |
| GO:0048864 | stem cell development | 4.20E-06 | 1.33E-04 |
| GO:0014032 | neural crest cell development | 8.55E-06 | 2.56E-04 |
| GO:0048665 | neuron fate specification | 8.55E-06 | 2.56E-04 |
| GO:0031128 | developmental induction | 9.87E-06 | 2.90E-04 |
| GO:2000736 | regulation of stem cell differentiation | 9.99E-06 | 2.92E-04 |
| GO:0060688 | regulation of morphogenesis of a branching structure | 1.18E-05 | 3.39E-04 |
| GO:0001667 | ameboidal-type cell migration | 1.40E-05 | 3.91E-04 |
| GO:2000738 | positive regulation of stem cell differentiation | 1.42E-05 | 3.95E-04 |
| GO:0048638 | regulation of developmental growth | 1.86E-05 | 5.12E-04 |
| GO:0051149 | positive regulation of muscle cell differentiation | 2.00E-05 | 5.48E-04 |
| GO:0010464 | regulation of mesenchymal cell proliferation | 3.01E-05 | 7.86E-04 |
| GO:0009791 | post-embryonic development | 3.07E-05 | 7.96E-04 |
| GO:0021915 | neural tube development | 3.22E-05 | 8.23E-04 |
| GO:0007507 | heart development | 4.16E-05 | 1.04E-03 |
| GO:1903706 | regulation of hemopoiesis | 4.54E-05 | 1.11E-03 |
| GO:0048557 | embryonic digestive tract morphogenesis | 4.57E-05 | 1.11E-03 |
| GO:2000136 | regulation of cell proliferation involved in heart morphogenesis | 4.57E-05 | 1.11E-03 |
| GO:0072079 | nephron tubule formation | 4.57E-05 | 1.11E-03 |
| GO:0030857 | negative regulation of epithelial cell differentiation | 4.95E-05 | 1.19E-03 |
| GO:0048806 | genitalia development | 4.95E-05 | 1.19E-03 |
| GO:2001054 | negative regulation of mesenchymal cell apoptotic process | 5.15E-05 | 1.23E-03 |
| GO:2000177 | regulation of neural precursor cell proliferation | 6.91E-05 | 1.62E-03 |
| GO:0002053 | positive regulation of mesenchymal cell proliferation | 6.95E-05 | 1.62E-03 |
| GO:1901360 | organic cyclic compound metabolic process | 7.35E-05 | 1.69E-03 |
| GO:1905209 | positive regulation of cardiocyte differentiation | 8.15E-05 | 1.83E-03 |
| GO:0061217 | regulation of mesonephros development | 8.15E-05 | 1.83E-03 |
| GO:1901215 | negative regulation of neuron death | 8.37E-05 | 1.87E-03 |
| GO:0046622 | positive regulation of organ growth | 9.07E-05 | 1.99E-03 |
| GO:0071407 | cellular response to organic cyclic compound | 1.07E-04 | 2.33E-03 |
| GO:0002063 | chondrocyte development | 1.12E-04 | 2.43E-03 |
| GO:0048048 | embryonic eye morphogenesis | 1.12E-04 | 2.43E-03 |
| GO:0046620 | regulation of organ growth | 1.15E-04 | 2.48E-03 |
| GO:0010634 | positive regulation of epithelial cell migration | 1.20E-04 | 2.56E-03 |
| GO:0050680 | negative regulation of epithelial cell proliferation | 1.20E-04 | 2.56E-03 |
| GO:0048103 | somatic stem cell division | 1.23E-04 | 2.59E-03 |
| GO:0036003 | positive regulation of transcription from RNA polymerase II promoter in response to stress | 1.23E-04 | 2.59E-03 |
| GO:2000696 | regulation of epithelial cell differentiation involved in kidney development | 1.23E-04 | 2.59E-03 |
| GO:1905331 | negative regulation of morphogenesis of an epithelium | 1.23E-04 | 2.59E-03 |
| GO:0009798 | axis specification | 1.24E-04 | 2.59E-03 |
| GO:0048839 | inner ear development | 1.35E-04 | 2.81E-03 |
| GO:0048872 | homeostasis of number of cells | 1.41E-04 | 2.92E-03 |
| GO:2000243 | positive regulation of reproductive process | 1.49E-04 | 3.05E-03 |
| GO:0045926 | negative regulation of growth | 1.71E-04 | 3.49E-03 |
| GO:1901213 | regulation of transcription from RNA polymerase II promoter involved in heart development | 1.80E-04 | 3.61E-03 |
| GO:0090185 | negative regulation of kidney development | 1.80E-04 | 3.61E-03 |
| GO:0021904 | dorsal/ventral neural tube patterning | 1.80E-04 | 3.61E-03 |
| GO:0090101 | negative regulation of transmembrane receptor protein serine/threonine kinase signaling pathway | 1.89E-04 | 3.77E-03 |
| GO:0060479 | lung cell differentiation | 2.20E-04 | 4.33E-03 |
| GO:0021954 | central nervous system neuron development | 2.20E-04 | 4.33E-03 |
| GO:0090288 | negative regulation of cellular response to growth factor stimulus | 2.33E-04 | 4.54E-03 |
| GO:0001837 | epithelial to mesenchymal transition | 2.41E-04 | 4.68E-03 |
| GO:0060602 | branch elongation of an epithelium | 2.47E-04 | 4.77E-03 |
| GO:0072182 | regulation of nephron tubule epithelial cell differentiation | 2.47E-04 | 4.77E-03 |
| GO:0035385 | Roundabout signaling pathway | 2.67E-04 | 4.98E-03 |
| GO:0061074 | regulation of neural retina development | 2.67E-04 | 4.98E-03 |
| GO:0036302 | atrioventricular canal development | 2.67E-04 | 4.98E-03 |
| GO:1902866 | regulation of retina development in camera-type eye | 2.67E-04 | 4.98E-03 |
| GO:0021877 | forebrain neuron fate commitment | 2.67E-04 | 4.98E-03 |
| GO:0010453 | regulation of cell fate commitment | 2.85E-04 | 5.21E-03 |
| GO:0048566 | embryonic digestive tract development | 2.85E-04 | 5.21E-03 |
| GO:0060065 | uterus development | 2.85E-04 | 5.21E-03 |
| GO:0035909 | aorta morphogenesis | 2.85E-04 | 5.21E-03 |
| GO:0048713 | regulation of oligodendrocyte differentiation | 2.85E-04 | 5.21E-03 |
| GO:0090189 | regulation of branching involved in ureteric bud morphogenesis | 2.85E-04 | 5.21E-03 |
| GO:0042474 | middle ear morphogenesis | 2.85E-04 | 5.21E-03 |
| GO:0048565 | digestive tract development | 3.37E-04 | 6.04E-03 |
| GO:0014706 | striated muscle tissue development | 3.49E-04 | 6.24E-03 |
| GO:0061005 | cell differentiation involved in kidney development | 3.61E-04 | 6.39E-03 |
| GO:0072001 | renal system development | 3.61E-04 | 6.39E-03 |
| GO:0045992 | negative regulation of embryonic development | 3.61E-04 | 6.39E-03 |
| GO:0017145 | stem cell division | 3.61E-04 | 6.39E-03 |
| GO:0003007 | heart morphogenesis | 3.68E-04 | 6.48E-03 |
| GO:0043542 | endothelial cell migration | 3.68E-04 | 6.48E-03 |
| GO:0005102 | receptor binding | 3.93E-04 | 6.90E-03 |
| GO:0071772 | response to BMP | 3.95E-04 | 6.90E-03 |
| GO:0031369 | translation initiation factor binding | 4.46E-04 | 7.63E-03 |
| GO:2001026 | regulation of endothelial cell chemotaxis | 4.48E-04 | 7.63E-03 |
| GO:0070168 | negative regulation of biomineral tissue development | 4.48E-04 | 7.63E-03 |
| GO:0007399 | nervous system development | 4.53E-04 | 7.70E-03 |
| GO:0034641 | cellular nitrogen compound metabolic process | 5.48E-04 | 9.20E-03 |
| GO:0030858 | positive regulation of epithelial cell differentiation | 5.56E-04 | 9.29E-03 |
| GO:0050867 | positive regulation of cell activation | 5.61E-04 | 9.33E-03 |
| GO:0072132 | mesenchyme morphogenesis | 5.91E-04 | 9.76E-03 |
| GO:0060487 | lung epithelial cell differentiation | 6.72E-04 | 1.09E-02 |
| GO:0021983 | pituitary gland development | 6.72E-04 | 1.09E-02 |
| GO:0048546 | digestive tract morphogenesis | 6.85E-04 | 1.10E-02 |
| GO:2000179 | positive regulation of neural precursor cell proliferation | 6.91E-04 | 1.11E-02 |
| GO:0060349 | bone morphogenesis | 7.00E-04 | 1.12E-02 |
| GO:0008544 | epidermis development | 7.00E-04 | 1.12E-02 |
| GO:0048639 | positive regulation of developmental growth | 7.68E-04 | 1.21E-02 |
| GO:0001525 | angiogenesis | 7.99E-04 | 1.25E-02 |
| GO:0022029 | telencephalon cell migration | 9.17E-04 | 1.41E-02 |
| GO:0072175 | epithelial tube formation | 9.17E-04 | 1.41E-02 |
| GO:0001759 | organ induction | 9.67E-04 | 1.46E-02 |
| GO:1905276 | regulation of epithelial tube formation | 9.67E-04 | 1.46E-02 |
| GO:0048596 | embryonic camera-type eye morphogenesis | 9.67E-04 | 1.46E-02 |
| GO:0060037 | pharyngeal system development | 9.67E-04 | 1.46E-02 |
| GO:0021522 | spinal cord motor neuron differentiation | 9.67E-04 | 1.46E-02 |

**Table S30. Falconiformes specifically enriched KEGG pathways of genes in the highly conserved genomic regions (HCRs).** *P*-value was calculated by Fisher’s exact test.

| KEGG | Description | *P*-value | FDR |
| --- | --- | --- | --- |
| ko04013 | MAPK signaling pathway - fly | 7.31E-04 | 1.92E-02 |
| ko04010 | MAPK signaling pathway | 1.08E-03 | 2.39E-02 |

**Table S31. Passeriformes specific GO enrichment of genes in the highly conserved genomic regions (HCRs).** *P*-value was calculated by Fisher’s exact test.

| GO term | Description | *P*-value | FDR |
| --- | --- | --- | --- |
| GO:0046332 | SMAD binding | 3.50E-05 | 2.71E-03 |
| GO:0008038 | neuron recognition | 3.68E-05 | 2.83E-03 |
| GO:0044446 | intracellular organelle part | 2.33E-04 | 1.50E-02 |
| GO:0071634 | regulation of transforming growth factor beta production | 3.34E-04 | 2.03E-02 |
| GO:2001028 | positive regulation of endothelial cell chemotaxis | 4.92E-04 | 2.85E-02 |
| GO:0060573 | cell fate specification involved in pattern specification | 4.92E-04 | 2.85E-02 |
| GO:0044422 | organelle part | 7.32E-04 | 4.02E-02 |
| GO:0090575 | RNA polymerase II transcription factor complex | 8.86E-04 | 4.79E-02 |

**Table S32. Commonly enriched Gene Ontology (GO) categories of expanded gene families in the common ancestor of Strigiformes, brown kiwi and chuck-will’s widow.** GO categories also enriched in contracted gene families in nocturnal birds were excluded. *P*-value was calculated by Fisher’s exact test with a 5% FDR criterion.

| GO term | Description | *P*-value in the common ancestor of Strigiformes | *P*-value in brown kiwi | *P*-value in chuck- will’s widow |
| --- | --- | --- | --- | --- |
| GO:0050839 | cell adhesion molecule binding | 2.41E-03 | 4.55E-02 | 3.39E-02 |
| GO:0050772 | positive regulation of axonogenesis | 3.49E-03 | 8.37E-03 | 4.32E-03 |
| GO:0031253 | cell projection membrane | 2.55E-03 | 6.72E-03 | 3.01E-02 |
| GO:0070062 | extracellular exosome | 3.05E-04 | 6.80E-03 | 7.83E-03 |
| GO:0046872 | metal ion binding | 3.61E-03 | 5.81E-03 | 7.46E-03 |
| GO:0043547 | positive regulation of GTPase activity | 1.02E-03 | 1.08E-02 | 2.64E-02 |
| GO:0031012 | extracellular matrix | 5.79E-04 | 5.66E-03 | 2.29E-02 |
| GO:0043169 | cation binding | 3.44E-03 | 2.24E-03 | 9.13E-03 |
| GO:0043167 | ion binding | 1.64E-02 | 4.78E-04 | 8.25E-03 |
| GO:0050808 | synapse organization | 4.08E-02 | 2.10E-05 | 1.54E-02 |
| GO:0043087 | regulation of GTPase activity | 8.03E-04 | 1.95E-02 | 3.33E-02 |
| GO:0008360 | regulation of cell shape | 6.71E-04 | 4.86E-03 | 1.08E-02 |
| GO:0045927 | positive regulation of growth | 2.05E-03 | 1.87E-02 | 4.78E-02 |
| GO:0030030 | cell projection organization | 1.94E-05 | 2.49E-02 | 2.30E-02 |
| GO:0045111 | intermediate filament cytoskeleton | 1.20E-02 | 4.00E-02 | 5.64E-05 |
| GO:0031982 | vesicle | 1.43E-04 | 4.93E-03 | 1.25E-02 |
| GO:0005794 | Golgi apparatus | 2.79E-02 | 2.10E-02 | 1.02E-02 |
| GO:0065008 | regulation of biological quality | 7.10E-06 | 8.50E-04 | 1.29E-02 |
| GO:0071495 | cellular response to endogenous stimulus | 3.18E-04 | 4.51E-02 | 6.02E-03 |
| GO:0005886 | plasma membrane | 4.84E-06 | 7.69E-06 | 1.22E-03 |
| GO:0051716 | cellular response to stimulus | 3.92E-04 | 2.85E-02 | 3.60E-02 |
| GO:0044425 | membrane part | 2.67E-04 | 4.92E-06 | 3.93E-02 |
| GO:0044421 | extracellular region part | 3.37E-04 | 1.39E-04 | 3.31E-04 |
| GO:0005509 | calcium ion binding | 4.83E-02 | 8.05E-03 | 1.03E-02 |
| GO:0016020 | membrane | 6.78E-04 | 5.27E-06 | 4.01E-03 |
| GO:0043230 | extracellular organelle | 3.54E-04 | 7.43E-03 | 8.14E-03 |
| GO:0006996 | organelle organization | 1.47E-07 | 2.29E-02 | 2.45E-02 |
| GO:0071840 | cellular component organization or biogenesis | 3.46E-07 | 2.37E-02 | 3.64E-03 |
| GO:0044449 | contractile fiber part | 1.63E-04 | 1.68E-02 | 2.08E-02 |
| GO:0050770 | regulation of axonogenesis | 2.65E-03 | 2.39E-03 | 1.91E-02 |
| GO:0016043 | cellular component organization | 1.87E-07 | 1.83E-02 | 3.10E-03 |
| GO:1903561 | extracellular vesicle | 3.54E-04 | 7.38E-03 | 8.11E-03 |
| GO:0016324 | apical plasma membrane | 2.92E-03 | 1.30E-03 | 3.06E-02 |
| GO:0098590 | plasma membrane region | 4.13E-04 | 3.65E-04 | 1.07E-02 |
| GO:0010770 | positive regulation of cell morphogenesis involved in differentiation | 2.56E-05 | 4.07E-02 | 1.34E-02 |
| GO:0061387 | regulation of extent of cell growth | 9.81E-03 | 1.94E-02 | 7.24E-03 |
| GO:0030516 | regulation of axon extension | 4.75E-03 | 1.00E-02 | 4.52E-03 |

**Table S33. Commonly enriched KEGG pathways of expanded gene families in the common ancestor of Strigiformes, brown kiwi and chuck-will’s widow.** KEGG pathways also enriched in contracted gene families in nocturnal birds were excluded. *P*-value was calculated by Fisher’s exact test with a 5% FDR criterion.

| KEGG | Description | *P*-value in the common ancestor of Strigiformes | *P*-value in brown kiwi | *P*-value in chuck- will’s widow |
| --- | --- | --- | --- | --- |
| ko04976 | Bile secretion | 1.22E-07 | 1.92E-02 | 2.88E-07 |
| ko04015 | Rap1 signaling pathway | 1.79E-03 | 1.89E-03 | 1.06E-04 |

**Table S34. Gene Ontology (GO) enrichment of gene families that were contracted in size the present nocturnal bird species genomes.** *P*-value was calculated by Fisher’s exact test with a 5% FDR criterion.

**a.** High-quality nocturnal bird genomes vs. Diurnal bird genomes

| GO term | Description | Fold-change of Avg. gene number | *P*-value |
| --- | --- | --- | --- |
| GO:0102337 | 3-oxo-cerotoyl-CoA synthase activity | 0.26 | 2.77E-03 |
| GO:0102336 | 3-oxo-arachidoyl-CoA synthase activity | 0.19 | 2.77E-03 |
| GO:0102338 | 3-oxo-lignoceronyl-CoA synthase activity | 0.14 | 2.77E-03 |
| GO:0007252 | I-kappaB phosphorylation | 0.45 | 4.28E-03 |
| GO:0016151 | nickel cation binding | 0.25 | 5.67E-03 |
| GO:0018298 | protein-chromophore linkage | 0 | 7.79E-03 |
| GO:0009583 | detection of light stimulus | 0.23 | 8.15E-03 |
| GO:0009881 | photoreceptor activity | 0 | 8.77E-03 |
| GO:0003999 | adenine phosphoribosyltransferase activity | 0.31 | 1.13E-02 |
| GO:1900238 | regulation of metanephric mesenchymal cell migration by platelet-derived growth factor receptor-beta signaling pathway | 0.25 | 1.16E-02 |
| GO:2000591 | positive regulation of metanephric mesenchymal cell migration | 0.25 | 1.16E-02 |
| GO:0010512 | negative regulation of phosphatidylinositol biosynthetic process | 0.25 | 1.16E-02 |
| GO:0035793 | positive regulation of metanephric mesenchymal cell migration by platelet-derived growth factor receptor-beta signaling pathway | 0.25 | 1.16E-02 |
| GO:0001750 | photoreceptor outer segment | 0.12 | 1.18E-02 |
| GO:0046875 | ephrin receptor binding | 0.39 | 1.25E-02 |
| GO:0003913 | DNA photolyase activity | 0.28 | 1.69E-02 |
| GO:0046083 | adenine metabolic process | 0.31 | 1.73E-02 |
| GO:0043096 | purine nucleobase salvage | 0.31 | 1.73E-02 |
| GO:0046084 | adenine biosynthetic process | 0.31 | 1.73E-02 |
| GO:0006168 | adenine salvage | 0.31 | 1.73E-02 |
| GO:2000589 | regulation of metanephric mesenchymal cell migration | 0.25 | 1.73E-02 |
| GO:0044613 | nuclear pore central transport channel | 0.45 | 1.83E-02 |
| GO:0006323 | DNA packaging | 0.29 | 2.04E-02 |
| GO:0050819 | negative regulation of coagulation | 0.38 | 2.15E-02 |
| GO:0031952 | regulation of protein autophosphorylation | 0.38 | 2.15E-02 |
| GO:0004300 | enoyl-CoA hydratase activity | 0 | 2.25E-02 |
| GO:0006384 | transcription initiation from RNA polymerase III promoter | 0.31 | 2.30E-02 |
| GO:0007602 | phototransduction | 0 | 2.36E-02 |
| GO:0009328 | phenylalanine-tRNA ligase complex | 0.42 | 2.43E-02 |
| GO:0032722 | positive regulation of chemokine production | 0.48 | 2.69E-02 |
| GO:0045356 | positive regulation of interferon-alpha biosynthetic process | 0.48 | 2.87E-02 |
| GO:0006432 | phenylalanyl-tRNA aminoacylation | 0.42 | 2.87E-02 |
| GO:0009108 | coenzyme biosynthetic process | 0.42 | 2.87E-02 |
| GO:0048269 | methionine adenosyltransferase complex | 0.42 | 3.02E-02 |
| GO:0004826 | phenylalanine-tRNA ligase activity | 0.42 | 3.35E-02 |
| GO:0009931 | calcium-dependent protein serine/threonine kinase activity | 0.44 | 3.35E-02 |
| GO:0004478 | methionine adenosyltransferase activity | 0.42 | 3.35E-02 |
| GO:0061178 | regulation of insulin secretion involved in cellular response to glucose stimulus | 0.43 | 3.41E-02 |
| GO:0034154 | toll-like receptor 7 signaling pathway | 0.48 | 3.43E-02 |
| GO:0045354 | regulation of interferon-alpha biosynthetic process | 0.48 | 3.43E-02 |
| GO:0035092 | sperm chromatin condensation | 0.29 | 3.43E-02 |
| GO:0045416 | positive regulation of interleukin-8 biosynthetic process | 0.48 | 3.43E-02 |
| GO:0010511 | regulation of phosphatidylinositol biosynthetic process | 0.25 | 3.43E-02 |
| GO:0042599 | lamellar body | 0 | 3.62E-02 |
| GO:0008385 | IkappaB kinase complex | 0.45 | 3.62E-02 |
| GO:0045078 | positive regulation of interferon-gamma biosynthetic process | 0.48 | 3.99E-02 |
| GO:0071372 | cellular response to follicle-stimulating hormone stimulus | 0.43 | 3.99E-02 |
| GO:0045359 | positive regulation of interferon-beta biosynthetic process | 0.48 | 3.99E-02 |
| GO:0035791 | platelet-derived growth factor receptor-beta signaling pathway | 0.25 | 3.99E-02 |
| GO:0045414 | regulation of interleukin-8 biosynthetic process | 0.48 | 3.99E-02 |
| GO:0006556 | S-adenosylmethionine biosynthetic process | 0.42 | 3.99E-02 |
| GO:0009582 | detection of abiotic stimulus | 0.12 | 4.31E-02 |
| GO:0010857 | calcium-dependent protein kinase activity | 0.44 | 4.45E-02 |
| GO:0048013 | ephrin receptor signaling pathway | 0.39 | 4.46E-02 |
| GO:0009581 | detection of external stimulus | 0.08 | 4.47E-02 |
| GO:0045357 | regulation of interferon-beta biosynthetic process | 0.48 | 4.55E-02 |
| GO:0071072 | negative regulation of phospholipid biosynthetic process | 0.25 | 4.55E-02 |
| GO:0045335 | phagocytic vesicle | 0.48 | 4.92E-02 |
| GO:0035197 | siRNA binding | 0.48 | 4.99E-02 |

**b.** All nocturnal bird genomes including low-quality genomes vs. Diurnal bird genomes

| GO term | Description | Fold-change of Avg. gene number | *P*-value |
| --- | --- | --- | --- |
| GO:0036093 | germ cell proliferation | 0.49 | 9.16E-03 |
| GO:0002176 | male germ cell proliferation | 0.47 | 9.16E-03 |
| GO:0097058 | CRLF-CLCF1 complex | 0.42 | 1.02E-02 |
| GO:0004300 | enoyl-CoA hydratase activity | 0.33 | 1.57E-02 |
| GO:0006384 | transcription initiation from RNA polymerase III promoter | 0.46 | 1.76E-02 |
| GO:0030656 | regulation of vitamin metabolic process | 0.43 | 2.09E-02 |
| GO:0060556 | regulation of vitamin D biosynthetic process | 0.42 | 2.09E-02 |
| GO:0006896 | Golgi to vacuole transport | 0.49 | 2.68E-02 |
| GO:0070120 | ciliary neurotrophic factor-mediated signaling pathway | 0.37 | 2.83E-02 |
| GO:0003360 | brainstem development | 0.48 | 2.83E-02 |
| GO:0048699 | generation of neurons | 0.45 | 3.34E-02 |
| GO:0004064 | arylesterase activity | 0.44 | 3.65E-02 |
| GO:0005127 | ciliary neurotrophic factor receptor binding | 0.42 | 3.65E-02 |
| GO:0070561 | vitamin D receptor signaling pathway | 0.42 | 4.09E-02 |
| GO:0008582 | regulation of synaptic growth at neuromuscular junction | 0.48 | 4.09E-02 |
| GO:0048680 | positive regulation of axon regeneration | 0.46 | 4.09E-02 |
| GO:0070572 | positive regulation of neuron projection regeneration | 0.45 | 4.09E-02 |
| GO:0042765 | GPI-anchor transamidase complex | 0.48 | 4.54E-02 |
| GO:0034388 | Pwp2p-containing subcomplex of 90S preribosome | 0.48 | 4.54E-02 |
| GO:2000810 | regulation of bicellular tight junction assembly | 0.45 | 4.88E-02 |

**Table S35. Gene Ontology (GO) enrichment of gene families that were expanded in the present nocturnal bird species genomes.** *P*-value was calculated by Fisher’s exact test with a 5% FDR criterion.

**a.** High-quality nocturnal bird genomes vs. Diurnal bird genomes

| GO term | Description | Fold-change of Avg. gene number | *P*-value |
| --- | --- | --- | --- |
| GO:0043167 | ion binding | 2.13 | 2.17E-07 |
| GO:0046872 | metal ion binding | 2.12 | 1.17E-05 |
| GO:0043169 | cation binding | 2.15 | 2.00E-05 |
| GO:0070938 | contractile ring | 2.32 | 2.47E-05 |
| GO:0019219 | regulation of nucleobase-containing compound metabolic process | 2.19 | 2.71E-05 |
| GO:0051252 | regulation of RNA metabolic process | 2.19 | 5.53E-05 |
| GO:0010556 | regulation of macromolecule biosynthetic process | 2.22 | 5.81E-05 |
| GO:0045892 | negative regulation of transcription, DNA-templated | 2.15 | 5.95E-05 |
| GO:0060976 | coronary vasculature development | 2.01 | 6.16E-05 |
| GO:0031344 | regulation of cell projection organization | 2.05 | 6.65E-05 |
| GO:2000113 | negative regulation of cellular macromolecule biosynthetic process | 2.2 | 6.88E-05 |
| GO:0006355 | regulation of transcription, DNA-templated | 2.23 | 7.22E-05 |
| GO:0010558 | negative regulation of macromolecule biosynthetic process | 2.23 | 7.33E-05 |
| GO:1902679 | negative regulation of RNA biosynthetic process | 2.23 | 7.53E-05 |
| GO:1903507 | negative regulation of nucleic acid-templated transcription | 2.23 | 7.53E-05 |
| GO:1903506 | regulation of nucleic acid-templated transcription | 2.24 | 8.23E-05 |
| GO:0080090 | regulation of primary metabolic process | 2.13 | 8.50E-05 |
| GO:2001141 | regulation of RNA biosynthetic process | 2.25 | 8.61E-05 |
| GO:0032502 | developmental process | 2.15 | 8.99E-05 |
| GO:0006351 | transcription, DNA-templated | 2.24 | 9.07E-05 |

**b.** All nocturnal bird genomes including low-quality genomes vs. Diurnal bird genomes

| GO term | Description | Fold-change of Avg. gene number | *P*-value |
| --- | --- | --- | --- |
| GO:0043167 | ion binding | 2.82 | 1.98E-06 |
| GO:0005524 | ATP binding | 2.57 | 3.72E-06 |
| GO:0032559 | adenyl ribonucleotide binding | 2.64 | 4.46E-06 |
| GO:0030554 | adenyl nucleotide binding | 2.66 | 5.09E-06 |
| GO:0036094 | small molecule binding | 2.7 | 5.53E-06 |
| GO:0043168 | anion binding | 2.73 | 7.19E-06 |
| GO:0016773 | phosphotransferase activity, alcohol group as acceptor | 2.75 | 2.29E-05 |
| GO:0016301 | kinase activity | 2.96 | 4.32E-05 |
| GO:0000166 | nucleotide binding | 2.73 | 7.86E-05 |
| GO:1901265 | nucleoside phosphate binding | 2.74 | 7.86E-05 |
| GO:0006793 | phosphorus metabolic process | 2.74 | 9.37E-05 |
| GO:0004672 | protein kinase activity | 2.91 | 9.75E-05 |
| GO:0014069 | postsynaptic density | 2.69 | 1.03E-04 |
| GO:0099572 | postsynaptic specialization | 2.65 | 1.03E-04 |
| GO:0004714 | transmembrane receptor protein tyrosine kinase activity | 65 | 1.05E-04 |
| GO:0035639 | purine ribonucleoside triphosphate binding | 2.82 | 1.70E-04 |
| GO:0032555 | purine ribonucleotide binding | 2.88 | 2.08E-04 |
| GO:0007167 | enzyme linked receptor protein signaling pathway | 3.61 | 2.29E-04 |
| GO:0007169 | transmembrane receptor protein tyrosine kinase signaling pathway | 3.89 | 2.42E-04 |
| GO:0017076 | purine nucleotide binding | 2.94 | 2.48E-04 |
| GO:0032553 | ribonucleotide binding | 2.97 | 2.80E-04 |

**Table S36. Gene Ontology (GO) enrichment of PSGs that were shared in two and/or three nocturnal bird groups.** *P*-value was calculated by Fisher’s exact test.

| GO term | Description | *P*-value | *FDR* |
| --- | --- | --- | --- |
| GO:1990823 | response to leukemia inhibitory factor | 3.98E-03 | 2.67E-01 |
| GO:1990830 | cellular response to leukemia inhibitory factor | 3.98E-03 | 2.67E-01 |
| GO:0070129 | regulation of mitochondrial translation | 5.38E-03 | 2.67E-01 |
| GO:0008406 | gonad development | 5.86E-03 | 2.67E-01 |
| GO:0035025 | positive regulation of Rho protein signal transduction | 7.94E-03 | 2.67E-01 |
| GO:0007416 | synapse assembly | 1.09E-02 | 2.67E-01 |
| GO:0007283 | spermatogenesis | 1.43E-02 | 2.67E-01 |
| GO:0048232 | male gamete generation | 1.43E-02 | 2.67E-01 |
| GO:0007098 | centrosome cycle | 1.44E-02 | 2.67E-01 |
| GO:0022414 | reproductive process | 1.72E-02 | 2.67E-01 |
| GO:0043588 | skin development | 1.82E-02 | 2.67E-01 |
| GO:0008584 | male gonad development | 1.82E-02 | 2.67E-01 |
| GO:0006614 | SRP-dependent cotranslational protein targeting to membrane | 1.82E-02 | 2.67E-01 |
| GO:0007276 | gamete generation | 1.98E-02 | 2.67E-01 |
| GO:0034097 | response to cytokine | 2.08E-02 | 2.67E-01 |
| GO:0046579 | positive regulation of Ras protein signal transduction | 2.24E-02 | 2.67E-01 |
| GO:0006613 | cotranslational protein targeting to membrane | 2.24E-02 | 2.67E-01 |
| GO:0051301 | cell division | 2.30E-02 | 2.67E-01 |
| GO:0038202 | TORC1 signaling | 2.39E-02 | 2.67E-01 |
| GO:0090646 | mitochondrial tRNA processing | 2.39E-02 | 2.67E-01 |
| GO:0050832 | defense response to fungus | 2.39E-02 | 2.67E-01 |
| GO:0006590 | thyroid hormone generation | 2.39E-02 | 2.67E-01 |
| GO:1900483 | regulation of protein targeting to vacuolar membrane | 2.39E-02 | 2.67E-01 |
| GO:0042088 | T-helper 1 type immune response | 2.39E-02 | 2.67E-01 |
| GO:0033148 | positive regulation of intracellular estrogen receptor signaling pathway | 2.39E-02 | 2.67E-01 |
| GO:0030309 | poly-N-acetyllactosamine metabolic process | 2.39E-02 | 2.67E-01 |
| GO:0010626 | negative regulation of Schwann cell proliferation | 2.39E-02 | 2.67E-01 |
| GO:0010624 | regulation of Schwann cell proliferation | 2.39E-02 | 2.67E-01 |
| GO:0043308 | eosinophil degranulation | 2.39E-02 | 2.67E-01 |
| GO:0043307 | eosinophil activation | 2.39E-02 | 2.67E-01 |
| GO:0032817 | regulation of natural killer cell proliferation | 2.39E-02 | 2.67E-01 |
| GO:0032819 | positive regulation of natural killer cell proliferation | 2.39E-02 | 2.67E-01 |
| GO:0035493 | SNARE complex assembly | 2.39E-02 | 2.67E-01 |
| GO:1902187 | negative regulation of viral release from host cell | 2.39E-02 | 2.67E-01 |
| GO:0000964 | mitochondrial RNA 5'-end processing | 2.39E-02 | 2.67E-01 |
| GO:0000963 | mitochondrial RNA processing | 2.39E-02 | 2.67E-01 |
| GO:0000966 | RNA 5'-end processing | 2.39E-02 | 2.67E-01 |
| GO:0051140 | regulation of NK T cell proliferation | 2.39E-02 | 2.67E-01 |
| GO:0051142 | positive regulation of NK T cell proliferation | 2.39E-02 | 2.67E-01 |
| GO:0045900 | negative regulation of translational elongation | 2.39E-02 | 2.67E-01 |
| GO:0032725 | positive regulation of granulocyte macrophage colony-stimulating factor production | 2.39E-02 | 2.67E-01 |
| GO:1902713 | regulation of interferon-gamma secretion | 2.39E-02 | 2.67E-01 |
| GO:1902715 | positive regulation of interferon-gamma secretion | 2.39E-02 | 2.67E-01 |
| GO:0002278 | eosinophil activation involved in immune response | 2.39E-02 | 2.67E-01 |
| GO:0016576 | histone dephosphorylation | 2.39E-02 | 2.67E-01 |
| GO:0046884 | follicle-stimulating hormone secretion | 2.39E-02 | 2.67E-01 |
| GO:0071474 | cellular hyperosmotic response | 2.39E-02 | 2.67E-01 |
| GO:0031204 | posttranslational protein targeting to membrane, translocation | 2.39E-02 | 2.67E-01 |
| GO:0030311 | poly-N-acetyllactosamine biosynthetic process | 2.39E-02 | 2.67E-01 |
| GO:0060253 | negative regulation of glial cell proliferation | 2.39E-02 | 2.67E-01 |
| GO:0032645 | regulation of granulocyte macrophage colony-stimulating factor production | 2.39E-02 | 2.67E-01 |
| GO:0098928 | presynaptic signal transduction | 2.39E-02 | 2.67E-01 |
| GO:0051133 | regulation of NK T cell activation | 2.39E-02 | 2.67E-01 |
| GO:0051135 | positive regulation of NK T cell activation | 2.39E-02 | 2.67E-01 |
| GO:1903564 | regulation of protein localization to cilium | 2.39E-02 | 2.67E-01 |
| GO:0099526 | presynapse to nucleus signaling pathway | 2.39E-02 | 2.67E-01 |
| GO:0032740 | positive regulation of interleukin-17 production | 2.39E-02 | 2.67E-01 |
| GO:0043520 | regulation of myosin II filament assembly | 2.39E-02 | 2.67E-01 |
| GO:1903568 | negative regulation of protein localization to ciliary membrane | 2.39E-02 | 2.67E-01 |
| GO:0043519 | regulation of myosin II filament organization | 2.39E-02 | 2.67E-01 |
| GO:1903567 | regulation of protein localization to ciliary membrane | 2.39E-02 | 2.67E-01 |
| GO:1903565 | negative regulation of protein localization to cilium | 2.39E-02 | 2.67E-01 |
| GO:0042060 | wound healing | 2.69E-02 | 2.71E-01 |
| GO:0051057 | positive regulation of small GTPase mediated signal transduction | 2.69E-02 | 2.71E-01 |
| GO:0007131 | reciprocal meiotic recombination | 2.69E-02 | 2.71E-01 |
| GO:0035825 | homologous recombination | 2.69E-02 | 2.71E-01 |
| GO:0006310 | DNA recombination | 2.83E-02 | 2.73E-01 |
| GO:0098840 | protein transport along microtubule | 3.18E-02 | 2.73E-01 |
| GO:0099118 | microtubule-based protein transport | 3.18E-02 | 2.73E-01 |
| GO:0072599 | establishment of protein localization to endoplasmic reticulum | 3.18E-02 | 2.73E-01 |
| GO:0042073 | intraciliary transport | 3.18E-02 | 2.73E-01 |
| GO:0045047 | protein targeting to ER | 3.18E-02 | 2.73E-01 |
| GO:0048609 | multicellular organismal reproductive process | 3.22E-02 | 2.73E-01 |
| GO:0035023 | regulation of Rho protein signal transduction | 3.35E-02 | 2.73E-01 |
| GO:0042102 | positive regulation of T cell proliferation | 3.71E-02 | 2.73E-01 |
| GO:0071345 | cellular response to cytokine stimulus | 4.09E-02 | 2.73E-01 |
| GO:0051225 | spindle assembly | 4.26E-02 | 2.73E-01 |
| GO:0070972 | protein localization to endoplasmic reticulum | 4.26E-02 | 2.73E-01 |
| GO:0031023 | microtubule organizing center organization | 4.26E-02 | 2.73E-01 |
| GO:1903046 | meiotic cell cycle process | 4.28E-02 | 2.73E-01 |
| GO:0048608 | reproductive structure development | 4.28E-02 | 2.73E-01 |
| GO:0006972 | hyperosmotic response | 4.72E-02 | 2.73E-01 |
| GO:0032926 | negative regulation of activin receptor signaling pathway | 4.72E-02 | 2.73E-01 |
| GO:1900060 | negative regulation of ceramide biosynthetic process | 4.72E-02 | 2.73E-01 |
| GO:0032729 | positive regulation of interferon-gamma production | 4.72E-02 | 2.73E-01 |
| GO:0097286 | iron ion import | 4.72E-02 | 2.73E-01 |
| GO:1902525 | regulation of protein monoubiquitination | 4.72E-02 | 2.73E-01 |
| GO:0043306 | positive regulation of mast cell degranulation | 4.72E-02 | 2.73E-01 |
| GO:0002888 | positive regulation of myeloid leukocyte mediated immunity | 4.72E-02 | 2.73E-01 |
| GO:0051968 | positive regulation of synaptic transmission, glutamatergic | 4.72E-02 | 2.73E-01 |
| GO:0032660 | regulation of interleukin-17 production | 4.72E-02 | 2.73E-01 |
| GO:1902916 | positive regulation of protein polyubiquitination | 4.72E-02 | 2.73E-01 |
| GO:0051382 | kinetochore assembly | 4.72E-02 | 2.73E-01 |
| GO:1903595 | positive regulation of histamine secretion by mast cell | 4.72E-02 | 2.73E-01 |
| GO:0035418 | protein localization to synapse | 4.72E-02 | 2.73E-01 |
| GO:0045176 | apical protein localization | 4.72E-02 | 2.73E-01 |
| GO:1903593 | regulation of histamine secretion by mast cell | 4.72E-02 | 2.73E-01 |
| GO:0050910 | detection of mechanical stimulus involved in sensory perception of sound | 4.72E-02 | 2.73E-01 |
| GO:1905342 | positive regulation of protein localization to kinetochore | 4.72E-02 | 2.73E-01 |
| GO:0032925 | regulation of activin receptor signaling pathway | 4.72E-02 | 2.73E-01 |
| GO:0060005 | vestibular reflex | 4.72E-02 | 2.73E-01 |
| GO:0032274 | gonadotropin secretion | 4.72E-02 | 2.73E-01 |
| GO:0033008 | positive regulation of mast cell activation involved in immune response | 4.72E-02 | 2.73E-01 |
| GO:0033692 | cellular polysaccharide biosynthetic process | 4.72E-02 | 2.73E-01 |
| GO:1904424 | regulation of GTP binding | 4.72E-02 | 2.73E-01 |
| GO:0035655 | interleukin-18-mediated signaling pathway | 4.72E-02 | 2.73E-01 |
| GO:0014902 | myotube differentiation | 4.72E-02 | 2.73E-01 |
| GO:0046597 | negative regulation of viral entry into host cell | 4.72E-02 | 2.73E-01 |
| GO:0002323 | natural killer cell activation involved in immune response | 4.72E-02 | 2.73E-01 |
| GO:0090155 | negative regulation of sphingolipid biosynthetic process | 4.72E-02 | 2.73E-01 |
| GO:0090156 | cellular sphingolipid homeostasis | 4.72E-02 | 2.73E-01 |
| GO:0046641 | positive regulation of alpha-beta T cell proliferation | 4.72E-02 | 2.73E-01 |
| GO:0043320 | natural killer cell degranulation | 4.72E-02 | 2.73E-01 |
| GO:1902914 | regulation of protein polyubiquitination | 4.72E-02 | 2.73E-01 |
| GO:1905340 | regulation of protein localization to kinetochore | 4.72E-02 | 2.73E-01 |
| GO:0032275 | luteinizing hormone secretion | 4.72E-02 | 2.73E-01 |
| GO:0070131 | positive regulation of mitochondrial translation | 4.72E-02 | 2.73E-01 |
| GO:1903335 | regulation of vacuolar transport | 4.72E-02 | 2.73E-01 |
| GO:0006620 | posttranslational protein targeting to endoplasmic reticulum membrane | 4.72E-02 | 2.73E-01 |
| GO:0002573 | myeloid leukocyte differentiation | 4.84E-02 | 2.73E-01 |

**Table S37. List of genes showing accelerated *d_N_*/*d_S_* in the nocturnal birds**

| Gene | ω Nocturnal birds branches | | | |  | ω outgroups |
| --- | --- | --- | --- | --- | --- | --- |
|  | The ancestral branch of Strigiformes | Chuck-will’s-widow | Brown kiwi | Merged | |  |
| *RDH8* | 0.4098 | 0.2631 | 0.4149 | 0.3573 | | 0.0630 |
| *PDE6B* | 0.2270 | 0.2128 | 0.1369 | 0.1720 | | 0.0579 |
| *LOC100858777* | 0.5158 | 0.2636 | 0.6814 | 0.4287 | | 0.1009 |
| *PPEF2* | 0.2939 | 0.3382 | 0.2188 | 0.2936 | | 0.1081 |
| *SLC24A1* | 1.8907 | 0.7179 | 0.6050 | 0.7305 | | 0.2656 |
| *LOC770429* | 1.8810 | 0.4821 | 0.4012 | 0.5092 | | 0.1971 |
| *ACTN1* | 0.4484 | 0.0408 | 0.0320 | 0.1136 | | 0.0226 |
| *CPLX4* | 2.0519 | 1.5421 | 0.1849 | 0.5086 | | 0.0671 |
| *GRK7* | 1.2989 | 0.7244 | 0.2820 | 0.5997 | | 0.2282 |
| *GBE* | 0.5387 | 0.6466 | 2.3148 | 0.7049 | | 0.1294 |
| *ENO2* | 0.2890 | 0.0457 | 0.0976 | 0.0833 | | 0.0080 |
| *MUM1* | 1.3949 | 0.8561 | 0.4759 | 0.7825 | | 0.3691 |
| *BEND3* | 0.0906 | 0.1107 | 0.6125 | 0.2253 | | 0.0757 |
| *KCNIP2* | 0.0627 | 0.1542 | 0.0286 | 0.0945 | | 0.0177 |
| *PDC* | 4.0970 | 0.2920 | 0.5936 | 0.4637 | | 0.1031 |
| *RUNDC3B* | 0.4267 | 0.3511 | 0.1407 | 0.3266 | | 0.0697 |
| *TMEM136-2* | 0.8141 | 0.3970 | 0.2174 | 0.4389 | | 0.0890 |
| *RHO* | 0.0620 | 0.1328 | 0.1149 | 0.1197 | | 0.0431 |
| *RRH* | 0.4084 | 0.2198 | 0.4350 | 0.3110 | | 0.1312 |
| *CRYAA* | 0.0935 | 0.3236 | 0.1191 | 0.2035 | | 0.0460 |
| *FAM83A* | 0.2657 | 0.2274 | 0.3965 | 0.3106 | | 0.1495 |
| *RLBP1* | 0.1033 | 0.0674 | 0.1733 | 0.1110 | | 0.0351 |
| *ZNF407* | 1.2569 | 0.6139 | 0.5760 | 0.6260 | | 0.3920 |
| *SGCG* | 0.0752 | 0.1392 | 0.2843 | 0.1695 | | 0.0306 |
| *CEP55* | 2.1652 | 1.0990 | 0.4801 | 0.9708 | | 0.4224 |
| *CASR* | 0.0434 | 0.0661 | 0.0999 | 0.0766 | | 0.0289 |
| *FKBP6* | 0.9669 | 0.6302 | 0.4384 | 0.7471 | | 0.3342 |
| *CHST10* | 0.1899 | 0.2257 | 0.4136 | 0.2634 | | 0.0656 |
| *PDLIM3* | 1.1664 | 0.2484 | 0.4213 | 0.3938 | | 0.1494 |
| *ADRA1A* | 0.2044 | 0.1836 | 0.4901 | 0.2776 | | 0.0884 |
| *GPR146* | 0.2949 | 0.2438 | 0.4403 | 0.2995 | | 0.0568 |
| *FHL1* | 0.7873 | 0.1589 | 0.1746 | 0.2654 | | 0.0652 |
| *RP11-463C8.4* | 0.4798 | 0.7896 | 0.6159 | 0.6598 | | 0.2637 |
| *RFXANK* | 1.0069 | 0.3856 | 1.3770 | 0.7037 | | 0.1394 |
| *DRD2* | 0.1987 | 0.1552 | 0.9198 | 0.2854 | | 0.1019 |
| *NPHP3* | 0.1000 | 0.1792 | 0.1727 | 0.1714 | | 0.0884 |
| *AAK1* | 0.1670 | 0.4417 | 0.1812 | 0.2513 | | 0.1322 |
| *PRICKLE1* | 0.1153 | 0.0451 | 0.1724 | 0.0961 | | 0.0375 |
| *KIF26B* | 0.1321 | 0.1685 | 0.2146 | 0.1836 | | 0.1123 |
| *CALHM2* | 0.0371 | 0.0694 | 0.1310 | 0.0838 | | 0.0294 |
| *HSPA2* | 0.0830 | 0.0681 | 0.0162 | 0.0455 | | 0.0131 |
| *BEGAIN* | 0.1063 | 0.2151 | 0.2644 | 0.2021 | | 0.0875 |
| *C3H2orf71* | 1.4416 | 0.6548 | 0.4084 | 0.5798 | | 0.3960 |
| *GAS7* | 0.1300 | 0.2561 | 0.1441 | 0.2105 | | 0.0705 |
| *RPE65* | 0.1248 | 0.0339 | 0.1030 | 0.0692 | | 0.0310 |
| *ARL6* | 0.8677 | 0.2652 | 0.6592 | 0.3914 | | 0.0567 |
| *LRRN3* | 0.3543 | 0.1801 | 0.3305 | 0.2472 | | 0.1234 |
| *TSNAX* | 0.3182 | 0.3185 | 0.2394 | 0.2957 | | 0.0851 |
| *SPNS3* | 1.2045 | 0.5252 | 0.6965 | 0.5962 | | 0.3011 |
| *OPN4-1* | 0.5534 | 0.4355 | 0.4605 | 0.4597 | | 0.2325 |
| *DCLK3* | 0.3268 | 0.3693 | 0.6218 | 0.4242 | | 0.2054 |
| *FBXO21* | 1.3829 | 0.1001 | 0.2019 | 0.1695 | | 0.0516 |
| *LOC428319* | 1.1610 | 0.1940 | 0.7668 | 0.3171 | | 0.1562 |
| *C1H12ORF4* | 0.1800 | 0.1460 | 0.1880 | 0.1647 | | 0.0609 |
| *BEST1* | 0.2125 | 0.1894 | 0.1408 | 0.1806 | | 0.1041 |
| *GSG1L2* | 0.9560 | 0.4434 | 0.5273 | 0.5901 | | 0.2750 |
| *ACER1* | 0.4939 | 0.5089 | 0.2949 | 0.4532 | | 0.1643 |
| *SHANK2* | 0.1780 | 0.0744 | 0.1410 | 0.1268 | | 0.0629 |
| *CCDC51* | 1.4425 | 0.3465 | 0.5441 | 0.4442 | | 0.2130 |
| *TRMT10A* | 0.3208 | 2.2641 | 0.6503 | 0.7142 | | 0.2537 |
| *G6PC2* | 0.1540 | 0.5672 | 0.8092 | 0.4531 | | 0.1321 |
| *LOC107052104* | 0.6353 | 0.6346 | 0.3803 | 0.5589 | | 0.3019 |
| *QSOX2* | 0.2591 | 0.3615 | 0.3344 | 0.3289 | | 0.1679 |
| *TFEC* | 0.3179 | 0.5371 | 0.2269 | 0.3406 | | 0.1057 |
| *GPRIN2* | 1.3676 | 0.7682 | 0.5216 | 0.6787 | | 0.4014 |
| *TOX* | 0.6672 | 0.3162 | 0.2454 | 0.2865 | | 0.0838 |
| *STX18* | 0.9082 | 0.4426 | 0.2132 | 0.3017 | | 0.0945 |
| *HSF3* | 0.7621 | 0.2831 | 0.7257 | 0.3688 | | 0.1769 |
| *INPP5J* | 0.1166 | 0.1781 | 0.2149 | 0.1717 | | 0.0954 |
| *TEKT1* | 0.8477 | 0.9842 | 0.6502 | 0.8270 | | 0.3987 |
| *CSE1L* | 0.0598 | 0.0506 | 0.0378 | 0.0481 | | 0.0153 |
| *MDM4* | 0.7803 | 0.5120 | 0.2855 | 0.4364 | | 0.1861 |
| *MYOT* | 0.7442 | 0.2031 | 0.2157 | 0.2403 | | 0.1066 |
| *NOS2* | 0.8812 | 0.3062 | 0.3670 | 0.3453 | | 0.2163 |

**Table S38. Olfactory receptors identified in 25 avian genomes.** *P*-value was calculated by Mann-Whitney *U* test, after removing two outlier species, chicken and zebra finch.

| Species | Number of intact (functional) OR genes | | | | | Number of  partial OR genes | Number of pseudogenes |
| --- | --- | --- | --- | --- | --- | --- | --- |
|  | Total | α | γ all | γ | γ-c |  |  |
| Eurasian eagle-owl | 56 | 11 | 45 | 27 | 18 | 94 | 104 |
| Northern spotted owl | 88 | 11 | 77 | 30 | 47 | 167 | 154 |
| Oriental scops-owl | 38 | 9 | 29 | 22 | 7 | 172 | 82 |
| Barn owl | 30 | 7 | 23 | 16 | 7 | 67 | 74 |
| Downy woodpecker | 19 | 0 | 19 | 16 | 3 | 36 | 33 |
| Golden eagle | 39 | 9 | 30 | 25 | 5 | 55 | 73 |
| Eastern buzzard | 38 | 9 | 29 | 19 | 10 | 48 | 66 |
| Bald eagle | 40 | 10 | 30 | 20 | 10 | 40 | 66 |
| Zebra finch | 167 | 1 | 166 | 3 | 163 | 284 | 377 |
| American crow | 13 | 4 | 9 | 8 | 1 | 26 | 32 |
| Budgerigar | 26 | 2 | 24 | 23 | 1 | 45 | 38 |
| Saker falcon | 25 | 7 | 18 | 14 | 4 | 24 | 51 |
| Peregrine falcon | 25 | 8 | 17 | 14 | 3 | 25 | 49 |
| Common kestrel | 25 | 10 | 15 | 14 | 1 | 22 | 42 |
| Little egret | 90 | 7 | 83 | 18 | 65 | 120 | 166 |
| Crested ibis | 33 | 5 | 28 | 24 | 4 | 55 | 74 |
| Hoatzin | 53 | 9 | 44 | 24 | 20 | 84 | 126 |
| Killdeer | 49 | 10 | 39 | 32 | 7 | 59 | 86 |
| Chuck-will's-widow | 34 | 6 | 28 | 26 | 2 | 54 | 58 |
| Anna's hummingbird | 23 | 2 | 21 | 17 | 4 | 72 | 85 |
| Common cuckoo | 29 | 7 | 22 | 17 | 5 | 38 | 53 |
| Rock dove | 36 | 7 | 29 | 18 | 11 | 122 | 112 |
| Chicken | 279 | 7 | 272 | 40 | 232 | 426 | 539 |
| Common ostrich | 53 | 22 | 31 | 26 | 5 | 27 | 71 |
| Brown kiwi | 53 | 15 | 38 | 30 | 8 | 63 | 101 |
| *P*-value (nocturnal vs. diurnal) | 0.053 | 0.079 | 0.080 | **0.027** | 0.11 | - | - |
| *P*-value (owls vs. others) | 0.084 | 0.095 | 0.10 | 0.18 | **0.040** | - | - |

**Table S39. The diversity of olfactory receptors in 25 avian genomes.** *P*-value was calculated by Mann-Whitney *U* test.

| Species | Diversity of OR genes by Shannon entropy | | | |
| --- | --- | --- | --- | --- |
|  | α | γ all | γ | γ-c |
| Eurasian eagle-owl | 1.001 | 1.305 | 1.249 | 0.610 |
| Northern spotted owl | 0.978 | 1.076 | 1.265 | 0.478 |
| Oriental scops-owl | 0.895 | 1.178 | 1.142 | 0.432 |
| Barn owl | 0.884 | 1.267 | 1.125 | 0.773 |
| Downy woodpecker | - | 1.169 | 1.148 | 0.174 |
| Golden eagle | 0.862 | 1.247 | 1.218 | 0.317 |
| Eastern buzzard | 0.847 | 1.156 | 1.143 | 0.285 |
| Bald eagle | 0.897 | 1.159 | 1.139 | 0.357 |
| Zebra finch | - | 0.485 | 0.651 | 0.433 |
| American crow | 0.682 | 1.098 | 0.998 | - |
| Budgerigar | 0.350 | 0.989 | 0.947 | - |
| Saker falcon | 0.918 | 1.193 | 1.061 | 0.521 |
| Peregrine falcon | 0.855 | 1.143 | 1.062 | 0.236 |
| Common kestrel | 0.873 | 1.084 | 1.055 | - |
| Little egret | 0.761 | 0.954 | 1.150 | 0.624 |
| Crested ibis | 0.731 | 1.361 | 1.312 | 0.492 |
| Hoatzin | 0.898 | 1.310 | 1.281 | 0.680 |
| Killdeer | 0.965 | 1.307 | 1.277 | 0.448 |
| Chuck-will's-widow | 0.823 | 1.237 | 1.188 | 0.323 |
| Anna's hummingbird | 0.379 | 1.204 | 1.144 | 0.274 |
| Common cuckoo | 0.845 | 1.247 | 1.223 | 0.256 |
| Rock dove | 0.826 | 1.273 | 1.142 | 0.618 |
| Chicken | 0.803 | 0.866 | 1.081 | 0.600 |
| Common ostrich | 1.037 | 1.219 | 1.169 | 0.416 |
| Brown kiwi | 1.045 | 1.267 | 1.151 | 0.620 |
| *P*-value (nocturnal vs. diurnal) | **0.027** | 0.16 | 0.13 | 0.086 |
| *P*-value (owls vs. others) | **0.041** | 0.34 | 0.24 | 0.094 |

**Table S40. Sensory system associated genes showing accelerated *d_N_*/*d_S_* in the nocturnal birds**

| Associated  sensory system | Gene | ω Three nocturnal branches | | | | ω outgroups |
| --- | --- | --- | --- | --- | --- | --- |
|  |  | The ancestral branch of Strigiformes | Chuck-will’s-widow | Brown kiwi | Merged |  |
| Hearing | *ATG5* | 0.1894 | 0.0971 | 0.0854 | 0.1051 | 0.0627 |
| Hearing | *ATP8B1* | 0.1148 | 0.1124 | 0.1165 | 0.1142 | 0.0993 |
| Hearing | *CDH2* | 0.1848 | 0.0898 | 0.0644 | 0.0849 | 0.0416 |
| Hearing | *CEMIP* | 0.1241 | 0.1136 | 0.1311 | 0.1214 | 0.0907 |
| Hearing | *CUX1* | 0.2219 | 0.0996 | 0.0738 | 0.0891 | 0.0677 |
| Hearing | *DFNB59* | 0.0979 | 0.0961 | 0.1630 | 0.1050 | 0.0872 |
| Hearing | *DVL3* | 3.0003 | 0.0398 | 0.0185 | 0.0248 | 0.0183 |
| Hearing | *ENTPD2* | 0.1328 | 0.1406 | 0.1790 | 0.1609 | 0.1169 |
| Hearing | *EYA1* | 0.1798 | 0.1716 | 0.1312 | 0.1740 | 0.0259 |
| Hearing | *FAM65B* | 0.1754 | 0.1673 | 0.1396 | 0.1566 | 0.1225 |
| Hearing | *FGF3* | 0.6052 | 0.1124 | 0.1930 | 0.1730 | 0.1061 |
| Hearing | *FREM2* | 0.1839 | 0.1486 | 0.1488 | 0.1496 | 0.1444 |
| Hearing | *GRXCR1* | 0.9108 | 0.1006 | 0.1889 | 0.1392 | 0.0757 |
| Hearing | *LGR5* | 0.1291 | 0.1884 | 0.2737 | 0.2186 | 0.1274 |
| Hearing | *MBP* | 0.1625 | 0.1769 | 0.2855 | 0.1977 | 0.1532 |
| Hearing | *MCOLN3* | 0.1917 | 0.2126 | 0.1496 | 0.1889 | 0.1478 |
| Hearing | *OPA1* | 0.1441 | 0.0704 | 0.0874 | 0.0784 | 0.0561 |
| Hearing | *OTOA* | 0.6539 | 0.4380 | 0.3596 | 0.4281 | 0.3303 |
| Hearing | *OTOGL* | 0.6029 | 0.3451 | 0.2580 | 0.2942 | 0.2554 |
| Hearing | *PTPRQ* | 0.3033 | 0.2292 | 0.2375 | 0.2354 | 0.1941 |
| Hearing | *RAC1* | 0.1033 | 0.3430 | 0.0275 | 0.2079 | 0.0208 |
| Hearing | *SCN8A* | 0.0291 | 0.0687 | 0.0540 | 0.0567 | 0.0162 |
| Hearing | *SCRIB* | 0.0782 | 0.1259 | 0.1898 | 0.0979 | 0.0761 |
| Hearing | *SLC26A2* | 0.2770 | 0.2236 | 0.2412 | 0.2346 | 0.1955 |
| Hearing | *SLITRK6* | 0.1920 | 0.2145 | 0.1722 | 0.1987 | 0.1465 |
| Hearing | *SUPT6H* | 0.0885 | 0.0242 | 0.0338 | 0.0334 | 0.0187 |
| Hearing | *TMEM132E* | 0.0570 | 0.0430 | 0.0341 | 0.0412 | 0.0282 |
| Hearing | *TTC8* | 0.1442 | 0.1771 | 0.3274 | 0.1974 | 0.1363 |
| Hearing | *USH1G* | 0.2130 | 0.0363 | 0.0614 | 0.0652 | 0.0301 |
| Hearing | *USH2A* | 0.3548 | 0.3061 | 0.3425 | 0.3377 | 0.2257 |
| Hearing | *XPC* | 0.3814 | 0.2982 | 0.5424 | 0.3752 | 0.2890 |
| Hearing/Circadian rhythm | *BDNF* | 0.3643 | 0.1203 | 0.0777 | 0.1429 | 0.0373 |
| Hearing/Circadian rhythm | *DRD2* | 0.1987 | 0.1552 | 0.9198 | 0.2854 | 0.1019 |
| Circadian rhythm | *RPE65* | 0.1248 | 0.0339 | 0.1030 | 0.0692 | 0.0310 |
| Circadian rhythm | *ARNTL2* | 1.3777 | 0.1533 | 0.2053 | 0.2008 | 0.1372 |
| Circadian rhythm | *CASP1* | 0.3965 | 0.6995 | 0.6113 | 0.6194 | 0.3946 |
| Circadian rhythm | *DPYD* | 0.2093 | 0.2351 | 0.2043 | 0.2171 | 0.1772 |
| Circadian rhythm | *FBXL21* | 0.1398 | 0.1204 | 0.2340 | 0.1404 | 0.0931 |
| Circadian rhythm | *GSK3B* | 78.8389 | 0.3286 | 0.3998 | 0.4242 | 0.0608 |
| Circadian rhythm | *KCNH7* | 0.1898 | 0.2387 | 0.1485 | 0.1890 | 0.1152 |
| Circadian rhythm | *MAPK10* | 0.9519 | 0.1575 | 0.1085 | 0.1469 | 0.0507 |
| Circadian rhythm | *NOS2* | 0.8811 | 0.3062 | 0.3670 | 0.3453 | 0.2163 |
| Circadian rhythm | *NR1D2* | 0.1797 | 0.2452 | 0.1391 | 0.1988 | 0.1274 |
| Circadian rhythm | *PHLPP1* | 0.0568 | 0.0580 | 0.1597 | 0.0965 | 0.0525 |
| Circadian rhythm | *PMCH* | 999 | 0.6953 | 0.6341 | 0.6973 | 0.4137 |
| Circadian rhythm | *PPARA* | 0.4024 | 0.0610 | 0.0966 | 0.0858 | 0.0373 |
| Circadian rhythm | *PPARGC1A* | 0.2171 | 0.4580 | 0.2960 | 0.3532 | 0.1827 |
| Circadian rhythm | *PRKG2* | 0.2854 | 0.2165 | 0.3120 | 0.2688 | 0.1993 |
| Circadian rhythm | *SLC6A4* | 0.5796 | 0.1080 | 0.1155 | 0.1328 | 0.0888 |

**Supplementary Methods**

**Genome and transcriptome sequencing**

We sequenced the genomes and transcriptomes from 20 avian species (16 birds of prey and four non-raptor birds) using the Illumina HiSeq platforms (HiSeq2000, HiSeq2500, and HiSeq4000) according to the manufacturer’s sample preparation protocols. The detailed sequencing platform information for each species and data are shown below. To build reference genome assemblies of the four raptor species, we constructed eleven genomic libraries with different insert sizes (170bp, 500bp, 700bp, 2 Kb, 5 Kb, 10 Kb, and 15 Kb for the Eurasian eagle-owl, oriental scops-owl, and eastern buzzard, 350bp, 550bp, 2 Kb, 5 Kb, 10 Kb, and 15 Kb for common kestrel) for each species.

| Species | Common name | Data | Sequencing platform |
| --- | --- | --- | --- |
| *Bubo bubo* | Eurasian eagle-owl | Assembly | HiSeq2500 for short insert libraries,  HiSeq2000 for long-mate pair libraries |
|  |  | Transcriptome | HiSeq2000 |
| *Otus sunia* | Oriental scops-owl | Assembly | HiSeq2500 for short insert libraries, HiSeq2000 for long-mate pair libraries |
|  |  | Transcriptome | HiSeq2500 |
| *Strix nivicolum* | Himalayan owl | Resequencing | HiSeq2000, HiSeq2500 |
|  |  | Transcriptome | HiSeq2000 |
| *Ninox japonica* | Northern boobook | Resequencing | HiSeq2000, HiSeq2500 |
|  |  | Transcriptome | HiSeq2000 |
| *Asio otus* | Long-eared owl | Resequencing | HiSeq2000, HiSeq2500 |
|  |  | Transcriptome | HiSeq2000 |
| *Asio flammeus* | Short-eared owl | Resequencing | HiSeq4000 |
| *Otus semitorques* | Japanese scops-owl | Resequencing | HiSeq4000 |
| *Buteo japonicus* | Eastern buzzard | Assembly | HiSeq2500 for short insert libraries, HiSeq2000 for long-mate pair libraries |
|  |  | Transcriptome | HiSeq2500 |
| *Accipiter nisus* | Eurasian sparrowhawk | Resequencing | HiSeq2500 |
|  |  | Transcriptome | HiSeq2500 |
| *Accipiter gentilis* | Northern goshawk | Resequencing | HiSeq2500 |
|  |  | Transcriptome | HiSeq2500 |
| *Haliaeetus albicilla* | White-tailed eagle | Resequencing | HiSeq4000 |
| *Pernis ptilorhynchus* | Oriental honey-buzzard | Resequencing | HiSeq4000 |
| *Milvus migrans* | Black kite | Resequencing | HiSeq4000 |
| *Accipiter soloensis* | Chinese sparrowhawk | Resequencing | HiSeq4000 |
| *Falco tinnunculus* | Common kestrel | Assembly | HiSeq2500 for shot-insert  and long-mate pair libraries |
|  |  | Resequencing | HiSeq2500 |
|  |  | Transcriptome | HiSeq2500 |
| *Falco subbuteo* | Eurasian hobby | Resequencing | HiSeq2500 |
|  |  | Transcriptome | HiSeq2500 |
| *Picus canus* | Grey-headed woodpecker | Resequencing, | HiSeq2000 |
|  |  | Transcriptome | HiSeq2000 |
| *Egretta garzetta* | Little egret | Resequencing | HiSeq2000 |
|  |  | Transcriptome | HiSeq2500 |
| *Butorides striata* | Striated heron | Resequencing | HiSeq2000 |
|  |  | Transcriptome | HiSeq2500 |
| *Platalea minor* | Black-faced spoonbill | Resequencing | HiSeq2000 |
|  |  | Transcriptome | HiSeq2000 |

**Species identification**

Species of the sequenced samples were confirmed by mapping their DNA sequences to previously reported mitochondrial sequences (*COI* and *CYTB* genes) for closely related species using BWA-MEM [18] with default options. Variants were identified using the mpileup command in SAMtools [19]. The consensus sequences were generated using the vcf2fq command. The *COI* gene of common kestrel was sequenced by Sanger method. Phylogenetic reconstruction was performed using MrBayes 3.2 software [20] with the “lset=mixed rates=invgamma” substitution model specifications. Species sampling in the phylogeny was designed to include all species from the families Accipitridae, Ardeidae, Falconidae, Picidae, Strigidae, and Threskiornithidae that occur in South Korea. Species that could not be included were *Aviceda leuphotes* (Accipitridae), *Dendrocopos hyperythrus* (Picidae), and *Threskiornis melanocephalus* (Threskiornithidae). In case of the two latter species, congeneric species were included. KM364882 sequence was first attributed to *B. buteo burmanicus*, a junior synonym of *B. refectus*; the sampling locality is outside the known range of *B. refectus* and suggests that it is a misidentified *B. japonicus*. The latter hypothesis is also supported by comparative analyses with the tRNAGlu-Pseudo-control Region sequences [21].

**Sequence filtering criteria**

To reduce sequencing error effects in assembling the bird of prey genomes (Eurasian eagle-owl, oriental scops-owl, eastern buzzard, and common kestrel), we filtered out PCR duplicated, low quality, and adaptor contaminated reads. The filtering criteria for exclusion were as follows:

1) Reads were considered PCR duplicates if read1 (left) and read2 (right) of the two paired end reads were identical. The PCR duplicated reads were filtered out remaining one unique read pairs.
2) Reads with sequencing adapter contamination were filtered out.

Sequencing adapter left= "*GATCGGAAGAGCACACGTCTGAACTCCAGTCAC*"

Sequencing adapter right= "*GATCGGAAGAGCGTCGTGTAGGGAAAGAGTGT*"

3) Reads with ambiguous base (N) for more than 5% of the reads were filtered out.
4) Reads with an average base quality below 20 (<Q20) were filtered out.
5) Reads with junction adapter contamination for mate-pair libraries were filtered out.

Junction adapter left = "*CTGTCTCTTATACACATCT*"

Junction adapter right = "*AGATGTGTATAAGAGACAG*"

6) To filter out low-quality read ends, three bases of 5’-end and eight bases of 3’-end of each read from short insert libraries were trimmed.
7) Each read from long-mate pair libraries were trimmed into 50bp (one base of 3’-end of each read for Eurasian eagle-owl, oriental scops-owl, and eastern buzzard; 51 bases of 3’-end of each read for common kestrel)

**Repeat annotation**

For the annotation of repetitive elements for the assembled bird of prey genomes, we searched the bird of prey genomes for tandem repeats using the Tandem Repeats Finder (version 4.07b) [22]. Transposable elements (TEs) were identified in the genomes by homology-based and *ab initio*-based approaches. For the homology-based approach, we identified repeats using Repbase (version 19.03) [23] with RepeatMasker (version 4.0.5) [24] and RMBlast (version 2.2.28) [25]. For the *ab initio*-based approach, we used RepeatModeler (version 1.0.7) [26]. All predicted repetitive elements were merged for statistics by in-house scripts. Roughly ~9.2 % of the bird of prey genomes were predicted as transposable elements, which are similar in composition to the other avian genomes [2].
